# Reducing haystacks to needles – ViralClust: A Nextflow pipeline to cluster viral sequences

**DOI:** 10.64898/2026.01.30.702815

**Authors:** Sandra Triebel, Kevin Lamkiewicz, Tom Eulenfeld, Manja Marz

**Affiliations:** RNA Bioinformatics and High-Throughput Analysis, Friedrich Schiller University Jena, Jena, 07743, Germany; European Virus Bioinformatics Center, Friedrich Schiller University Jena, Jena, 07743, Germany

**Keywords:** Virus, Clustering, Nextflow, Bioinformatics, Viral Evolution

## Abstract

**Background:** The rapid accumulation of viral genome sequences presents major challenges for downstream analysis tools, including tools for multiple sequence alignments, phylogeny, and genome/alignment visualization, due to computational constraints and sampling biases caused by outbreak-driven over-representation. Selecting representative genomes through clustering offers a principled alternative to random subsampling, yet choosing appropriate clustering strategies remains non-trivial and context dependent.

**Results:** Here, we present `ViralClust`, a modular `Nextflow` pipeline for bias-aware representative selection from large viral genome datasets. `ViralClust` integrates five distinct clustering algorithms (`CD-HIT-EST`, `SUMACLUST`, `VSEARCH`, `MMSeqs2`, and `HDBSCAN`) within a unified workflow, enabling direct comparison of clustering outcomes and flexible adaptation to diverse biological questions, considering a balanced phylogenic distribution of the selected sequences. We evaluated `ViralClust` on six RNA and DNA virus datasets ranging from 632 to 156,586 sequences and spanning genome lengths from 890 to 197,185 nucleotides. Across all datasets, clustering reduced dataset size by ∼95 % or more while preserving genetic diversity across species, genera, and families, and effectively mitigating biases introduced by outbreaks, partial genomes, and sequence orientation artifacts.

**Conclusions:** By supporting whole-genome clustering and scalable representative selection, `ViralClust` enables efficient and reproducible downstream analyses that would otherwise be computationally infeasible. Our framework provides a flexible foundation for large-scale viral genomics and supports future applications in comparative analysis and virus classification.

**Key Points:**

- `ViralClust` is a modular `Nextflow` pipeline for selecting representative viral genomes from (very) large sequence datasets.
- It combines multiple clustering approaches to reduce dataset size while minimizing bias and preserving genetic diversity.
- Tests on six RNA and DNA virus datasets show reductions of ∼95 % across a wide range of genome sizes and sequence counts.
- `ViralClust` enables efficient and reproducible downstream analyses that are otherwise impractical with full viral genome collections.

## Background

Viral genome sequences are becoming increasingly available [1, 2, 3, 4, 5]. For example, for SARS-CoV-2 alone, nearly 9.2 million uploaded nucleotide sequences are accessible (NCBI Virus, 19.11.2025). The use of such large datasets of viral sequences often poses problems for downstream analyses, such as sequence alignment, phylogeny, and visualization, in terms of memory usage, run time, and visualization [6, 7]. Calculating a multiple sequence alignment (MSA) of available viral genomes of a family is often computationally not feasible. Disregarding the technical demands, an MSA with tens of thousands of entries may not be informative for most users, since the overall alignment quality potentially decreases. black-Several strategies have been proposed to handle large-scale viral datasets. Reference-based alignment approaches map sequences against a single reference genome, reducing computational over-head but potentially missing diversity in divergent strains [8, 9, 10]. Similarly, k-mer-based methods enable rapid, alignment-free comparison of sequences and have been widely used for taxonomic classification and similarity estimation [11, 12, 13]. However, both approaches are primarily designed for sequence comparison rather than dataset reduction, and neither directly addresses the problem of selecting a representative, diversity-preserving subset of genomes for downstream analysis. Besides the size of the datasets, the over-representation of a certain species due to an outbreak might also introduce a bias into the data. To reduce the number of entries, random sampling of genomes does not solve the introduced bias of over-represented species it may even increase the effect of the bias. Consequently, the aim is to reduce the dataset of viral sequences without losing genetic diversity. blackIn this context, cluster-based representative sequence selection offers a distinct advantage: by grouping sequences according to their similarity and retaining one representative per cluster, the resulting subset reflects the full breadth of genetic diversity present in the original dataset, regardless of sampling biases introduced by outbreaks or sequencing efforts. Instead of random selection, clustering viral genomes based on, e.g., sequence similarity and selecting a representative genome for each cluster results in a better-balanced subset of input viruses. While the idea of clustering is simple, the correct method is often hard to find and depends on the underlying scientific question. Depending on the biological question and distribution of biological data, the principle of *compactness* is most widely used, aiming to keep intra-cluster variation small. It is implemented in tools such as `CD-HIT-EST` [14], `SUMACLUST` [15], `VSEARCH` [16], and `MMSeqs2` [17, 18]. In this context, the sequence identity determines whether a sequence belongs to a cluster or forms a new, independent cluster. Thus, compactness clustering is based on sequence homology. In most implementations, a sequence is only compared against one representative of established clusters. This method is used frequently for viral clades [19, 20, 21]. However, evolution is a continuous process, and many novel observed viruses might not be unambiguously assigned to the calculated clusters based on static sequence similarity thresholds alone. An algorithm following the principle of *connectedness* may be suitable for such continuous data. The principle of connectedness exploits the idea that neighboring data points should belong to the same cluster. Therefore, algorithms following this principle tend to detect smaller, local components within the data that are not concentrated around a center point [22]. For datasets with viral genomes such that the data points are not well-described by high-dimensional spheres, as is the case for compactness clusters, we propose a tool such as `HDBSCAN` [23]. Virologists, mostly lacking computational backgrounds, often find themselves unable to navigate the intricacies of algorithmic advantages and disadvantages, leaving them uncertain about selecting the most suitable tool for their analyses. For bioinformaticians, evaluating which tool to choose presents its own challenges, as the absence of a definitive ground truth complicates the determination of which sequences best represent a cluster.

Here, we present `ViralClust`, a `Nextflow` pipeline designed to select representative viral genomes, by a streamline clustering analysis for virology research. Unlike conventional methods, `ViralClust` integrates clustering via five distinct existing tools, enabling direct and comprehensive comparison of the results. One key feature of `ViralClust` is its robust data pre-processing capabilities. We address common database errors, such as inconsistencies in sequence orientation (-ssRNA sequences found in both positive and negative orientations) and ambiguous gene arrangements in ambisense viruses. By correcting these mistakes, we enhance the accuracy of clustering outcomes. Moreover, `ViralClust` introduces innovative methodologies previously unexplored in virology research. For instance, full genome clustering based on k-mers of nucleotide sequences utilizing Hierarchical Density-Based Clustering (HDB) represents a novel approach in virus research, offering previously inaccessible insights. Finally, we provide comprehensive statistics, including cluster sizes, and visualize the sequence selection in rough phylogenetic representations without explicitly generating phylogenies of the viral families. Additionally, our HTML output facilitates easy cluster sorting, including direct links to corresponding NCBI genomes for further analysis. All parts of the developed pipeline are written modular, and the user may adjust parameters or exchange and add complete tools. `ViralClust` offers the possibility of adding manually additional viral **g**enomes **o**f **i**nterest (GOI) to the input data. We evaluated the performance of `ViralClust` on six different datasets of RNA and DNA viruses varying in number of available sequences, sequence length, and sequence diversity, such as segmentation and orientation.

## Data Description

### Viral data

We downloaded viral genomes labeled as `‘complete’` of *Orthoebolavirus, Hepacivirus hominis* (HCV), *Orthoflavivirus, Flaviviridae, Alphainfluenzavirus influenzae* (formerly *Influenza A virus*, IAV), and *Monkeypox virus* from the NCBI Virus database, see Tab. 1. All sequences used in this study are deposited in the zenodo repository (Link: https://doi.org/10.5281/zenodo.18413373) in `fasta` format.

**Table 1.** Overview of datasets used in this study. `ViralClust` reduces the number of input genomes to a representative set of genomes. Data was downloaded from NCBI Virus. IAV – *Alphainfluenzavirus influenzae*; ∅ SeqLen – average sequence length; #Seq – number of input genomes / segments; contains only records labeled as “`complete`”; PreProc – number of sequences after pre-processing; %nr – percentage of non-redundant sequences; Number of genomes after clustering – only considering clusters with a size greater than one; cd – `CD-HIT-EST`; sum – `SUMACLUST`; vse – `VSEARCH`; mmseq – `MMSeqs2`; hdb – `HDBSCAN`; Time – run time of `ViralClust` measured by `Nextflow`; blackPeak RAM – peak RSS (resident set size) of `ViralClust` measured by `Nextflow`. blackA detailed overview of time and memory consumption is shown in supplementary Tab. S1.

| Taxonomic clade | ∅ SeqLen | #Seq | PreProc | %nr | Number of genomes after clustering | | | | | Time | Peak RAM |
| --- | --- | --- | --- | --- | --- | --- | --- | --- | --- | --- | --- |
|  |  |  |  |  | cd | suma | vse | mmseq | hdb |  |  |
| *Orthoebolavirus* | 18.9 kb | 632 | 443 | 70.09 | 8 | 7 | 7 | 11 | 11 | 12m | 11.4 G |
| *Hepacivirus hominis* | 9.6 kb | 1,075 | 1,071 | 99.63 | 55 | 30 | 58 | 55 | 23 | 14m | 10.7 G |
| *Orthoflavivirus* | 9.2–11 kb | 8,052 | 7,681 | 95.39 | 69 | 71 | 70 | 63 | 101 | 1h 22m | 43.2 G |
| *Flaviviridae* | 9–13 kb | 9,895 | 9,478 | 95.79 | 220 | 180 | 221 | 211 | 194 | 2h 47m | 52.6 G |
| IAV (1) | 2,341 nt | 83,543 | 56,973 | 68.20 | 66 | 390 | 90 | 280 | 1,412 | 3h 27m | 273.9 G |
| IAV (2) | 2,341 nt | 82,569 | 56,212 | 68.07 | 56 | 338 | 73 | 216 | 1,311 | 3h 24m | 270.9 G |
| IAV (3) | 2,233 nt | 83,400 | 55,862 | 66.98 | 63 | 305 | 75 | 265 | 1,371 | 3h 16m | 268.8 G |
| IAV (4, H) | 1,775 nt | 156,586 | 108,514 | 69.30 | 300 | 1,237 | 464 | 1,574 | 2,307 | 8h 36m | 510.0 G |
| IAV (5) | 1,565 nt | 84,653 | 49,465 | 58.43 | 46 | 262 | 76 | 179 | 1,218 | 2h 28m | 238.3 G |
| IAV (6, N) | 1,409 nt | 114,639 | 73,786 | 64.36 | 228 | 773 | 329 | 640 | 1,622 | 4h 9m | 572.2 G |
| IAV (7) | 1,027 nt | 96,928 | 46,371 | 47.84 | 25 | 185 | 38 | 269 | 981 | 1h 47m | 225.1 G |
| IAV (8) | 890 nt | 85,944 | 42,308 | 49.23 | 51 | 218 | 69 | 264 | 802 | 1h 28m | 207.2 G |
| *Monkeypox virus* | 190–200 kb | 1,077 | 999 | 92.76 | 14 | 1 | 3 | 23 | 2 | 16h 0m | 6.5 G |

### ‘Ground Truth’ of Virus Phylogeny

To compare the results of `ViralClust`, we considered the classification of the ICTV [24, 25] as ‘ground truth’ for *Orthoebolavirus, Orthoflavivirus, Flaviviridae*. For *Hepacivirus hominis* and *Monkeypox virus* we included the NCBI taxonomy. The number of genera, subgenera and/or species is derived from https://ictv.global/taxonomy, see Tab. S2. For *Alphainfluenzavirus influenzae* we included the classification according to the Centers for Disease Control and Prevention (CDC)^1^ into 18 different H subtypes and 11 different N subtypes. This theoretically results in 198 possible subtypes, but 136 subtypes are currently listed in the NCBI taxonomy (23.01.2026).

## Analyses

### The ViralClust pipeline

Our pipeline `ViralClust` takes a multiple `fasta` file as input and applies five different cluster tools to the data to select representative genomes. We calculate and report basic statistics for each of the cluster tools. Based on the respective representative genomes, `ViralClust` further calculates a multiple sequence alignment (MSA) and a rough phylogenetic representation (parameter `--eval`). Fig. 1 gives a general overview of the pipeline. Additionally to the clustering step, the user can retrieve an optional basic evaluation of the results. This evaluation is no recommendation for any algorithm or tool, as the choice of clustering depends on the underlying scientific question. However, it gives a broad overview of the data and may help determine the appropriate cluster method. If applicable, `ViralClust` finds GenBank accession IDs in the `fasta` records identifier and retrieves meta information from the NCBI database. Lastly, the user can specify a second, optional `fasta` file containing genomes that must be present in the final set of representative genomes. This may prove useful if comparative analyses between viruses are carried out and the user wants to include a specific, unpublished viral strain or isolate in the analysis.

**Figure 1.**
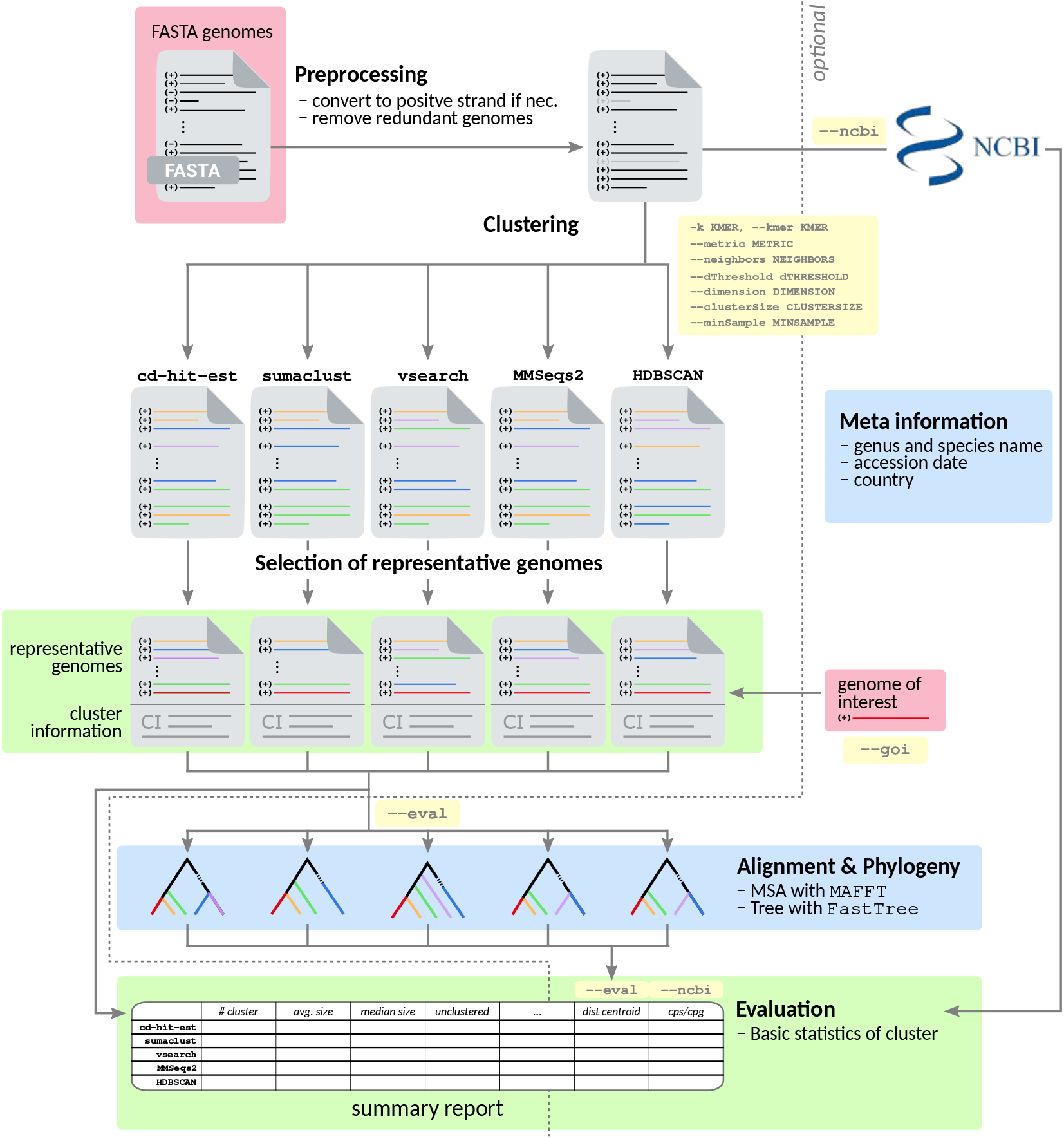
The general `ViralClust` pipeline. *Pre-processing*. Input genomes are filtered for redundant genomes and oriented to the positive strand determined via an ORF density scan. *Clustering*. Five different tools are pre-installed. However, each can be exchanged by other customized clustering tools. For each clustering method applied, `ViralClust` provides an overview of which sequence falls into which cluster. *Representative genomes*. Further, a `fasta` file with representative sequences for each cluster is provided for each tool. `--goi` – Genomes of interest (GOI) or unpublished genomes can be added to the representative genomes. *Evaluation*. The basic statistics of the cluster results, including average and median cluster size and the average distance of two representative genomes are given for each cluster tool. *Optional parameters*. `--ncbi` – For each applicable sequence, *i*.*e*. for each `fasta` record with a valid GenBank accession ID in its identifier, `ViralClust` retrieves additional information from the NCBI database and includes it in the summary report. `--eval` – For each clustering method `ViralClust` can provide an alignment of representative genomes and a phylogenetic representation of these genomes.

#### Workflow of ViralClust

We implemented `ViralClust` in `Nextflow` (v22.10.06) [26]. We developed our scripts with `Python` (compatible with v3.6+). We employ individual `conda` (v4.9.2) [27] environments for each step per default to guarantee no conflicts during installation and usage. `ViralClust` is publicly available at https://github.com/rnajena/viralclust.

We further provide a test dataset with a small tutorial on using `ViralClust`. In the following, we describe the pipeline depicted in Fig. 1 in detail, including implemented tools and parameter configurations.

##### Pre-processing

blackTo ensure high-quality data for subsequent clustering, `ViralClust` discards sequences with more than 10 % of non-`ACGU` characters. The user can adjust this filtering process using the `--max_ambiguous` parameter. Both strands are then scanned for ORF density to ensure that all sequences have the same orientation. Especially in the case of single-stranded negative RNA viruses, genomes are sometimes uploaded into the database in different orientations. This causes cluster algorithms to separate genomes based on this orientation, potentially jeopardizing the complete clustering approach. We calculate ORF density by translating an input genome into all six reading frames. We define an open reading frame (ORF) as a sequence of at least 200 amino acids between two stop codons or between a single stop codon and the 5’ or 3’ end of the sequence. This makes the procedure robust against missing stop codons at the end of the sequence and missing start codons. We calculate the forward-strand ORF density by summing the lengths of the ORFs in the three forward reading frames, then dividing by the sequence length. The backward-strand ORF density is calculated analogously. This method accounts for real or erroneous frameshifts. We use the strand with the higher ORF density for further calculations. The step is enabled by default; the user can disable it using the parameter `sort_off`. To remove 100 % redundant genomes in the input dataset, we apply the `easy-linclust` module of `MMSeqs2` (v14) [17, 18], with parameters `--min-seq-id 1.0`.

##### Clustering

Per default, `ViralClust` applies five different clustering methods. Depending on which scientific virological question is asked, one of the tools might be more suitable than another. Due to the modularity of the pipeline, the user can also choose to switch off the predefined cluster tools and/or add individual cluster methods. Currently, we implemented `CD-HIT-EST` (v4.8.1) [14], `SUMACLUST` (v1.0.31) [15], `VSEARCH` (v2.15) [16] from `vsearch`, `MMSeqs2` (v14) [17, 18], and `HDBSCAN` (v0.8.26) [23] as individual `Nextflow` modules. Whenever applicable, we set the parameters of the tools to match each other to make them comparable. By default, `ViralClust` instructs `CD-HIT-EST`, `SUMACLUST`, `VSEARCH`, and `MMSeqs2` to cluster sequences with an identity of at least 90 % (default value of `CD-HIT-EST`). blackSince `MMSeqs2` offers a variety of clustering modules, we have integrated `easy-linclust` (default) and `easy-cluster`.

##### *Connectedness Clustering with* *HDBSCAN*

As a novel approach to cluster viral genomes, we further implemented a combination of `PCA` and `HDBSCAN` [23] in `ViralClust`. *(1) k-mer distributions:* For this approach, we translate each input genome into a vector of k-mer frequencies (default *k* = 7), neglecting k-mers consisting of non-canonical nucleotides. *(2) Dimension reduction:* For the `PCA`, we use the module as implemented in `scikit-learn` (v1.2.2) [28]. Instead of the dimension reduction with `PCA`, we further allow using `UMAP` for the reduction. *(3) Clustering:* For efficient clustering, we reduce the resulting 4^*k*^-dimensional (default *k* = 7) vector space and thus provide an embedding for the clustering algorithm `HDBSCAN`. Per default, we either reduce the dimension to *d* = 50 or until the sum of explained variance of the components is at least 0.7. After dimension reduction, each viral genome is represented by a vector of maximal length *d* (*d* = 50). `HDBSCAN` can be seen as a hybrid approach of hierarchical-based (connectivity) and density-based (compactness) clustering approaches. For two vectors, `HDBSCAN` first calculates mutual reachability distance. Per default, we use the cosine distance implemented in `SciPy` (v1.4.1) [29] as metric. However, we allow for a plethora of pre-defined distances, which can be used via the `--metric` parameter of our `HDBSCAN` module. We implemented the (standardized) Euclidean, Manhattan, Chebyshev, Minkowski, Canberra, Braycurtis, and Mahalanobis distance metrics via the `SciPy` library [29]. Since the used implementation of `HDBSCAN` cannot deal with the cosine distance, we normalized the vectors using the L2 normalization. Subsequently, we use `HDBSCAN` with the Euclidean distance on these normalized vectors^2^.

The user can override or provide additional parameters for each tool via the `nextflow run viralclust.nf` command. Exemplarily, for `HDBSCAN`, the user can use the flag `--hdbscan_help` to get an overview of implemented options, which can be used using `--hdbscan_params`. Analogously, help messages and parameters for the other tools can be used and accessed via their respective commands.

##### Determination of representative genomes

The implemented tools report representative genomes based on sequence length (`CD-HIT-EST`, `SUMACLUST`), sequence abundance (`VSEARCH`), and selection within cascaded clustering (`MMSeqs2`). For the `HDBSCAN` module, we calculate centroid sequences based on the average pairwise distance between all vectors in the same cluster. The sequence with the minimum average distance is selected as the representative genome for this cluster. Given a distance metric *d*, the centroid sequence 𝒸 of a cluster *C* with size *N* is the one that has the lowest average distance 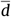 to all other sequences *S*_*i*_ (*i* ∈ [1, *N*]) of the cluster: 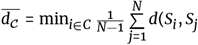. Thus, each cluster is represented by an input genome, not an ‘artificial’ median genome. Per default, we use the cosine distance as metric *d* but also allow for other metrics. In `ViralClust`, only clusters with a size of more than one are defined as ‘real’ clusters and the corresponding representatives are saved in a file. blackClusters of size one are therefore not considered clusters, but named ‘singletons’ here, and are stored in a separate file. The combined set of cluster representatives and ‘singleton’ sequences is provided in a third file. This allows the user to choose which set of sequences is most suitable for the research question.

##### Optional parameters

*(1) Inclusion of a genome of interest:* In addition to the general input file, the user can specify a second `fasta` file with the `--goi` flag. This `fasta` file should contain one (or more) **g**enome(s) **o**f **i**nterest (GOI). If the user provides such a file, `ViralClust` will include the contained genomes in the final set of representative genomes. Further, in the result reports of the individual cluster tools, the GOIs are specifically marked. With this option, users can add their unpublished in-lab strain or mutant to downstream analyses and can further determine the closest phylogenetic viruses based on the resulting clusters. This also enables comparisons between the representative sequences and specific reference genomes, which are not necessarily cluster representatives. *(2) Metadata Information:* With the `--ncbi` flag `ViralClust` links additional information provided by the NCBI [30]. For this, we can sequence identifiers for GenBank accession ID. If found, we use the accession ID to retrieve additional information about the taxonomy (species and genus name), accession date, and the country the sequence originated from. `ViralClust` combines this metadata with the overview of clusters for each tool. *(3) Alignment and Phylogeny:* If the user sets the flag `--eval` of `ViralClust`, we further create a multiple sequence alignment (default: `MAFFT` (v7.4) [31]) of the representative genomes for each cluster tool. We subsequently use the alignments to build phylogenetic representations (default: `FastTree` (v2.1.10) [32]), which we finally visualize using `Newick Utilities` (v1.6) [33]. If `--eval` is enabled, we include the average phylogenetic distance calculated by `FastTree` between two representative genomes in the final summary report. Per default, `FastTree` uses the Jukes-Cantor distance model [34].

### Benchmark ViralClust by a variety of datasets

We applied `ViralClust` to six viral datasets of single-stranded RNA and double-stranded DNA viruses varying in number of sequences, sequence length, and sequence diversity. For an overview of the datasets used in this study, we refer to Tab. 1, in which we provide the total number of available sequences, the number of sequences after the implemented filtering steps, and the number of clusters resulting from each tool. Given these input sets, we report the number of clusters as calculated by the individual tools in their respective column, run time, and memory usage, see Tab. 1 and S1. All results generated by `ViralClust` are deposited in the zenodo repository: https://doi.org/10.5281/zenodo.18413373.

blackTo provide an objective basis for comparing tool performance across datasets, we evaluated all five clustering algorithms using a scoring framework comprising five component scores: the Reduction Score (RS), Singleton Penalty Score (SPS), Clustering Quality Score (CQS), Taxonomic Concordance Score (TCS), and Cluster-to-Taxonomy Ratio (CTR), aggregated into an Overall Score (OS) with a maximum of 5. A detailed description of each metric is provided in the Methods section. The resulting scores are summarized in Tab. 2. All scores are shown in Tab. S3.

**Table 2.** blackOverall Score (OS) of the five clustering algorithms integrated into `ViralClust` across all evaluated datasets. The OS is the sum of five component scores - Reduction Score (RS), Singleton Penalty Score (SPS), Clustering Quality Score (CQS), Taxonomic Concordance Score (TCS), and Cluster-to-Taxonomy Ratio (CTR) — each in the range [0, 1], giving a maximum OS of 5. Higher values indicate better overall clustering performance with respect to dataset reduction, singleton rate, phylogenetic cluster separation, and taxonomic concordance. IAV – *Alphainfluenzavirus influenzae*; cd – `CD-HIT-EST`; sum – `SUMACLUST`; vse – `VSEARCH`; mmseq – `MMSeqs2`; hdb – `HDBSCAN`. All scores are shown in Tab. S3.

| Taxonomic clade | Overall Score |  |  |  |  |
| --- | --- | --- | --- | --- | --- |
|  | cd | suma | vse | mmseq | hdb |
| *Orthoebolavirus* | 4.622 | 4.682 | 4.687 | 3.940 | 3.720 |
| *Hepacivirus hominis* | 3.926 | 3.798 | 4.040 | 3.963 | 3.998 |
| *Orthoflavivirus* | 4.331 | 4.141 | 4.340 | 4.508 | 3.294 |
| *Flaviviridae* | 3.516 | 3.523 | 3.540 | 3.555 | 3.022 |
| IAV (1) | 4.422 | 4.852 | 4.930 | 4.424 | 4.384 |
| IAV (2) | 4.446 | 4.836 | 4.909 | 4.401 | 4.414 |
| IAV (3) | 4.396 | 4.829 | 4.911 | 4.502 | 4.365 |
| IAV (4, H) | 4.655 | 4.838 | 4.938 | 4.561 | 4.474 |
| IAV (5) | 4.385 | 4.871 | 4.925 | 4.415 | 4.239 |
| IAV (6, N) | 4.612 | 4.859 | 4.929 | 4.571 | 4.385 |
| IAV (7) | 4.545 | 4.687 | 4.785 | 4.693 | 4.131 |
| IAV (8) | 4.605 | 4.808 | 4.873 | 4.665 | 4.247 |
| *Monkeypox virus* | 2.690 | 2.311 | 4.161 | 2.763 | 4.996 |
| Sum of Scores | 55.151 | 57.035 | 59.968 | 54.961 | 53.669 |

blackThe scores reveal a considerable spread both across tools and across datasets, with OS values ranging from 2.311 (`SUMACLUST`, *Monkeypox virus*) to 4.996 (`HDBSCAN`, *Monkeypox virus*), reflecting the diversity of the evaluated datasets rather than a consistent ranking of tools. No single tool dominates across all datasets, and tool choice should therefore be guided by the biological question and dataset composition. Several general trends emerge from the scores and are discussed in detail in the respective subsections below. Compactness-based tools (`CD-HIT-EST`, `SUMACLUST`, `VSEARCH`) tend to score consistently well across RNA virus datasets, producing fewer, more compact clusters that achieve strong dataset reduction, but may collapse rare or biologically meaningful lineages in taxonomically diverse datasets. Noteworthy, `SUMACLUST` discards sequences containing characters other than `ACGT`, making it even more dependent on the quality of the input data and possibly difficult to use the results as a representative genomic dataset for subsequent analyses. `MMSeqs2` produces higher cluster granularity, better preserving sequence-level diversity at the cost of larger representative sets, though its behavior becomes less predictable at very large dataset sizes. `HDBSCAN` is particularly well suited for datasets with continuous diversity distributions, but carries a fundamental limitation for datasets containing rare subtypes, which may not form sufficiently dense regions to be recognized as independent clusters and are instead treated as noise. These observations are intended as a starting point for tool selection; the scoring framework provided by `ViralClust` allows users to reproduce the same evaluation on their own datasets.

#### Small and biased dataset - Orthoebolavirus

We tested the *Orthoebolavirus* dataset consisting of only 632 genomes to see how fine-grained `ViralClust` can work. *Ortho-ebolavirus* is a genus within the *Filoviridae* family (ICTV phylogeny see Fig. S1). The genomes are negative single-stranded RNA and about 18.9 kb in length [35]. In comparison to other viral clades, this dataset includes a small number (six) of well-described species according to the ICTV, see Tab. 3, with a notable presence with 461/632 sequences of *Orthoebolavirus zairense* (Ebola virus, EBOV), a human pathogen over-represented due to several outbreaks. We aim to investigate the selection of representative genomes with `ViralClust` for such a small, well-defined, and unbalanced dataset. The pre-processing of `ViralClust` reduced the dataset size by 30 % to 443 genomes. Subsequent clustering resulted in 7 to 11 representative genomes, see Tab. 1. In general, all five clustering tools could distinguish the species in individual clusters, see Fig. 2. A conventional representation of the phylogeny of the representative genomes and the corresponding ICTV classification can be found in Fig. S2. Additionally, in Fig. S3 a `RAxML-NG` phylogenetic tree based on a full-genome `MAFFT` alignment can be found as an overview for the 443 individual sequences. `CD-HIT-EST`, `SUMACLUST`, `VSEARCH` and `MMSeqs2` revealed correctly one cluster representative for the five species BOMV, BDBV, RESTV, SUDV, and TAFV each. A notable exception is `HDBSCAN`, which could not identify the species TAFV, Fig. 2B and Fig. S2F. All tools clustered TAFV genomes (light blue) correctly together with BDBV (dark green). Each of the tools selected at least one natural EBOV strains (`CD-HIT-EST`:1, `SUMACLUST`:1, `VSEARCH`:1, `MMSeqs2`:4, `HDBSCAN`:7). Additionally, `CD-HIT-EST`, `SUMACLUST`, `VSEARCH` and `MMSeqs2` selected up to two representatives of EBOV derived from recombination or reverse genetics experiments (see Fig. 2 dashed gray box). `HDBSCAN` selected only natural EBOV sequences (isolated from viral infections). Interestingly, these comparatively high number of representative genomes represent geographic and temporal separation of EBOV outbreaks.

**Figure 2.**
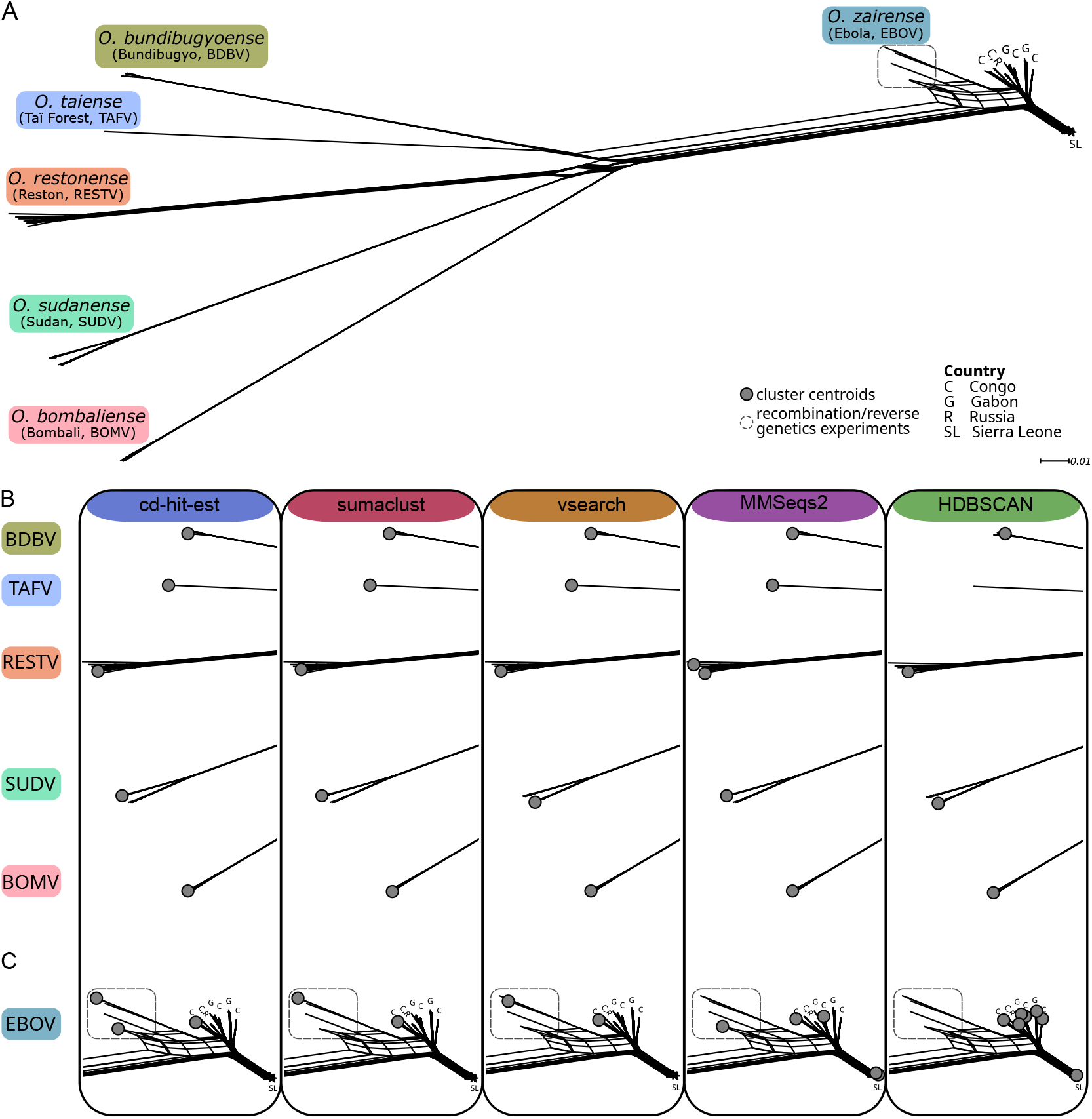
(A) Split graph of the *Orthoebolavirus* dataset with labeled species. (B) Cluster representatives of `CD-HIT-EST`, `SUMACLUST`, `VSEARCH`, `MMSeqs2`, and `HDBSCAN` in the subtrees of BDBV, TAFV, RESTV, SUDV, and BOMV (top to bottom). (C) Cluster representatives in the EBOV subtree. Gray dots display the cluster representative sequences selected by the individual tools. The Zaire *Orthoebolavirus* subtree shows a distinction by geographic origin, indicated as follows: C – Congo; G – Gabon; R – Russia; SL – Sierra Leone. The split graph is reconstructed using `SplitsTree` based on the `MAFFT` alignment of the complete genome sequences (443 genomes) resulting from the first filtering steps of `ViralClust`.

**Table 3.** *Orthoebolavirus* species including virus name and abbreviation (Abbr.) according to ICTV and number of downloaded sequences per species (#Seqs). 93 sequences were unclassified regarding their species.

| Species | Virus name | Abbr. | #Seqs |
| --- | --- | --- | --- |
| *O. bombaliense* | Bombali | BOMV | 8 |
| *O. bundibugyoense* | Bundibugyo | BDBV | 24 |
| *O. restonense* | Reston | RESTV | 22 |
| *O. sudanense* | Sudan | SUDV | 21 |
| *O. taiense* | Taï Forest | TAFV | 3 |
| *O. zairense* | Ebola | EBOV | 461 |

#### Species Hepacivirus hominis and genus Orthoflavivirus from the family Flaviviridae

*Flaviviridae* is a large family of positive single-stranded RNA viruses consisting of four genera: *Orthoflavivirus, Pestivirus, Pegivirus* and *Hepacivirus* [36], with a total of 97 species (ICTV, see Fig. S4). *Flaviviridae* includes human pathogenic viruses such as Zika virus, tick-borne encephalitis virus (TBEV), dengue virus, and hepatitis C virus (HCV). Genomes are approximately 9–13 kb in length [36]. We chose three datasets to compare results on species, genus, and family level. On the species level, we chose *Hepacivirus hominis* (HCV infecting humans) as a well-described and phylogenetically closely related species. On a genus level, we chose *Orthoflavivirus* due to the well-described dataset being very diverse (ICTV: 53 species). We chose *Flaviviridae* on the family level. The datasets *Hepacivirus hominis* and *Orthoflavivirus* are complete subsets of the dataset *Flaviviridae*. Finally, our focus was on how the selected representatives from species and genus study are consistent within the selection from the entire virus family. Preprocessing within the `ViralClust` pipeline reduced the datasets by 1–5 %, see Tab. 1 and 4.

**Table 4.**
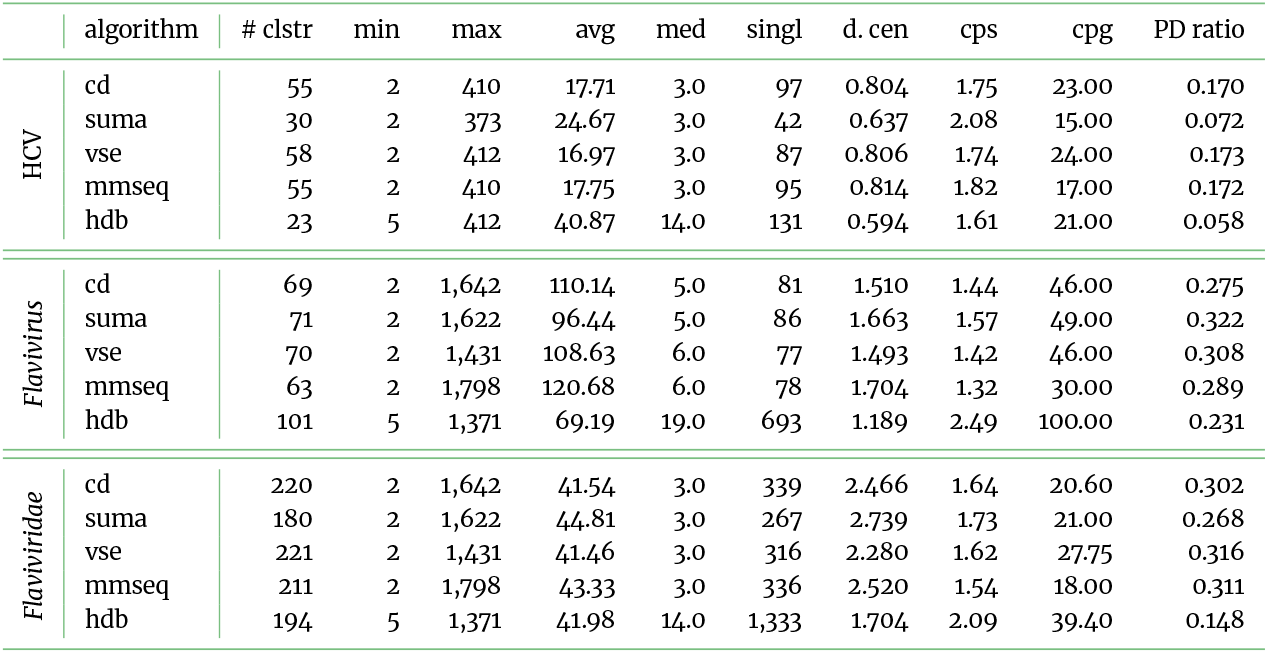
Cluster statistics for *Hepacivirus hominis* (HCV), *Orthoflavivirus* (*Flavivirus*), and *Flaviviridae* datasets. For each tool, the basic cluster statistics are listed. cd – `CD-HIT-EST`; sum – `SUMACLUST`; vse – `VSEARCH`; mmseq – `MMSeqs2`; hdb – `HDBSCAN`; # clstr – number of clusters (with a size greater than one); min/max – size of smallest/largest cluster; avg/med – average/median cluster size; d. cen – average phylogenetic distance between two representative genomes; singl – number of singletons (= cluster with only one element); cps – average cluster per species; cpg – average cluster per genus; blackPD ratio – ratio between Faith’s phylogenetic distance (PD) of the full tree and the representative tree. The values cps and cpg give a measurement of how much a single species/genus is divided into different groups.

##### Species Hepacivirus hominis

*Hepacivirus hominis* (HCV) consists of eight genotypes [37, 38, 39], however, only seven are listed in the ICTV, see Fig. S4 yellow rectangle. With the neighbour-net method we gained 13 clearly distinguishable clusters, see Fig. 3. This difference can also be explained by the observations of virologists in the past: (i) Two additional clusters, besides the seven downloaded genotypes, are formed by recombinants: the natural recombinants (e.g. 2k/1b or 2b/1b, Fig. 3 – close related to genotype 1b) [40, 41, 42] and the artificial recombinants (Fig. 3 – between genotype 1 and 2) [43, 44, 45]. Note: further recombinants are mixed into the genotype 2. (ii) Three additional clusters are formed within genotype 1 – in total, divided into genotype 1, 1a, 1b, and rare genotypes 1 (1c–1n). Currently, 14 subtypes are listed in the ICTV [36, 46, 47]; (iii) One additional cluster exists due to the diverse genotype 6: Currently, 33 subtypes are known for genotype 6 according to the ICTV [36, 48], with subtype 6a being the first one discovered [49]. All sequences part of subtype 6a cluster distinguishable from other subtypes, with the notable exception of subtype 6b, which more close to 6a than other subtypes. The tools of `ViralClust` calculated between 23 (`HDBSCAN`) and 58 (`VSEARCH`) representative genomes (see Fig. 3 colored circles, and Tab. 1). Although any of the methods represents 3–8 times more clusters than the currently recognized number of HCV genotypes, using more representatives may be necessary to capture the considerable diversity within genotypes. Among the evaluated methods, `HDBSCAN` produced the largest clusters (minimum size 5, average size 40.87, median size 14; see Tab. 4), resulting in the lowest total number of clusters. All clustering tools in `ViralClust` selected representative genomes distributed across the phylogenetic tree. However, substantial differences in the number of representatives were observed for the different genotypes, see Fig. 3. Interestingly, none of the tools was able to identify genotype 7 as an individual cluster.

**Figure 3.**
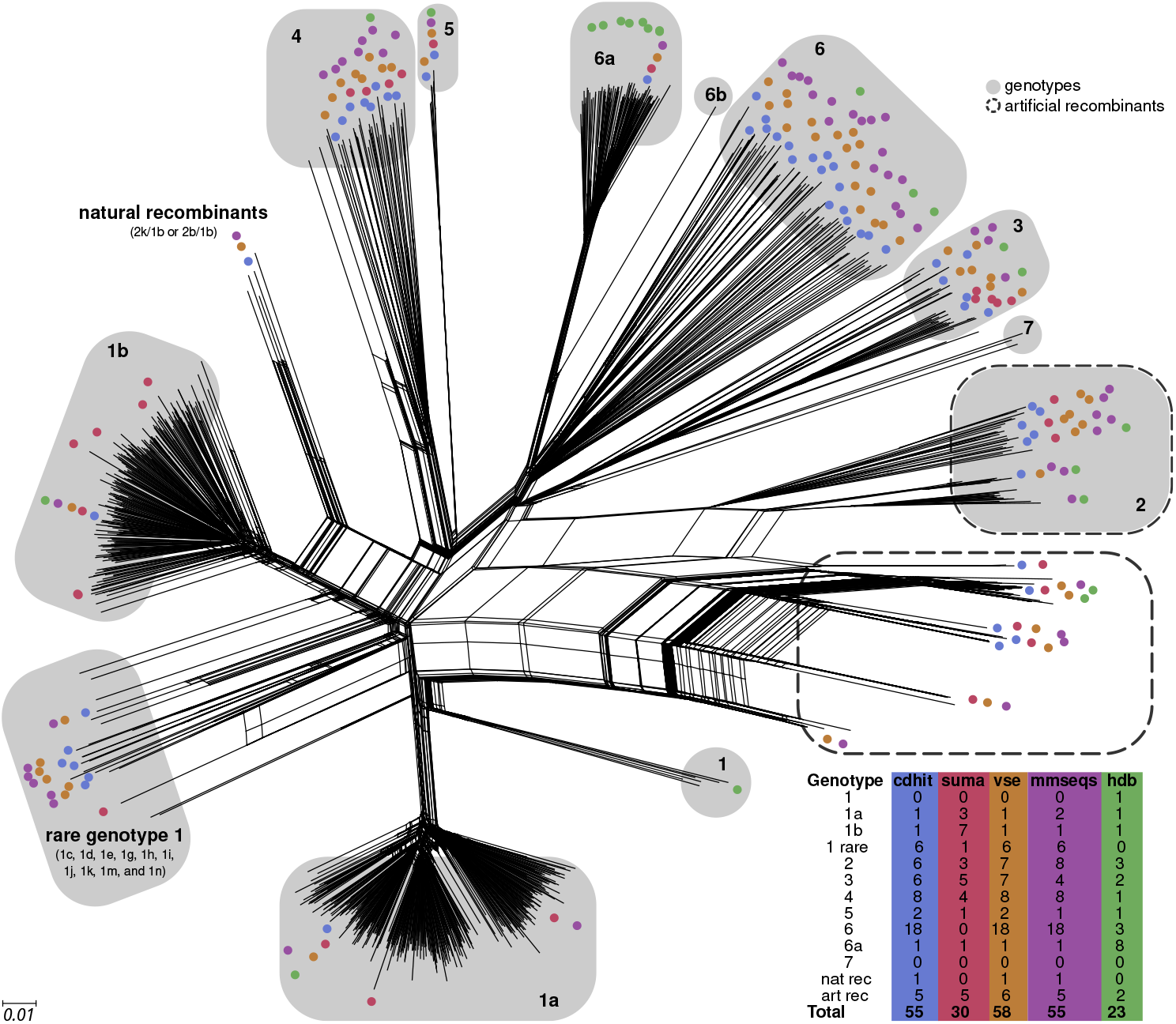
Splits graph of HCV with labeled genotypes (gray), artificial (dashed box) and natural recombinants, and cluster representatives of the five tools incorporated in `ViralClust` (`CD-HIT-EST`, `SUMACLUST`, `VSEARCH`, `MMSeqs2`, and `HDBSCAN`). Numbers in gray boxes represent genotypes. The tree can be divided into 13 subtrees, whereas genotype 1 consists of four subtrees (1, 1a, 1b, and the rare genotypes 1c–1n); and all subtype 6a genomes are clearly distinguishable from all other subtypes summarized here as genotype 6. All known natural recombinants (e.g. 2k/1b or 2b/1b) form a separate subtree most closely related to genotypes 1b and 4. Some artificial recombinants mix with genotype 2 genomes, whereas others form a subtree between genotypes 1 and 2. The number of representatives per genotype is shown in the table at the bottom right. The split graph is reconstructed using `SplitsTree` based on the `MAFFT` alignment of the complete genome sequences (1,071 genomes) resulting from the first filtering steps of `ViralClust`.

Across all five clustering algorithms, the largest cluster was consistently identified as genotype 1a. Each tool selected 1–3 representatives for this cluster. While four tools (`CD-HIT-EST`, `VSEARCH`, `MMSeqs2`, `HDBSCAN`) selected the same representative genome for genotype 1b, `SUMACLUST` identified seven. Notably, `HDBSCAN` did not select any rare subtypes of genotype 1, whereas the other tools included one (`SUMACLUST`) or six (`CD-HIT-EST`, `VSEARCH`, `MMSeqs2`) representatives. This is likely due to the k-mer–based similarity measure used by `HDBSCAN`, which appears to group the rare genotype 1 subtypes more tightly, leading to no additional representatives. A small subtree annotated as genotype 1 (Fig. 3, bottom right) was only found to be an independent cluster by `HDBSCAN`, which selected a single genome from these variants. The large number of subtypes within genotype 6 can be explained by the diversity of this genotype, see Fig. 3, even tough the tools showed picked different representatives. The tools `CD-HIT-EST`, `VSEARCH`, and `MMSeqs2` selected 18 representative genomes for genotype 6, whereas `HDBSCAN` identified substantially fewer representatives (3). These differences may be attributed to the distinct algorithmic strategies employed: `HDBSCAN` is based on k-mer similarity, while the other tools combine heuristic approaches with (pairwise) sequence alignments, which may increase sensitivity to finer sequence variation. Interestingly, `SUMACLUST` did not identify genotype 6 as a distinct cluster. This may be explained by the filtering step in `SUMACLUST`, which removes sequences containing non-ACGU characters and may have excluded genotype 6 genomes prior to clustering. In contrast, `HDBSCAN` selected eight representatives for subtype 6a, whereas the remaining tools selected only one. This difference may be influenced by the high abundance and relative homogeneity of subtype 6a genomes, in combination with the density-based clustering approach used by `HDBSCAN`. Several genomes in the dataset were labeled as recombinants leading to a split in the bottom right part of the tree (Fig. 3, dashed box) and potentially inflating the number of clusters. These recombinant genomes are annotated as a mix between genotypes 1 and 2. Overall, all tools except `HDBSCAN` produced broadly comparable results. While `HDBSCAN` tended to select more representatives in densely connected regions (e.g. genotype 6a), it selected fewer genomes in other parts of the tree. When examining individual sequences, they are usually assigned to the same cluster, although the chosen representative may differ slightly, see Fig. 4.

**Figure 4.**
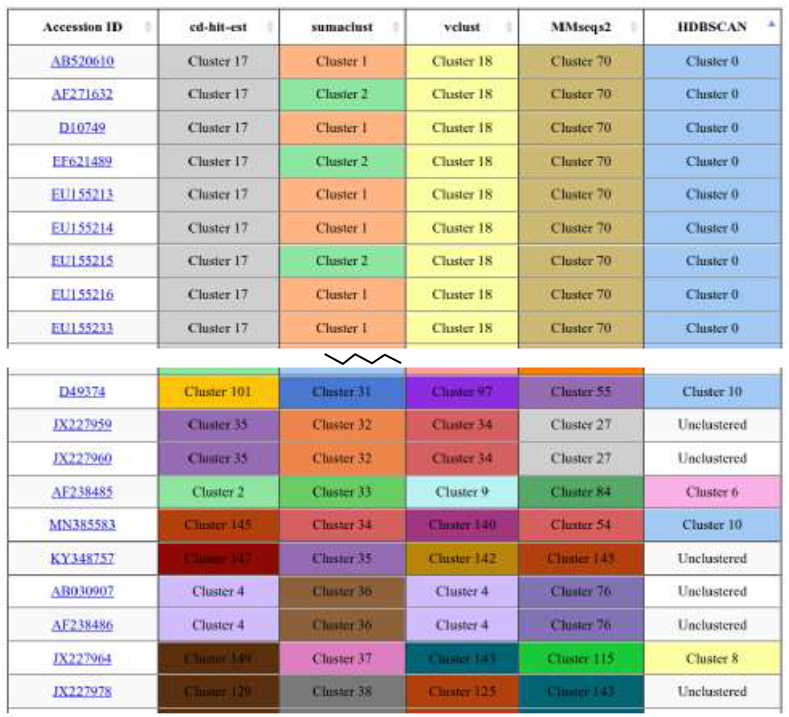
Optional output of `ViralClust` in web-based format (HTML) for the dataset HCV. The interactive visualization allows the user to (i) sort the table by tool results or accession IDs, (ii) view and compare color-coded clusters, and (iii) access the NCBI entry for each sequence (link, if available). For most sequences, the clustering results of the different tools are consistent (top). Differences between the tools become apparent (bottom), where some sequences form single-sequence clusters (e.g. `D49374`), others consistently form small clusters across tools (e.g. `AB030907` and `AF238486`), and individual sequences are not assigned to any cluster by specific tools, such as the `HDBSCAN`’unclustered’ sequences. The complete table is provided in the electronic supplement.

##### Genus Orthoflavivirus

The clustering of 8,052 *Orthoflavivirus* resulted in 63–101 clusters with the different tools, whereas `MMSeqs2` serves with the smallest number of clusters and `HDBSCAN` with the highest, see Tab. 4. As for HCV, the number of clusters are higher than the number of species (53). The selection of more than one genome per species as representative can be sufficient to cover greater diversity across the whole genus of *Orthoflavivirus*. Despite minor differences, all five clustering algorithms produced broadly similar results, with representatives distributed across the phylogenetic representation of the dataset (see Fig. S5).

When comparing the selected representatives with the ICTV species assignments, which contains exactly one reference genome per species, see Fig. S4 violet background, we find (a) clusters which contain exactly one of the defining ICTV species, as expected; (b) ICTV species representatives not part of any cluster (= singletons); (c) clusters, which have no defining ICTV species, which might point to a generally more fine-grained clustering; and (d) some clusters have been merged by specific tools, so that two (or more) ICTV species could be found in one cluster, which should not have happened in order to represent the current phylogeny well.

The different tools found between 16 and 23 clusters containing exactly one ICTV species representative, see Tab. 5, whereas 18–23 ICTV reference genomes were not assigned to any cluster with more than one sequence, and thus remain as singletons within the `ViralClust` pipeline. This likely reflects recently added species for which only limited sequence data are currently available. An important detail is that `SUMACLUST` excludes any sequences containing characters other than `A`, `C`, `G`, or `T`. As a result, the ICTV representative genome for St. Louis encephalitis was removed from the dataset during preprocessing. Between 45 and 84 clusters contained no ICTV reference genome. These clusters likely represent intraspecific diversity or unclassified lineages, providing additional resolution beyond the current ICTV taxonomy. `CD-HIT-EST`, `SUMACLUST`, `VSEARCH`, and `MMSeqs2` agreed to cluster two reference genomes of ICTV together into one cluster: Saboya (`NC_033697`) together with Potiskum (`NC_029054`). `HDBSCAN` also merged Saboya (`NC_033697`) together with other species, but in this case together with Modoc (`NC_003635`), Uganda S (`NC_033698`), and Wesselsbron (`EU707555`). For a more details see Tab. S4 and S5. However, if higher resolution is required, we recommend including singletons as well. In this case, users should be aware that incorporating singletons may lead to over-representation of certain regions within the dataset. `HDBSCAN` often adapts to the density of subtrees: for very dense clusters, it selects several representatives (see Fig. S5 top right), whereas clusters consisting of a single sequence are typically not chosen as representatives and instead remain unassigned. If such outliers are of interest for downstream analysis, users should manually include these singletons in subsequent steps (as these sequences are saved in a separate file). As a result, `HDBSCAN` yields the lowest average cluster size (69.19) and the largest number of singletons (693). An important observation concerns the previously proposed subclassification within *Orthoflavivirus* according to their vector type [36]. These analyses are based solely on parts of the the RdRp, see Fig. S4. Our analysis is based on complete genomes, indicating at least for subtrees, that the phylogeny can not be drawn based on the vector-type, see Fig. 5.

**Figure 5.**
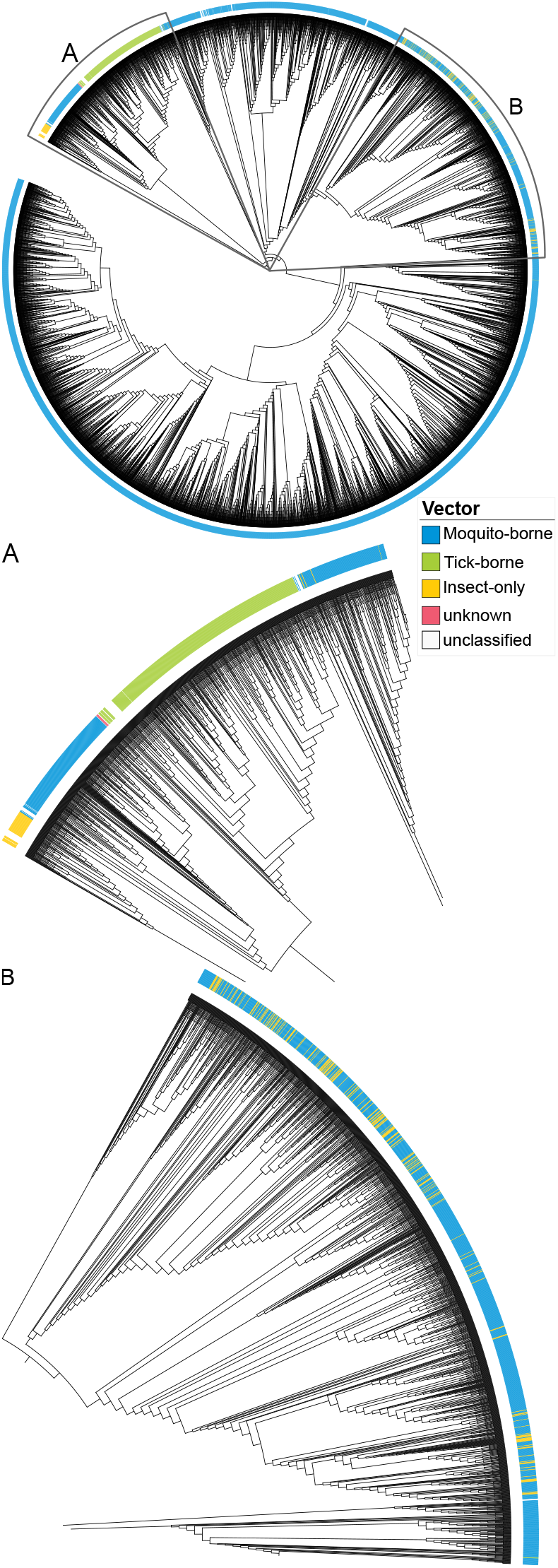
Phylogenetic representation based on the *Orthoflavivirus* dataset with color-coded vectors. Within the subtrees A and B, parts of the tree do not follow a vector-based classification. The complete and enlargeable tree is shown in Fig. S6. The tree is reconstructed using `FastTree` based on the `MAFFT` alignment of the complete genome sequences (7,681 genomes) resulting from the first filtering steps of `ViralClust`.

**Table 5.** Comparing the clustering results of the *Orthoflavivirus* dataset with the species chosen for the phylogeny of ICTV, see Fig. S4. 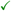 – number of clusters with exactly 1 ICTV representative (in agreement with known phylogeny); Singleton – Number of clusters on containing one ICTV representative; 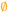 – number of clusters without ICTV representative (showing additional information for downstream analysis); Discrepancies – (a) number of cluster with >= 2 ICTV representatives (e.g., ‘1×4’ for four ICTV representatives in one cluster); (b) number of ICTV representatives that are ‘singletons’ (these sequences are alone a cluster); (c) other issues (e.g., `SUMACLUST` removes sequences including nucleotides *n* ∉ {*A, C, G, T*}). For a more detailed overview see Tab. S4 and S5.

|  | ✓ | Singleton | Ø | Discrepancies |
| --- | --- | --- | --- | --- |
| cd | 23 | 18 | 45 | 1×2 |
| suma | 20 | 20 | 50 | 1×2<br>St. Louis encephalitis<br>(DQ525916) removed |
| vse | 23 | 18 | 46 | 1×2 |
| mmseq | 23 | 18 | 39 | 1×2 |
| hdb | 16 | 23 | 84 | 1×4 |

##### Family Flaviviridae

Clustering of all 9,895 *Flaviviridae* species resulted in 180–221 clusters with `SUMACLUST` generating the lowest number of clusters and `VSEARCH` the highest, see Tab. 4. These results are twice as much as the reported 97 species from ICTV, Fig. S4, as for the subsets HCV species and *Orthoflavivirus* genus.

The average cluster size remains about the same for all five algorithms. `MMSeqs2` produced the overall largest cluster (1,798 sequences), whereas `HDBSCAN` (1,371) and `SUMACLUST` (1,431) produced the smallest, with 200–400 fewer sequences. `HDBSCAN` also produced the largest number of singletons, with a notably high fraction (1,333), see Tab. 4. The ICTV phylogeny (Fig. S4) was constructed using the RdRp region of 125 sequences [36]. To evaluate how well this single-gene phylogeny reflects whole-genome evolutionary relationships, we compared it against a phylogeny inferred from 9,895 complete *Flaviviridae* genomes. While the overall topology of the full genome based phylogeny aligns broadly with the RdRp based ICTV classification, Fig. S7, several notable discrepancies emerged. Specifically, as visible in Fig. 6 (from left to right) (i) 25 genomes annotated as orthoflaviviruses clustered together in a distinct outlier subtree; (ii) 94 HCV genomes appeared more closely related to most of the pegiviruses than to other hepaciviruses; (iii) a single *Pestivirus* genome (Trinbago virus) was located within this divergent HCV cluster (not visible); (iv) another three, more divergent, *Pegivirus* sequences clustered with the majority of HCV; and (v) nine *Orthoflavivirus* genomes were placed nearer to basal pestiviruses (Fig. 6A – blue part between yellow and green). These discrepancies may be partly explained by misannotations in public sequence databases, but they also highlight the inherent limitations of single-gene phylogenies for robust virus taxonomy.

**Figure 6.**
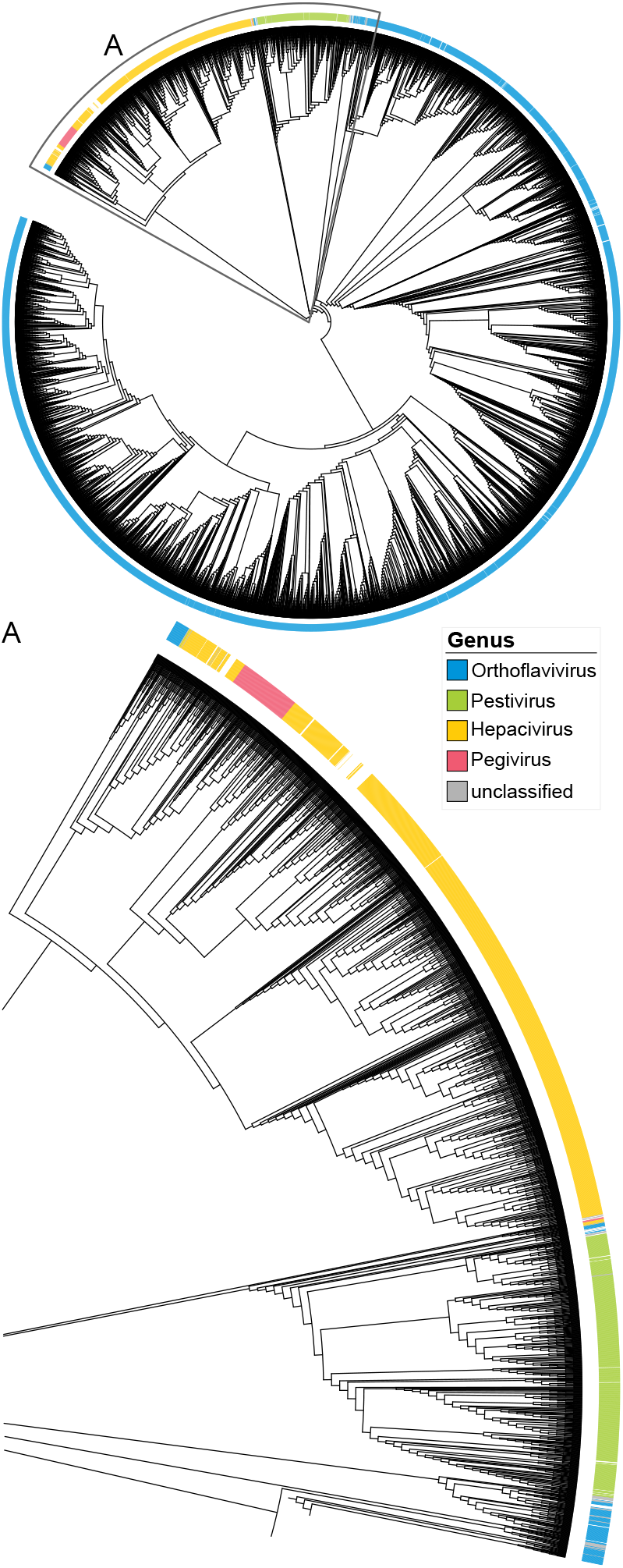
Phylogenetic representation of the *Flaviviridae* dataset with highlighted genera. A) parts of the tree that do not follow the classification by genus. Complete tree shown in Fig. S7. The tree is reconstructed using `FastTree` based on the `MAFFT` alignment of the complete genome sequences (9 478 genomes) resulting from the first filtering steps of `ViralClust`.

##### Comparing the datasets

We chose *Flaviviridae* on the family level (9,895 genomes) to test the robustness of the different clustering approaches, especially in comparison to clustering on the species level (HCV, 1,075 genomes true subset of *Flaviviridae*) and on the genus level (*Orthoflavivirus*, 8,052 genomes true subset of *Flaviviridae*), see Tab. 1. The fine-grained clustering observed at the species level is for some clustering algorithms also preserved at the family level, see Tab. 6. For example, within genomes of HCV, the methods `CD-HIT-EST`, `SUMACLUST`, and `VSEARCH` consistently formed the same clusters and selected the same representatives when compared between the HCV only and HCV clustered together with all other *Flaviviridae* datasets. `MMSeqs2` shows slight differences in the selection of representatives, see Fig. S8. Clustering the entire *Flaviviridae* family with `MMSeqs2` produced 56 clusters for HCV, compared to 55 clusters when only the HCV species was analyzed. Of these, 54 clusters were identical, while one cluster in the HCV clustering was split into two in the *Flaviviridae* analysis. Among the representatives, 49 were identical, and the remaining non-identical ones were phylogenetically close. In contrast, `HDBSCAN` showed larger differences in both cluster composition (only one identical cluster) and choice of representatives (six identical). When restricted to HCV, `HDBSCAN` produced twice as many clusters (23) compared to when HCV was analyzed as part of the *Flaviviridae* dataset (12). Genotypes 1a and 1b are represented by multiple genomes in the clustering of HCV, while these are grouped together in the results of *Flaviviridae*. This suggests that `HDBSCAN` is more sensitive to the chosen level and number of input sequences and adapts its clustering accordingly. When switching to the comparison on a genus level at the example of *Orthoflavivirus* only vs. *Orthoflavivirus* within *Flaviviridae*, all five algorithms generated the same largest cluster, containing dengue virus 1, see Tab. 4. `SUMACLUST` and `VSEARCH` again produced identical clusters and representatives, indicating robustness to the number of input sequences, see Tab. 6. For the genus *Orthoflavivirus*, `CD-HIT-EST` and `MMSeqs2` selected the same total number of representatives – 69 and 63 respectively, as in the corresponding *Orthoflavivirus* subsets of the *Flaviviridae* analysis. While `CD-HIT-EST` produced identical cluster compositions, 8 representatives differ. Similarly, `MMSeqs2` yielded 56 clusters with unchanged composition, but 48 identical representatives. In both cases, the differing representatives were phylogenetically close, and thus these discrepancies are considered minor (Fig. S9). In contrast, `HDBSCAN` generated substantially more clusters for *Orthoflavivirus* when clustering within the full *Flaviviridae* dataset (156 representatives) compared to clustering *Orthoflavivirus* alone (101 representatives). Of these, 27 clusters and 80 representatives were identical. The largest differences were observed in the mosquito-borne clade, where `HDBSCAN` selected considerably more representatives than the other algorithms (Fig. S9). The run time does not scale linearly with dataset size, see Tab. S1. Although the *Orthoflavivirus* dataset accounts for 81 % of the *Flaviviridae* sequences (7,681 of 9,478), `ViralClust` required only 48.5 % of the total run time (1 h 22 m of 2 h 47 m; Tab. 1). For the HCV dataset, the effect was less pronounced: comprising 11.3 % of the sequences in the *Flaviviridae* dataset, it required 8.3 % of the run time (14 m of 2 h 47 m; Tab. 1). The overall memory consumption scales almost linearly when comparing the *Flaviviridae* dataset with the *Orthoflavivirus* dataset (183.9 G, 164.3 G; Tab. S1). In contrast, this is not the case for the HCV dataset, where memory usage is four times higher than for *Flaviviridae* when comparing with the size of the datasets.

**Table 6.** Comparison of the clustering results of *Hepacivirus hominis* only vs. *Hepacivirus hominis* within *Flaviviridae* and *Orthoflavivirus* only vs. *Orthoflavivirus* within the *Flaviviridae* analysis. clstr-comp. – cluster composition, i.e. number of identical clusters (number of cluster in *Hepacivirus hominis* or *Orthoflavivirus*/number of cluster in *Flaviviridae*); repr – identical representatives; 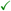 – results are identical (id.). cd – `CD-HIT-EST`; sum – `SUMACLUST`; vse – `VSEARCH`; mmseq – `MMSeqs2`; hdb – `HDBSCAN`.

| *Hepacivirus hominis*: only vs. within *Flaviviridae* |  |  |
| --- | --- | --- |
|  | clstr-comp. | repr |
| cd | ✓ (56 clstr) | ✓ |
| suma | ✓ (31 clstr) | ✓ |
| vse | ✓ (59 clstr) | ✓ |
| mmseq | 54 id. (55/56) | 49 id. |
| hdb | 1 id. (23/12) | 6 id. |
| *Orthoflavivirus*: only vs. within *Flaviviridae* |  |  |
|  | clstr-comp. | repr |
| cd | ✓ (69 clstr) | 57 id. |
| suma | ✓ (71 clstr) | ✓ |
| vse | ✓ (70 clstr) | ✓ |
| mmseq | 56 id. (63/74) | 48 id. |
| hdb | 27 id. (101/158) | 80 id. |

#### Alphainfluenzavirus influenzae as an example for segmented viruses including partial and reverse-complement sequences

*Alphainfluenzavirus influenzae* (formerly *Influenza A virus*, IAV) provides a representative case for studying segmented viral genomes and handling truncated reverse-complement sequences. Here, we address several key questions relevant for sequence clustering and analysis of such viruses: We first confirm that genotype represents surprisingly well to serotype classification. For the eight genome segments, sequences can be analyzed individually or concatenated; here we show that independent analysis captures meaningful differences. Partial genomes are reliably placed in phylogenetic trees, demonstrating robustness. Finally, reverse-complement sequences are detected through characteristic patterns (in this section highlighted with gray background); the current version of `ViralClust` automatically restores the positive orientation to simplify analysis.

IAV is a segmented (8 segments), negative-sense, single-stranded RNA virus belonging to the family *Orthomyxoviridae* (see Fig. S10). Classification is traditionally based on the two surface proteins hemagglutinin (H, segment 4) and neuraminidase (N, segment 6), with 18 H and 11 N subtypes reported by the CDC (see Fig. S11 and S12). Our analyses confirm that segment data reflect subtype classification well, demonstrating that genotype approximates subtype and that segment-wise analysis captures meaningful biological differences.

Each of the eight genome segments in our dataset contains on average 98,532 sequences, with the largest datasets corresponding to segments 4 (156,586) and 6 (114,639) and the smallest to segment 2 (82,569). Segment lengths range from 890 nt (segment 8) to 2,341 nt (segments 1 and 2), and non-redundant sequence fractions vary from 64—69 % for segments 1–4 and 6 to 47–49 % for segments 7 and 8.

Clustering results (90 % sequence identity, k-mer length 7) vary widely depending on segment and method, ranging from 25 clusters (`CD-HIT-EST`, segment 7) to 2,307 (`HDBSCAN`, segment 4), see Tab. 1, with the largest datasets (segments 4 and 6) also yielding the highest number of clusters across all five tools. blackThis wide range illustrates that tool choice has a substantial impact on the granularity of the resulting representative set, independent of the underlying biological diversity. `CD-HIT-EST` produced the lowest number of clusters for all segments, followed by `VSEARCH`. `MMSeqs2` and `SUMACLUST` are close to each other in terms of the number of clusters. `MMSeqs2` yielded a lower number of clusters for segments 1, 2, 3, 5 and 6, but more for 4, 7 and 8. `HDBSCAN` resulted in the highest number of clusters by far. blackThe consistently lower cluster counts of `CD-HIT-EST` and `VSEARCH` reflect their more aggressive merging strategies, which may be advantageous when the goal is maximal dataset reduction.

blackThe coverage of known subtypes provides a more biologically grounded basis for comparing tool performance than cluster counts alone. For segments 4 and 6, which encode the surface proteins, `CD-HIT-EST`, `SUMACLUST`, `VSEARCH`, and `MMSeqs2` select representatives covering all known subtypes, see Tab. 7 and Fig. 7 and 9. blackIn contrast, `HDBSCAN` fails to select representatives for H17 and H18 (segment 4) and N10 and N11 (segment 6). These are among the rarest subtypes in the dataset, and their absence from the `HDBSCAN` representative set is likely a fundamental consequence of density-based clustering: subtypes represented by very few sequences may not form sufficiently dense regions to be recognized as independent clusters, and are instead treated as noise.

**Figure 7.**
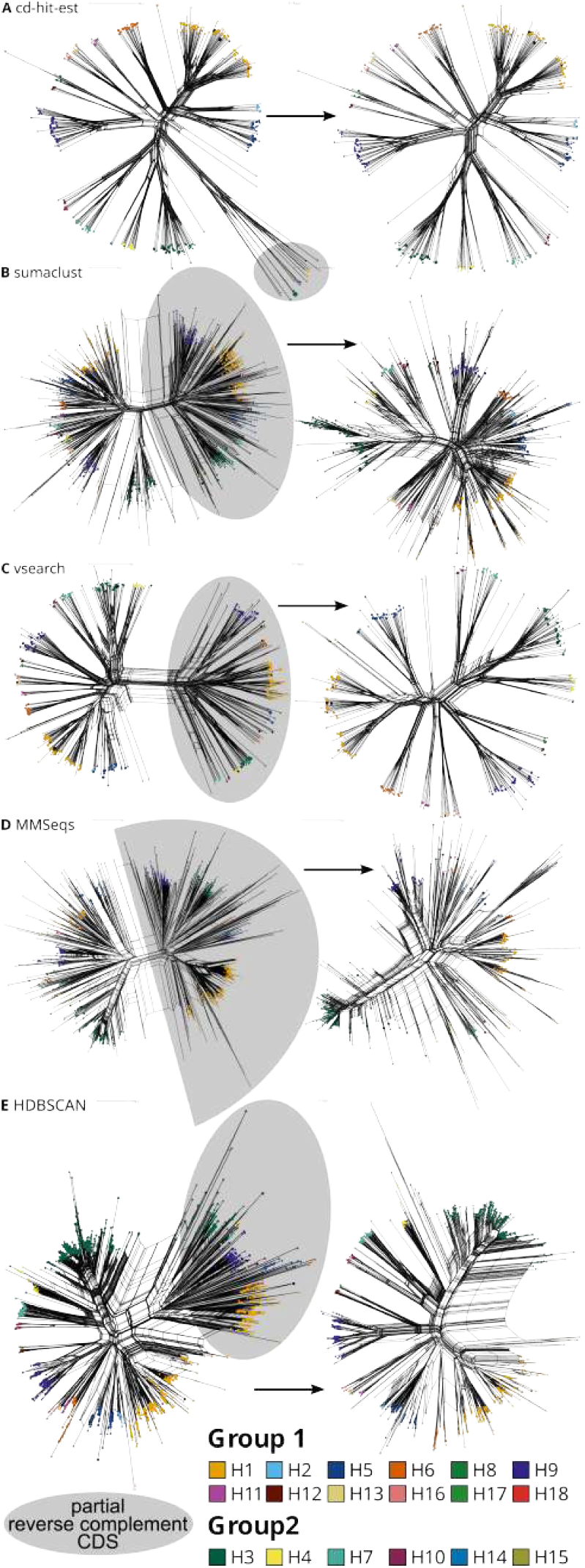
Split graphs of IAV segment 4 (H) representatives of (A) `CD-HIT-EST`, (B) `SUMACLUST`, (C) `VSEARCH`, (D) `MMSeqs2`, and (E) `HDBSCAN`. Left – all representatives (including partial and reverse-complement); Right – only complete genomes. The node size indicates cluster size. The split graphs are reconstructed using `SplitsTree` based on the `MAFFT` alignments. blackGray background highlights reverse-complement sequences (19.22 % of the non-redundant dataset). `HDBSCAN` did not select representatives for subtypes H17 and H18. They were manually and marked them as framed circles. For details see Fig. S13–S17.

**Table 7.** Number of representatives per subtype for segment 4 (H) and 6 (N). #repr number of representatives (= number of clusters with a size greater than one); cd – `CD-HIT-EST`; sum – `SUMACLUST`; vse – `VSEARCH`; mmseq – `MMSeqs2`; hdb – `HDBSCAN`.

| Subtype | #repr |  |  |  |  |
| --- | --- | --- | --- | --- | --- |
|  | cd | suma | vse | mmseq | hdb |
| Segment 4 (H) |  |  |  |  |  |
| H1 | 83 | 352 | 128 | 527 | 613 |
| H2 | 5 | 26 | 10 | 19 | 35 |
| H3 | 34 | 215 | 59 | 460 | 654 |
| H4 | 14 | 44 | 22 | 28 | 58 |
| H5 | 27 | 145 | 43 | 107 | 210 |
| H6 | 19 | 54 | 27 | 33 | 67 |
| H7 | 16 | 37 | 22 | 23 | 100 |
| H8 | 5 | 9 | 7 | 5 | 11 |
| H9 | 44 | 193 | 71 | 242 | 347 |
| H10 | 9 | 27 | 11 | 17 | 42 |
| H11 | 7 | 16 | 10 | 13 | 19 |
| H12 | 4 | 9 | 7 | 3 | 22 |
| H13 | 8 | 20 | 11 | 13 | 21 |
| H14 | 2 | 1 | 1 | 1 | 1 |
| H15 | 2 | 2 | 2 | 2 | 2 |
| H16 | 6 | 13 | 10 | 9 | 13 |
| H17 | 1 | 1 | 1 | 1 | 0 |
| H18 | 1 | 1 | 1 | 1 | 0 |
| Segment 6 (N) |  |  |  |  |  |
| N1 | 47 | 250 | 73 | 198 | 476 |
| N2 | 90 | 280 | 130 | 276 | 733 |
| N3 | 17 | 29 | 21 | 21 | 27 |
| N4 | 6 | 15 | 10 | 5 | 17 |
| N5 | 7 | 14 | 7 | 10 | 30 |
| N6 | 15 | 45 | 24 | 33 | 94 |
| N7 | 12 | 31 | 19 | 17 | 29 |
| N8 | 20 | 52 | 30 | 26 | 103 |
| N9 | 5 | 20 | 6 | 15 | 37 |
| N10 | 1 | 1 | 1 | 1 | 0 |
| N11 | 1 | 2 | 1 | 2 | 0 |

Partial genomes are reliably placed at the correct positions within phylogenetic trees, demonstrating the robustness of our approach even when sequences are incomplete. blackReverse-complement sequences, however, present a more disruptive challenge. blackComprising 19.22 % of the segment 4 dataset and 11.37 % of the segment 6 dataset, they arrange in a characteristic ‘mirror-like’ configuration in the phylogenetic tree. Such a pattern, if undetected, would cause clustering algorithms to treat forward and reverse-complement copies of the same sequence as unrelated, artificially inflating cluster counts and distorting representative selection. Therefore, we implemented `ViralClust` to detect reverse-complement sequences and to automatically restore their original orientation, simplifying downstream analysis, ensuring that strand orientation artifacts do not propagate into downstream analyses. blackTo quantify the impact of reverse-complement sequences on clustering results, we compared three scenarios for segments 4 (H) and 6 (N): clustering the full dataset including reverse-complement sequences, removing reverse-complement representatives post-hoc from the original results, and re-clustering after removing all reverse-complement sequences from the input (see Tab. S6). The presence of reverse-complement sequences substantially inflated the number of representatives across all tools, demonstrating that a considerable fraction of clusters in the uncorrected results arose purely from strand orientation artifacts rather than genuine sequence diversity. Notably, post-hoc removal of reverse-complement representatives and full re-clustering produced comparable results for most tools, suggesting that `ViralClust`’s automatic orientation correction prior to clustering is sufficient to avoid these artifacts without requiring a separate filtering step.

Given the variable results across tools and segments, we adjusted parameter ranges of `ViralClust` accordingly, with sequence similarity threshold between 70 and 90 %, see Tab. S7. `CD-HIT-EST` only allows for a minimum of 80 %. In the case of `HDBSCAN`, we tested k-mers from 1 to 7, see Tab. S8. blackThe choice of sequence similarity threshold directly reflects the biological question: a higher threshold (e.g. 90 %) retains finer sequence-level differences and is appropriate when the goal is to collapse near-identical sequences while preserving all circulating variants within a subtype, such as for outbreak subsampling. A lower threshold (e.g. 70 %) merges more divergent sequences into the same cluster and is better suited for broader comparative analyses, such as selecting one representative per subtype or lineage across a diverse dataset. For *Alphainfluenzavirus influenzae*, where 18 H and 11 N subtypes span considerable sequence diversity, the threshold choice determines whether the resulting representative set captures subtype-level or strain-level diversity. As expected, the number of clusters decreases with decreasing similarity threshold for `CD-HIT-EST`, `SUMACLUST`, and `VSEARCH` (see Fig. 8 and S23), blackconfirming that these tools respond predictably to threshold changes and offer users intuitive control over clustering granularity. blackThis predictable behavior makes threshold selection straightforward for `CD-HIT-EST`, `SUMACLUST`, and `VSEARCH`: users can inspect the resulting cluster counts at different thresholds, as reported in Tab. S7, and select the threshold that best matches the expected number of biological units for their dataset. `MMSeqs2` using the default module implemented in `ViralClust`, however, shows an atypical pattern: cluster counts remain roughly stable or occasionally increase at lower thresholds. We hypothesize that this behavior stems from the `easy-linclust` mode used within `ViralClust`, which is optimized for speed and may not handle the full combinatorial space of pairwise comparisons in datasets of this scale, leading to threshold-insensitive results, despite the fact that this module is described to be designed to handle large datasets. Users working with similarly large virus datasets should be aware of this limitation and consider validating `MMSeqs2` results against those of other tools. blackMoreover, users can choose the `easy-cluster` module in `ViralClust` instead (`--mmseqs_mode cluster`). Among the tools evaluated, `CD-HIT-EST` and `VSEARCH` offer the most predictable and computationally tractable behavior for large segmented virus datasets, while `HDBSCAN` provides the finest resolution but at the cost of potentially missing rare subtypes. `SUMACLUST` and `MMSeqs2` occupy intermediate positions, with `MMSeqs2 easy-linclust` requiring additional caution at large dataset sizes.

**Figure 8.**
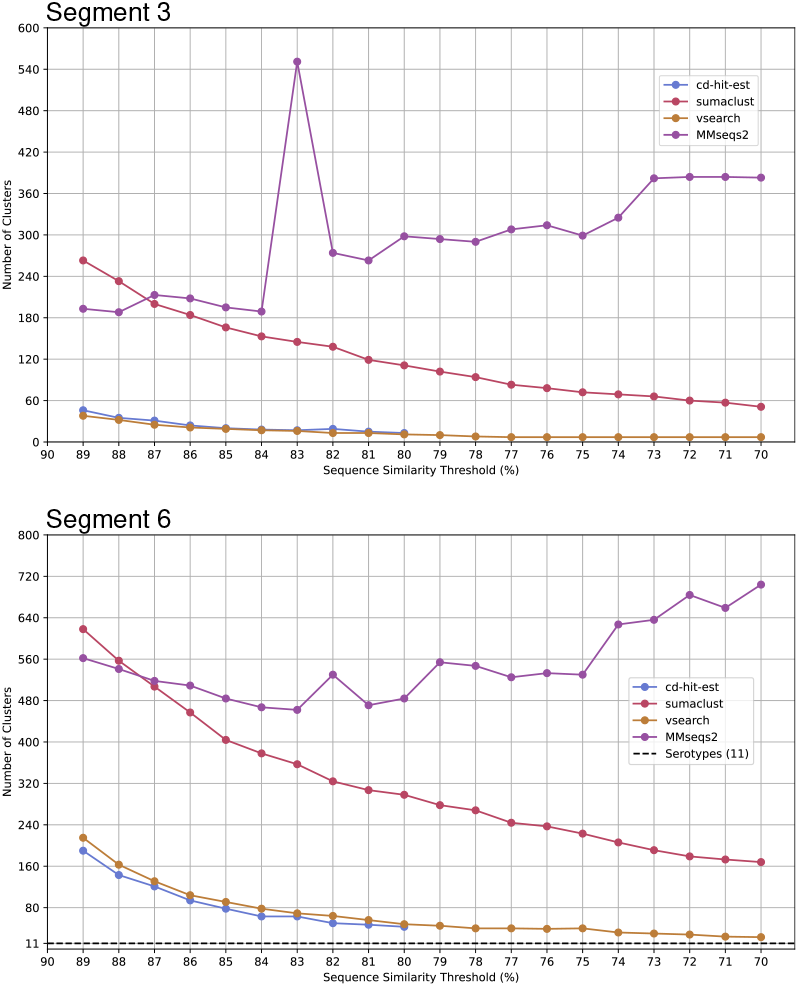
Number of clusters of IAV segments 3 and 6 with sequence similarity thresholds from 70 - 90 %. `CD-HIT-EST` only allows a minimum of 80 %. Only clusters with with a size greater than one are considered. For more details see Tab. S7 and S8.

**Figure 9.**
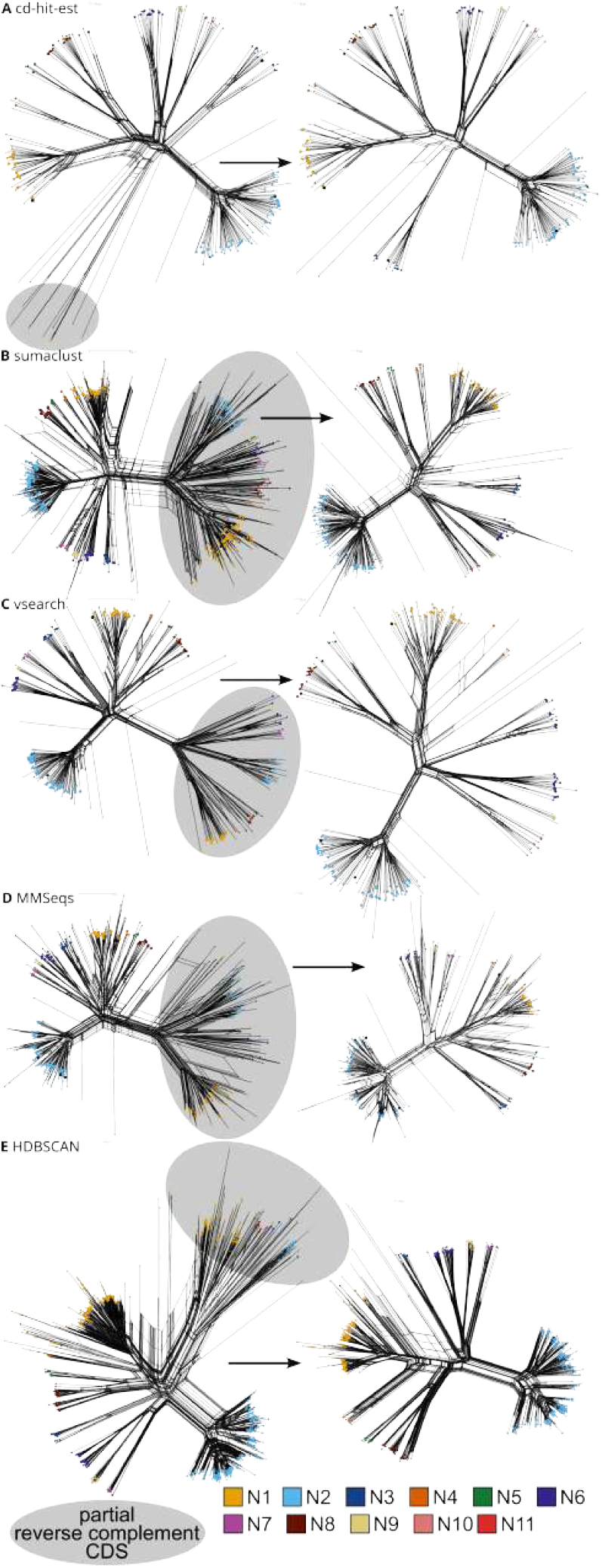
Split graphs of IAV segment 6 (N) representatives of (A) `CD-HIT-EST`, (B) `SUMACLUST`, (C) `VSEARCH`, (D) `MMSeqs2`, and (E) `HDBSCAN`. Left – all representatives (including partial and reverse-complement); Right – only complete genomes. The node size indicates cluster size. The split graphs are reconstructed using `SplitsTree` based on the `MAFFT` alignments. blackGray background highlights reverse-complement sequences (11.37 % of the non-redundant dataset). `HDBSCAN` did not select representatives for subtypes N10 and N11. They were manually and marked them as framed circles. For details see Fig. S18–S22.

#### Monkeypox virus as DNA virus example

To assess the challenges posed by long, repetitive DNA genomes to clustering algorithms, we downloaded 1,077 *Monkeypox virus* genomes (family *Poxviridae*, see Fig S24), with an average length of 197.185 kbp. 92.76 % of the sequences are non-redundant. The main challenge compared to the RNA virus datasets analyzed previously is the combination of much longer genome length and repetitive regions, which can affect both clustering performance and representative selection.

For `VSEARCH` and `CD-HIT-EST`, we increased the parameters to accommodate the longer *Monkeypox virus* genomes: `--maxseqlength 300000` (default: 50,000) and `-M 2500` (default: 400). This dataset required the most time and memory overall. Based on the two known genotypes (clade I and II), a biologically meaningful clustering would be expected to produce two or more clusters capturing this known diversity. `SUMACLUST`, `HDBSCAN`, and `VSEARCH` produced between 1 and 3 clusters, whereas `CD-HIT-EST` (14) and `MMSeqs2` (23) generated substantially more, see Tab. 1. blackThe behavior of `SUMACLUST` is particularly noteworthy: approximately 66 % of sequences (662 genomes) were discarded during clustering, most likely due to non-`ACGT` characters in repetitive genomic regions. This behavior causes the results from `SUMACLUST` to be unusable for subsequent analyses, since the cluster representatives cannot serve as representatives for the original dataset. Although `CD-HIT-EST` generated more clusters, it assigned 917 sequences to a single cluster. In contrast, `HDBSCAN` and `VSEARCH` produced largest clusters containing 566 and 565 sequences, respectively. `HDBSCAN` successfully assigned all sequences to clusters, while the other algorithms left between 1 and 69 sequences unclustered (`SUMACLUST` and `VSEARCH`: 1, `CD-HIT-EST`: 10, `MMSeqs2`: 69). `CD-HIT-EST` and `MMSeqs2` selected more representatives than the other tools. This difference likely reflects the underlying algorithmic principles: `CD-HIT-EST` uses a greedy incremental clustering approach that tends to select fewer representatives by aggressively merging similar sequences, whereas `MMSeqs2` implements a more sensitive, iterative clustering procedure that preserves more distinct representatives, especially in large, repetitive genomes. blackWhether finer granularity is desirable depends on the research question. For genotype- or subtype-level analyses, the higher resolution of `CD-HIT-EST` or `MMSeqs2` may be advantageous, while the compact output of `HDBSCAN` or `VSEARCH` may be preferable when the goal is maximal dataset reduction. The majority of the *Monkeypox virus* dataset consists of genotype IIb (see Fig. 10). The representatives selected by `SUMACLUST` and `HDBSCAN` reflect this dominant genotype, whereas `VSEARCH` representatives cover genotypes Ia and IIb. Both `CD-HIT-EST` and `MMSeqs2` resulted in representatives covering all genotypes present in the dataset. blackThis result highlights a broader principle: tools that produce more clusters are not inherently better or worse, but in outbreak-biased datasets dominated by a single genotype, higher cluster granularity may be necessary to ensure that minority lineages are retained in the representative set. For the *Monkeypox virus* dataset, `VSEARCH` offers the most complete genotypic coverage while preserving a number of clusters close to the number of genoytpes. `CD-HIT-EST` and `MMSeqs2` therefore offer a finer resolution, which may be because of their higher sensitivity to sequence-level variation in repetitive genomic regions.

**Figure 10.**
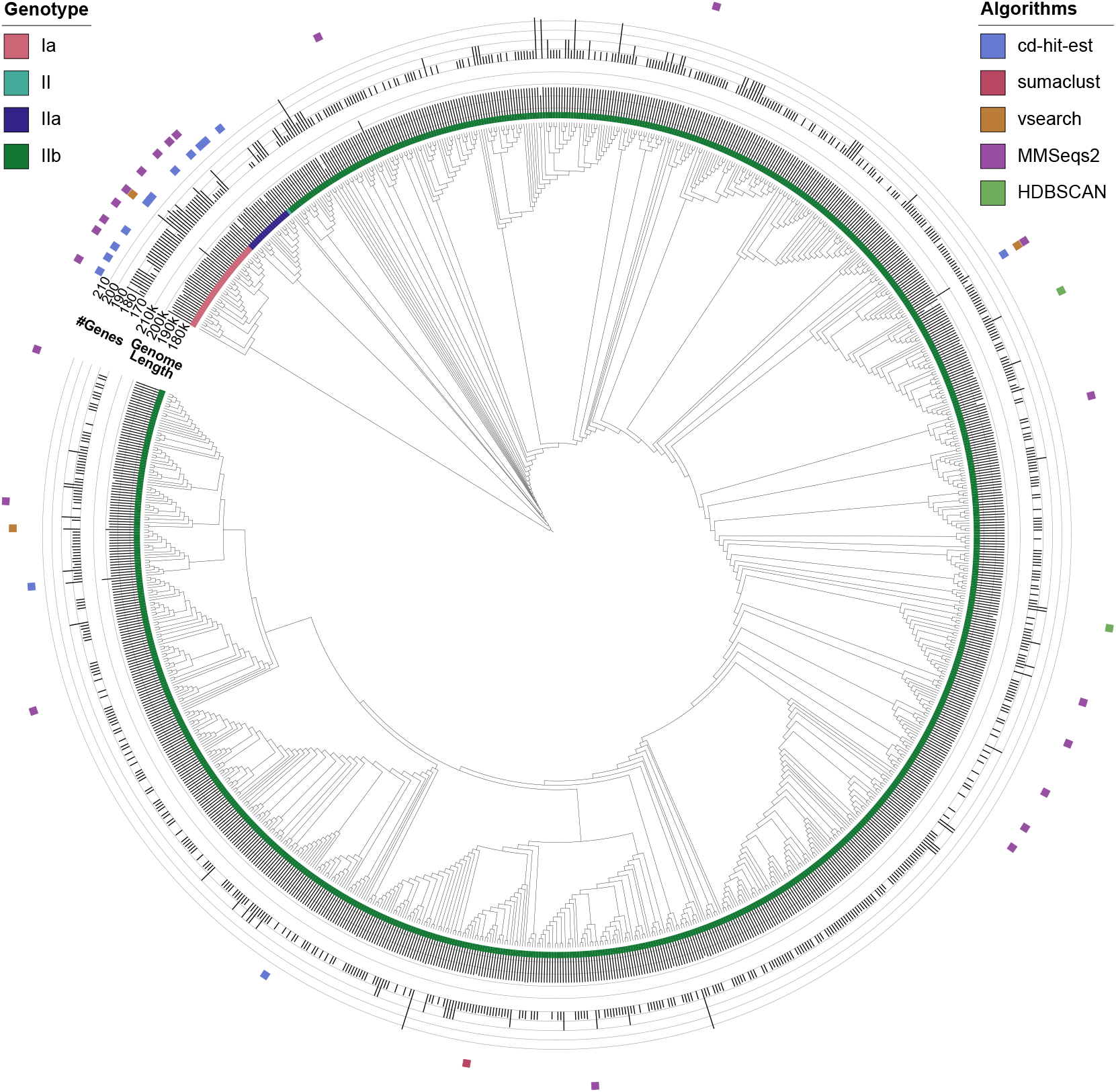
Phylogenetic reconstruction of the *Monkeypox virus* dataset with (i) genotypes, (ii) genome length, (iii) number of genes, and (iv) representative sequences selected by `CD-HIT-EST`, `SUMACLUST`, `VSEARCH`, `MMSeqs2`, and `HDBSCAN`. The tree is reconstructed using `parsnp` based on the complete genome sequences (999 genomes) resulting from the first filtering steps of `ViralClust`.

## Discussion

The rapid increase in publicly available viral genome sequences poses substantial challenges for downstream analyses, including multiple sequence alignment, phylogenetic inference, and comparative genomics. blackLarge and unevenly sampled datasets are often computationally infeasible to analyze in full - particularly for analyses such as multiple sequence alignment, phylogenetic inference, and comparative genomics - and are prone to biases introduced by outbreak-driven over-representation and technical inconsistencies. In this study, we address these challenges by presenting `ViralClust`, a modular pipeline designed to reduce large viral genome collections through clustering-based representative selection while preserving genetic diversity.

A key outcome of our analyses is that clustering-based reduction provides a robust alternative to random subsampling. Across six diverse RNA and DNA virus datasets, representative genome sets retained biologically meaningful diversity while reducing dataset size by 95 % on average. Importantly, clustering results were stable across taxonomic scales: representative sets generated at the family level still captured genotypes defined at the species level, as demonstrated for *Hepacivirus hominis* and members of the *Flaviviridae*. This observation indicates that representative selection does not necessarily obscure finer-scale evolutionary structure and can support comparative analyses across broad taxonomic ranges.

blackWhile the present evaluation focused on the preservation of phylogenetic topology and genetic diversity, the utility of representative selection extends to biologically relevant metadata such as host range, geographic distribution, temporal dynamics, and phenotypic characteristics. To the extent that such traits are reflected in genome sequence - as is the case, for example, for host-associated sequence signatures - clustering-based representative selection inherently captures this variation alongside genetic diversity. More broadly, the degree to which a representative set covers features of interest depends on the underlying research question, and we encourage users to evaluate their representative sets accordingly. As an example, we have previously shown in the context of Pestiviruses [50] that representatives selected by clustering adequately covered the host range present in the full dataset, supporting the biological relevance of the reduced sequence set beyond phylogenetic structure alone.

The evaluation further shows that clustering whole viral genomes is feasible across a wide range of genome sizes and architectures. Even large DNA virus genomes, such as those of *Monkeypox virus*, could be processed efficiently, demonstrating the scalability of the approach. In addition, dedicated pre-processing steps proved essential for obtaining interpretable clustering results.

These findings emphasize that representative selection is not solely dependent on the clustering algorithm itself but also on careful data preparation and evaluation.

blackWe note that recombination can influence clustering outcomes by generating conflicting similarity patterns across genomic regions, potentially leading to context-dependent results. This limitation is not specific to `ViralClust` but affects similarity- and distance-based clustering approaches in general. The extent of this effect depends on the biological question, taxonomic scale, and analyzed genomic regions. Therefore, for highly recombinant viruses, preprocessing steps such as recombination detection, masking, or restriction to conserved regions may be advisable prior to clustering, and clustering results should always be interpreted in their biological context.

Rather than identifying a single optimal clustering strategy, `ViralClust` integrates multiple clustering algorithms based on different underlying principles, including compactness and connectedness. This design reflects the absence of a universal ground truth for (viral) clustering and acknowledges that different biological questions may require different clustering assumptions. By enabling direct comparison of clustering outcomes within a unified framework, `ViralClust` supports informed methodological choices without prescribing a “best” solution. In summary, this work demonstrates that scalable, clustering-based representative selection can substantially reduce viral genome datasets while preserving relevant diversity and mitigating common sources of bias.

Through its modular design and integrated evaluation, `ViralClust` provides a flexible framework for virology research and serves as a reproducible starting point for downstream analyses of increasingly large viral genome collections. blackFor different biological questions other parameters of the integrated clustering tools might be tested further.

## Potential implications

Representative genome sets generated by `ViralClust` may facilitate a wide range of downstream analyses that are currently limited by dataset size or redundancy. For example, clustering-based representative selection can support the construction of high-quality multiple sequence alignments and enable RNA secondary structure prediction using representative alignments [51, 52, 50] as employed in Rfam [53], and large-scale analyses of endogenous viral elements [54, 55]. Similarly, workflows for virus–host prediction and comparative framework analyses, such as `VIDHOP` [56] and `POSEIDON` [57], may benefit from reduced yet diverse input datasets that remain biologically representative.

More broadly, whole-genome clustering may complement gene-centric approaches commonly used in viral comparative studies. While this work does not aim to redefine virus classification, our results suggest that genome-wide similarity patterns can provide additional context for studying viral diversity across taxonomic levels. We strongly advise that clustering-based representative selection be considered as a standard preprocessing step in future viral genome analyses, enabling more balanced, interpretable, and computationally tractable studies while remaining adaptable to specific research questions and analytical goals.

## Methods

### Tools integrated into ViralClust

We used existing tools for several steps in the `ViralClust` pipeline we designed.

#### Pre-processing

To remove 100 % redundant genomes in the input dataset, we apply the `easy-linclust` module of `MMSeqs2` (v14) [17, 18], with parameters `--min-seq-id 1.0`.

#### Clustering

Per default, `ViralClust` applies five different clustering methods: `CD-HIT-EST` (v4.8.1) [14], `SUMACLUST` (v1.0.31) [15], `VSEARCH` (v2.15) [16] from `vsearch`, `MMSeqs2` (v14) [17, 18], and `HDBSCAN` (v0.8.26) [23]. By default, `ViralClust` instructs `CD-HIT-EST`, `SUMACLUST`, `VSEARCH`, and `MMSeqs2` to cluster sequences with an identity of at least 90 % (default value of `CD-HIT-EST`). blackSince `MMSeqs2` offers a variety of clustering modules, we have integrated `easy-linclust` (default) and `easy-cluster`.

#### Connectedness Clustering

We have implemented a combination of dimension reduction and clustering in `ViralClust`. For this we used `PCA` or `UMAP` (`umap-learn` v0.4.4) [58], both implemented in `scikit-learn` (v1.2.2) [28] for dimension reduction and `HDBSCAN` for clustering [23]. Per default, we use `PCA` and reduce the dimension to either *d* = 50 or until the sum of the explained variance of the components is at least 0.7. When using `UMAP`, the dimensions are reduced to *d* = 50 with a minimum distance of 0.25, cosine distance as common usage in the context of embeddings and language [59], and random seed state of 42 as used in `UMAP`’s documentation to ensure reproducibility within this stochastic algorithm^3^. `HDBSCAN` can be seen as a hybrid approach of hierarchical-based (connectivity) and density-based (compactness) clustering approaches. For two vectors, `HDBSCAN` first calculates mutual reachability distance. Per default, we use the cosine distance implemented in `SciPy` (v1.4.1) [29] as metric. However, we allow a plethora of pre-defined distances, which can be used via the `--metric` parameter of our `HDBSCAN` module. We implemented the (standardized) Cosine (=default), Euclidean, Manhattan, Chebyshev, Minkowski, Canberra, Braycurtis, and Mahalanobis distance metrics via the `SciPy` library [29]. To accommodate the limitations of the current implementation of `HDBSCAN`, we have employed L2 normalization to preprocess the vectors. This ensures compatibility with distance calculations, as cosine distance is not supported. Subsequently, we use `HDBSCAN` with the Euclidean distance on these normalized vectors^4^.

### Visualization

Importantly, with `ViralClust`, our aim is not to construct a phylogeny of the viruses, but rather to ensure that the selected virus genomes are distributed evenly across the existing phylogenetic reconstruction, providing a rough overview of their diversity. Therefore, for the validation of our use cases, we additionally visualized the distribution of cluster representatives and NCBI RefSeq genomes across the phylogenetic representation with the Neighbornet algorithm implemented in `SplitsTree` (v4.19.2) [60, 61], generating split graphs, `FastTree` (v2.1.10) [32] as time-efficient tool for phylogenetic reconstruction of large datasets, and `parsnp` (v2.1.5) [62] for core-genome-based phylogenetic reconstruction (not part of the pipeline for the original data).

### Benchmarking and Scoring Metrics

blackAll `ViralClust` runs were conducted on a Linux-based x86_64 server equipped with four AMD Opteron 6376 processors comprising 64 logical CPU cores in total (8 cores per socket with 2 threads per core) and a maximum clock frequency of 2.3 GHz. The system architecture featured eight NUMA nodes and supported AMD-V virtualization. Computational tasks were executed in a shared-memory environment using the available multi-core CPU resources.

`ViralClust` offers a first evaluation based on several statistics using the parameter `--eval`. blackTo evaluate the performance of the five tools integrated into `ViralClust` across all six datasets presented here, we incorporated additional metrics determining the quality of the clustering output, summarized into one overall score. All scores are defined in the range [0, 1], where higher values indicate better performance, and the overall score has a maximum of 5.

black

#### Reduction Score

The Reduction Score (RS) measures how much a given tool reduces the input dataset through clustering, while accounting for sequences discarded by the clustering algorithm:

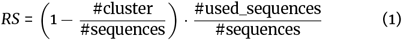

The first factor captures the proportion of sequences replaced by cluster representatives, while the second factor penalizes tools that discard a large fraction of input sequences rather than assigning them to clusters. A score of 1 indicates maximal reduction with no sequence loss.

black

#### Singleton Penalty Score

The Singleton Penalty Score (SPS) penalizes algorithms that leave many sequences unclustered, i.e. assigned to singleton clusters of size one:

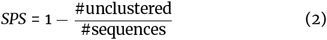

Singletons do not contribute to dataset reduction and may indicate over-fragmentation or, in the case of density-based methods such as `HDBSCAN`, sequences assigned to noise. A score of 1 indicates that no sequences were left unclustered.

black

#### Clustering Quality Score

The Clustering Quality Score (CQS) measures the separation between clusters relative to their internal cohesion, based on pairwise phylogenetic distances derived from a user-provided tree and the cluster assignment file of each tool:

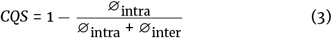

where ∅_intra_ and ∅_inter_ denote the average intra- and inter-cluster distances on the tree, respectively. A score close to 1 indicates well-separated clusters with low internal distances, whereas a score close to 0 indicates poor cluster separation. The CQS requires a Newick-formatted phylogenetic tree and the `.clstr` output file of each tool as input provided by `ViralClust`.

black

#### Taxonomic Concordance Score

The Taxonomic Concordance Score (TCS) measures how well the clustering reflects known taxonomy, based on NCBI taxonomic assignments of the input sequences:

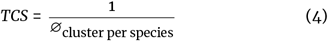

where ∅_cluster per species_ is the average number of clusters a single species is spread across. A score of 1 indicates that each species is entirely contained within a single cluster, with no fragmentation across cluster boundaries. Values below 1 indicate that species are split across multiple clusters. Note that TCS is sensitive to the completeness of NCBI taxonomic annotations and may be less reliable for novel or poorly characterized viruses.

black

#### Cluster-to-Taxonomy Ratio

The Cluster-to-Taxonomy Ratio (CTR) evaluates how closely the number of clusters produced by a tool matches the number of species defined by the ICTV:

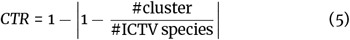

A score of 1 indicates that the number of clusters exactly matches the expected number of ICTV-defined species. Scores decrease symmetrically for both over- and under-clustering relative to this expectation, and are clamped to [0, 1].

black

#### Overall Score

The Overall Score (OS) aggregates all five metrics into a single summary value:

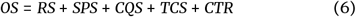

All five component scores contribute equally, each with a maximum of 1, giving an overall maximum of 5. A higher OS indicates a clustering result that simultaneously achieves strong dataset reduction, low singleton rate, good phylogenetic cluster separation, and concordance with known taxonomy at both the sequence annotation and ICTV species level.

## Availability of source code and requirements

`ViralClust` is available via GitHub: https://github.com/rnajena/viralclust.

## Data availability

The raw sequences and datasets supporting the results of this article are available in the zenodo repository: https://doi.org/10.5281/zenodo.18413373.

## Declarations

### Ethical Approval

Not applicable

### Consent for publication

Not applicable

### Author’s Contributions

Sandra Triebel: Data curation; Investigation; Software; Formal analysis; Validation; Visualization; Writing – review & editing. Tom Eulenfeld: Methodology; Software; Formal analysis. Kevin Lamkiewicz: Data curation; Investigation; Methodology; Software; Formal analysis; Validation; Visualization; Writing – original draft. Manja Marz: Supervision; Conceptualization; Validation; Writing – review & editing.

## Acknowledgements

We thank the Nextflow fairy Marie Lataretu! We further thank Franziska Hufsky for her `InkScape` magic.

## Footnotes

1 https://www.cdc.gov/flu/about/viruses-types.html Last Reviewed: 23.01.2026

2 as communicated in: https://github.com/scikit-learn-contrib/hdbscan/issues/69#issuecomment-317742362

3 www.umap-learn.readthedocs.io/en/latest/reproducibility.html

4 as communicated in: https://github.com/scikit-learn-contrib/hdbscan/issues/69#issuecomment-317742362

