## Supplementary Information for "Reducing haystacks to needles – ViralClust: A Nextflow pipeline to cluster viral sequences"

### `ViralClust` statistics

**Table S1.** Run time and memory usage of `ViralClust` steps measured by `Nextflow`. cd – `CD-HIT-EST`; sum – `SUMACLUST`; vse – `VSEARCH`; mmseq – `MMSeqs2`; hdb – `HDBSCAN`; #Sequences – number of input genomes/segments with accession date; only records labeled as `"complete"`; rem.-red. – remove redundancy; ∅ aln – average time/memory of alignment construction of representative genomes using `MAFFT`; ∅ tree – average time/memory of phylogenetic tree reconstruction of representative genomes using `FastTree`; eval – average time/memory of calculating basic cluster statistics; blackPeak RAM – peak RSS (resident set size) of `ViralClust` measured by `Nextflow`.

| Taxonomic clade | #Sequences | | Run time | | | | | | | | | |
|---|---|---|---|---|---|---|---|---|---|---|---|---|
| | | | Pre-processing | | Clustering | | | | | Evaluation | | |
| | 13.10.2022 | 17.11.2025 | sort | rem.-red. | cd | suma | vse | mmseq | hdb | ∅ aln | ∅ tree | eval |
| *Orthoebolavirus* | 632 | 691 | 13" | 7" | 20" | 31" | 4'21" | 1" | 27" | 58" | 3" | 13" |
| *Hepacivirus hominis* | 1,075 | 1,135 | 9" | 54" | 1'20" | 1'23" | 5'18" | 4" | 37" | 31" | 9" | 10" |
| *Orthoflavivirus* | 8,052 | 10,332 | 1'14" | 3' 4" | 2'29" | 21'13" | 30'45" | 5" | 13'36" | 1'21" | 24" | 11" |
| *Flaviviridae* | 9,895 | 12,676 | 1'35" | 4'28" | 11'36" | 44'35" | 55'53" | 10" | 22'54" | 3'12" | 1'41" | 12" |
| *Alphainfluenzavirus influenzae* (1) | 83,543 | 155,568 | 2'46" | 7'25" | 34" | 6'21" | 4'32" | 8" | 3h 00'19" | 7" | 33" | 15" |
| *Alphainfluenzavirus influenzae* (2) | 82,569 | 152,464 | 2'42" | 7' 2" | 35" | 5'55" | 4'48" | 8" | 2h 58'16" | 6" | 28" | 14" |
| *Alphainfluenzavirus influenzae* (3) | 83,400 | 154,548 | 2'39" | 6'12" | 42" | 5'16" | 4'29" | 8" | 2h 52'25" | 5" | 28" | 15" |
| *Alphainfluenzavirus influenzae* (4, H) | 156,586 | 238,758 | 3'38" | 4'58" | 1'30" | 36'39" | 4'38" | 27" | 7h 24'11" | 10" | 3'21" | 28" |
| *Alphainfluenzavirus influenzae* (5) | 84,653 | 158,491 | 2' 2" | 3'18" | 18" | 2'38" | 1'57" | 6" | 2h 14'38" | 3" | 16" | 13" |
| *Alphainfluenzavirus influenzae* (6, N) | 114,639 | 191,872 | 2'23" | 3'22" | 52" | 13'37" | 2'24" | 10" | 3h 41'39" | 4" | 35" | 16" |
| *Alphainfluenzavirus influenzae* (7) | 96,928 | 172,393 | 1'28" | 58" | 10" | 38" | 1' 4" | 4" | 1h 39'35" | 2" | 15" | 13" |
| *Alphainfluenzavirus influenzae* (8) | 85,944 | 160,084 | 1'10" | 49" | 11" | 39" | 49" | 3" | 1h 22'15" | 1" | 9" | 12" |
| *Monkeypox virus* | 1,077 | 4,070 | 9'50" | 3'15" | 1h 17'00" | 1h 38'54" | 18 h 9'10" | 2'45" | 33'32" | 1h 14'57" | 2'19" | 50" |

| Taxonomic clade | Memory usage | | | | | | | | | | |
|---|---|---|---|---|---|---|---|---|---|---|---|
| | Pre-processing | | Clustering | | | | | Evaluation | | | Peak RAM |
| | sort | rem.-red. | cd | suma | vse | mmseq | hdb | ∅ aln | ∅ tree | eval | |
| *Orthoebolavirus* | 13M | 58M | 169M | 21M | 31M | 3.5M | 335M | 1.8G | 88M | 6.2G | 11.4G |
| *Hepacivirus hominis* | 13M | 47M | 179M | 23M | 37M | 46M | 443M | 2.3G | 75M | 6.3G | 10.7G |
| *Orthoflavivirus* | 13M | 290M | 279M | 161M | 170M | 278M | 5.5G | 4.6G | 148M | 6.2G | 43.2G |
| *Flaviviridae* | 12M | 528M | 315M | 193M | 224M | 327M | 6.6G | 5.1G | 327M | 6.5G | 52.6G |
| *Alphainfluenzavirus influenzae* (1) | 12M | 796M | 293M | 286M | 600M | 619M | 3.3G | 316M | 78M | 6.2G | 273.9G |
| *Alphainfluenzavirus influenzae* (2) | 12M | 792M | 291M | 282M | 590M | 605M | 3.3G | 303M | 69M | 6.1G | 270.9G |
| *Alphainfluenzavirus influenzae* (3) | 12M | 775M | 283M | 270M | 558M | 586M | 3.3G | 217M | 73M | 6.0G | 268.8G |
| *Alphainfluenzavirus influenzae* (4, H) | 12M | 1.1G | 347M | 397M | 785M | 752M | 5.5G | 298M | 142M | 6.0G | 510.0G |
| *Alphainfluenzavirus influenzae* (5) | 12M | 600M | 236M | 183M | 339M | 268M | 3.0G | 205M | 48M | 6.3G | 238.3G |
| *Alphainfluenzavirus influenzae* (6, N) | 12M | 720M | 269M | 247M | 456M | 358M | 4.1G | 239M | 81M | 6.1G | 572.2G |
| *Alphainfluenzavirus influenzae* (7) | 12M | 347M | 206M | 126M | 258M | 17M | 2.7G | 142M | 31M | 6.0G | 225.1G |
| *Alphainfluenzavirus influenzae* (8) | 12M | 275M | 196M | 105M | 183M | 45M | 2.6G | 132M | 32M | 6.3G | 207.2G |
| *Monkeypox virus* | 20M | 1.2G | 2.8G | 259M | 619M | 1.2G | 4.5G | 260M | 1.3G | 6.4G | 6.5G |

### ICTV

**Table S2. "Ground truth" of virus phylogeny.** Number of genera, subgenera, species, subtypes, and/or genotypes according to ICTV, NCBI, and CDC. g – genera; sg – subgenera; s – species; st – subtypes; gt – genotypes; * – `https://www.cdc.gov/flu/about/viruses-types.html` Last Reviewed: 18.09.2024

| Taxonomic clade | # of | Source | Shown in | Link to ICTV |
|---|---|---|---|---|
| *Orthoebolavirus* | 6 s | ICTV [35] | Fig. S1 | *Orthoebolavirus* |
| *Hepacivirus hominis* | 8 gt | NCBI [38, 39] | Fig. S4 | *Hepacivirus* |
| *Orthoflavivirus* | 53 s | ICTV [36] | Fig. S4 | *Orthoflavivirus* |
| *Flaviviridae* | 4 g, 97 s | ICTV [36] | Fig. S4 | *Flaviviridae* |
| *Alphainfluenzavirus influenzae* | 137 st (> 130) | NCBI (CDC*) | Fig. S10 | *Orthomyxoviridae* |
| *Monkeypox virus* | 2 gt | NCBI [63] | Fig. S24 | *Orthopoxvirus* |

### Benchmark `ViralClust` by a variety of datasets

**Table S3.** Scoring metric sorted by dataset. RS – Reduction Score; SPS – Singleton Penalty Scor; CQS – Clustering Quality Score; TCS – Taxonomic Concordance Score (based on NCBI species); Overall Score – weighted sum of all scores above; IAV – *Alphainfluenzavirus influenzae*; cd – CD-HIT-EST; sum – SUMACLUST; vse – VSEARCH; mmseq – MMSeqs2; hdb – HDBSCAN;

| Taxonimic clade | Algorithm | RS | SPS | CQS | TCS | CTR | OS |
|---|---|---|---|---|---|---|---|
| | cd | 0.982 | 0.995 | 0.978 | 1.000 | 0.667 | 4.622 |
| | suma | 0.871 | 1.000 | 0.978 | 1.000 | 0.833 | 4.682 |
| *Orthoebolavirus* | vse | 0.984 | 1.000 | 0.977 | 0.893 | 0.833 | 4.687 |
| | mmseq | 0.975 | 0.991 | 0.987 | 0.820 | 0.167 | 3.940 |
| | hdb | 0.975 | 0.959 | 0.994 | 0.625 | 0.167 | 3.720 |
| | cd | 0.948 | 0.910 | 0.831 | 0.565 | 0.672 | 3.926 |
| | suma | 0.709 | 0.961 | 0.899 | 0.521 | 0.708 | 3.798 |
| *Hepacivirus hominis* | vse | 0.945 | 0.919 | 0.908 | 0.575 | 0.693 | 4.040 |
| | mmseq | 0.949 | 0.911 | 0.831 | 0.599 | 0.673 | 3.963 |
| | hdb | 0.979 | 0.878 | 0.835 | 0.633 | 0.673 | 3.998 |
| | cd | 0.991 | 0.989 | 0.959 | 0.694 | 0.698 | 4.331 |
| | suma | 0.894 | 0.989 | 0.961 | 0.637 | 0.660 | 4.141 |
| *Orthoflavivirus* | vse | 0.991 | 0.990 | 0.976 | 0.704 | 0.679 | 4.340 |
| | mmseq | 0.992 | 0.990 | 0.957 | 0.758 | 0.811 | 4.508 |
| | hdb | 0.987 | 0.910 | 0.901 | 0.402 | 0.094 | 3.294 |
| | cd | 0.977 | 0.964 | 0.965 | 0.610 | 0.000 | 3.516 |
| | suma | 0.862 | 0.972 | 0.967 | 0.578 | 0.144 | 3.523 |
| *Flaviviridae* | vse | 0.977 | 0.967 | 0.979 | 0.617 | 0.000 | 3.540 |
| | mmseq | 0.978 | 0.965 | 0.963 | 0.649 | 0.000 | 3.555 |
| | hdb | 0.980 | 0.859 | 0.705 | 0.478 | 0.000 | 3.022 |
| | cd | 0.999 | 0.999 | 0.550 | 0.990 | 0.884 | 4.422 |
| | suma | 0.929 | 0.994 | 0.956 | 0.990 | 0.983 | 4.852 |
| IAV (1) | vse | 0.998 | 1.000 | 0.956 | 0.990 | 0.986 | 4.930 |
| | mmseq | 0.995 | 0.982 | 0.619 | 0.943 | 0.885 | 4.424 |
| | hdb | 0.975 | 0.673 | 0.897 | 0.962 | 0.877 | 4.384 |
| | cd | 0.999 | 0.999 | 0.569 | 0.990 | 0.889 | 4.446 |
| | suma | 0.930 | 0.994 | 0.942 | 0.990 | 0.980 | 4.836 |
| IAV (2) | vse | 0.999 | 1.000 | 0.938 | 0.990 | 0.982 | 4.909 |
| | mmseq | 0.996 | 0.982 | 0.591 | 0.952 | 0.880 | 4.401 |
| | hdb | 0.977 | 0.689 | 0.903 | 0.962 | 0.883 | 4.414 |
| | cd | 0.999 | 0.999 | 0.529 | 0.990 | 0.879 | 4.396 |
| | suma | 0.933 | 0.994 | 0.934 | 0.990 | 0.978 | 4.829 |
| IAV (3) | vse | 0.999 | 1.000 | 0.940 | 0.990 | 0.982 | 4.911 |
| | mmseq | 0.995 | 0.982 | 0.625 | 1.000 | 0.900 | 4.502 |
| | hdb | 0.975 | 0.663 | 0.892 | 0.962 | 0.873 | 4.365 |
| | cd | 0.997 | 0.999 | 0.738 | 0.990 | 0.931 | 4.655 |
| | suma | 0.908 | 0.990 | 0.976 | 0.980 | 0.984 | 4.838 |
| IAV (4) | vse | 0.996 | 0.999 | 0.975 | 0.980 | 0.988 | 4.938 |
| | mmseq | 0.985 | 0.887 | 0.787 | 0.990 | 0.912 | 4.561 |
| | hdb | 0.979 | 0.689 | 0.940 | 0.971 | 0.895 | 4.474 |
| | cd | 0.999 | 0.999 | 0.520 | 0.990 | 0.877 | 4.385 |
| | suma | 0.952 | 0.995 | 0.951 | 0.990 | 0.983 | 4.871 |
| IAV (5) | vse | 0.998 | 1.000 | 0.952 | 0.990 | 0.985 | 4.925 |
| | mmseq | 0.996 | 0.988 | 0.586 | 0.962 | 0.883 | 4.415 |
| | hdb | 0.975 | 0.591 | 0.863 | 0.962 | 0.848 | 4.239 |
| | cd | 0.997 | 0.999 | 0.704 | 0.990 | 0.922 | 4.612 |
| | suma | 0.918 | 0.990 | 0.975 | 0.990 | 0.986 | 4.859 |
| IAV (6) | vse | 0.996 | 0.999 | 0.968 | 0.980 | 0.986 | 4.929 |
| | mmseq | 0.991 | 0.960 | 0.735 | 0.971 | 0.914 | 4.571 |
| | hdb | 0.978 | 0.652 | 0.907 | 0.971 | 0.877 | 4.385 |
| | cd | 0.999 | 1.000 | 0.637 | 1.000 | 0.909 | 4.545 |
| | suma | 0.955 | 0.996 | 0.800 | 0.990 | 0.946 | 4.687 |
| IAV (7) | vse | 0.999 | 1.000 | 0.829 | 1.000 | 0.957 | 4.785 |
| | mmseq | 0.994 | 0.978 | 0.782 | 1.000 | 0.939 | 4.693 |
| | hdb | 0.979 | 0.545 | 0.810 | 0.971 | 0.826 | 4.131 |
| | cd | 0.999 | 0.999 | 0.696 | 0.990 | 0.921 | 4.605 |
| | suma | 0.956 | 0.997 | 0.896 | 0.990 | 0.969 | 4.808 |
| IAV (8) | vse | 0.998 | 1.000 | 0.910 | 0.990 | 0.975 | 4.873 |
| | mmseq | 0.994 | 0.980 | 0.778 | 0.980 | 0.933 | 4.665 |
| | hdb | 0.981 | 0.583 | 0.863 | 0.971 | 0.849 | 4.247 |
| | cd | 0.986 | 0.990 | 0.514 | 0.200 | 0.000 | 2.690 |
| | suma | 0.337 | 0.999 | 0.475 | 0.000 | 0.500 | 2.311 |
| *Monkeypox virus* | vse | 0.997 | 0.999 | 0.998 | 0.667 | 0.500 | 4.161 |
| | mmseq | 0.978 | 0.931 | 0.604 | 0.250 | 0.000 | 2.763 |
| | hdb | 0.998 | 1.000 | 0.998 | 1.000 | 1.000 | 4.996 |

### *Orthoebolavirus*

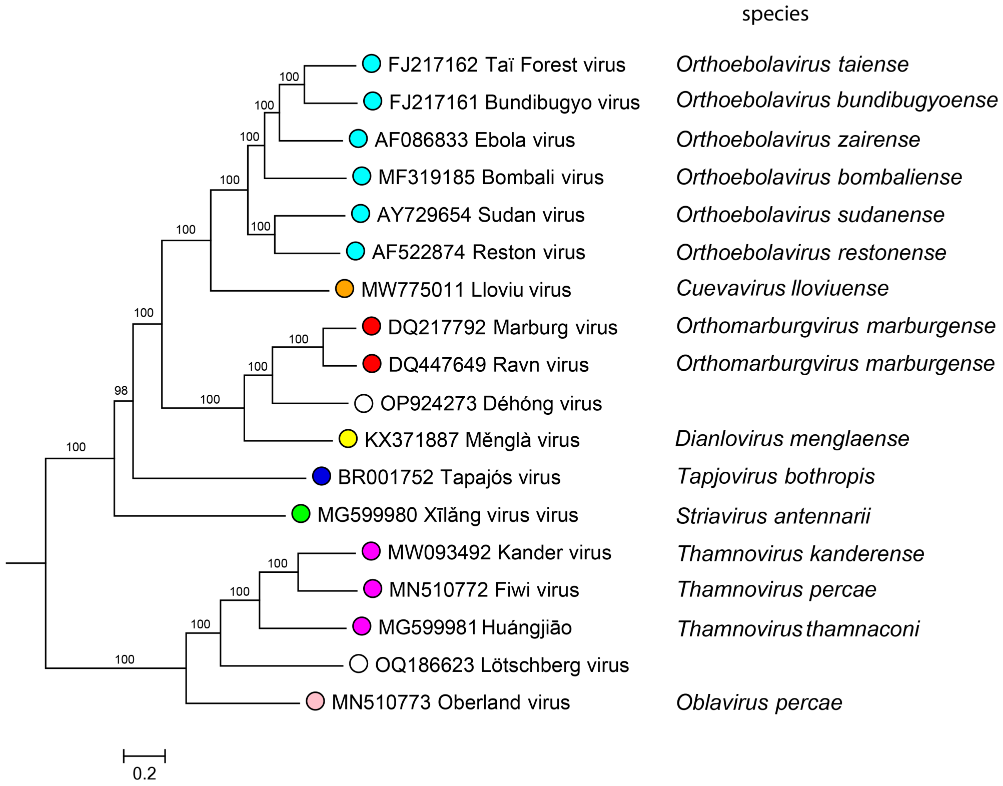


**Figure S1.** Phylogenetic tree of *Filoviridae* (ICTV) including *Orthoebolavirus* (cyan).

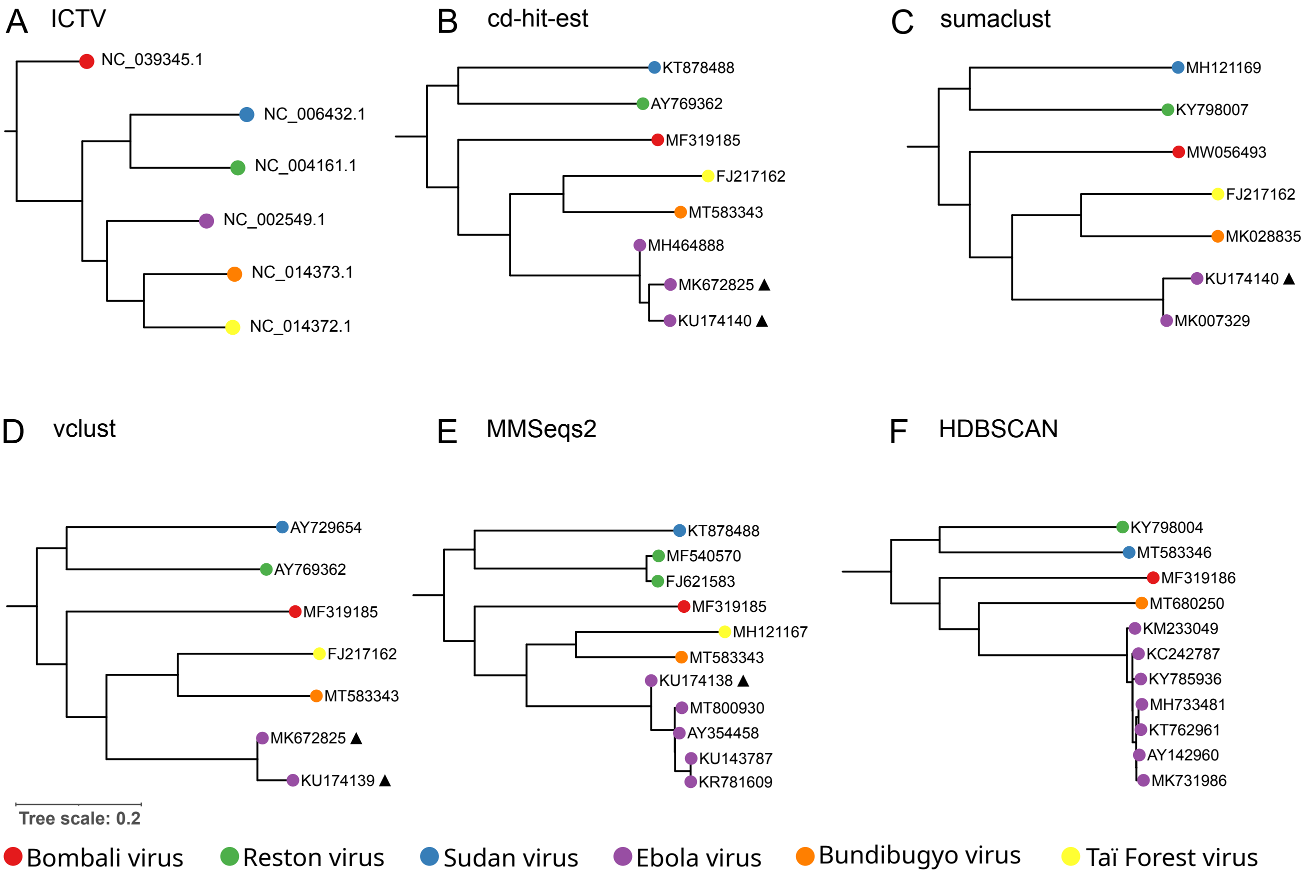


**Figure S2.** (A) Phylogenetic representation of ebolaviruses retrieved from the NCBI RefSeq database. The tree is based on the complete nucleotide sequence as described in the literature [64, 65]. (B–F) Phylogenetic representation of all representative ebolaviruses determined with `CD-HIT-EST`, `SUMACLUST`, `VSEARCH`, `MMSeqs2`, and `HDBSCAN`, respectively. Genomes derived from recombination or reverse genetics experiments are labeled with a triangle. Multiple sequence alignment and tree were calculated with `MAFFT` and `FastTree`, respectively.

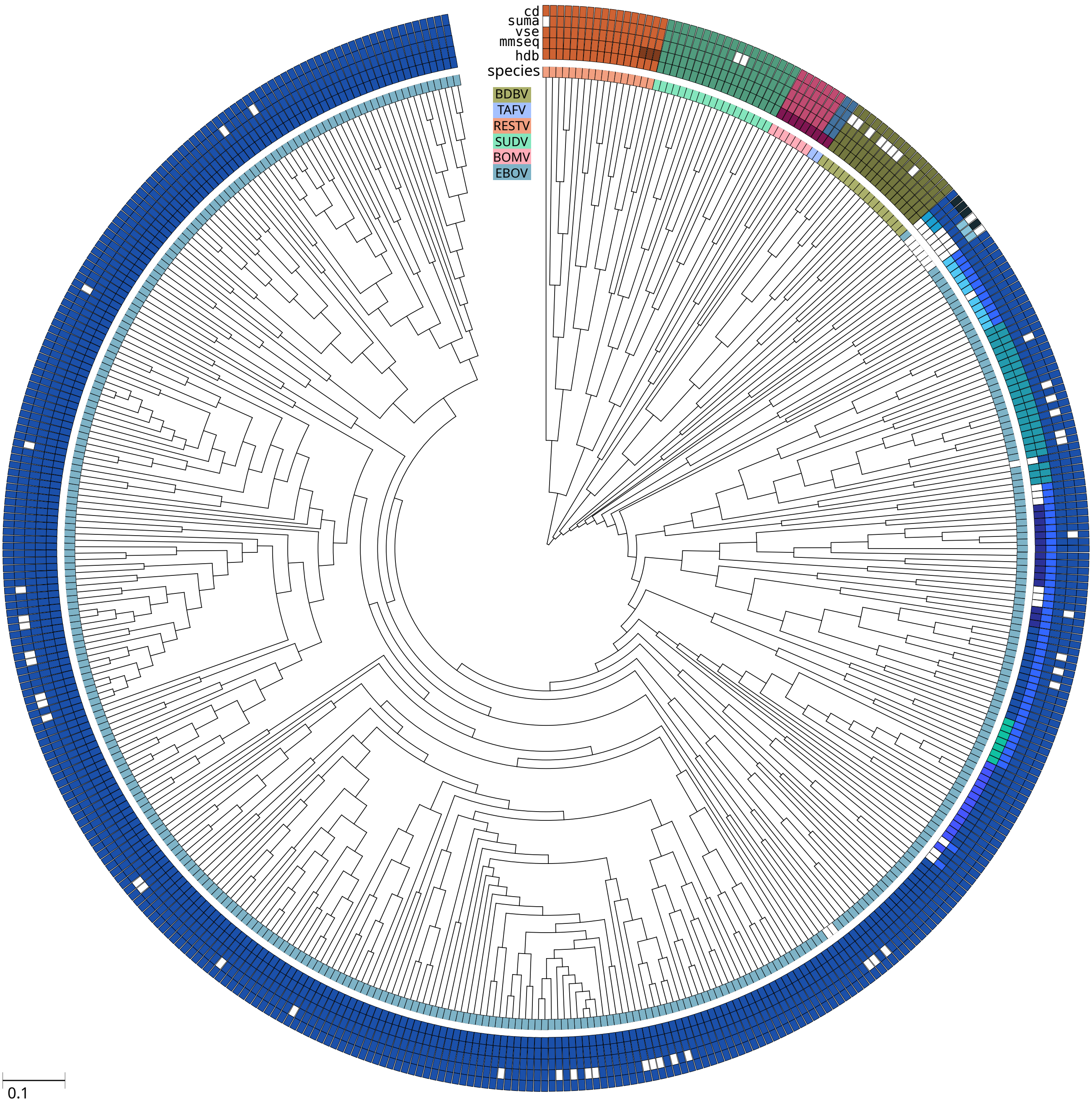


**Figure S3.** Phylogenetic representation of the *Orthoebolavirus* dataset. The tree is reconstructed using `RAxML-NG` based on the `MAFFT` alignment of the complete genome sequences (443 genomes) resulting from the first filtering steps of `ViralClust`. For detailed listing of parameters, we refer to the Methods section. The inner track around the tree represents the species information as retrieved from the NCBI database, serving as the 'ground truth'. Empty rectangles mark genomes for which no species information is available in the metadata. The top five tracks show the resulting clusters from `CD-HIT-EST`, `SUMACLUST`, `VSEARCH`, `MMSeqs2`, and `HDBSCAN`, respectively. Colors indicate the individual clusters identified by each tool and highlighting their level of agreement. We present singletons with empty rectangles.

### *Flaviviridae*

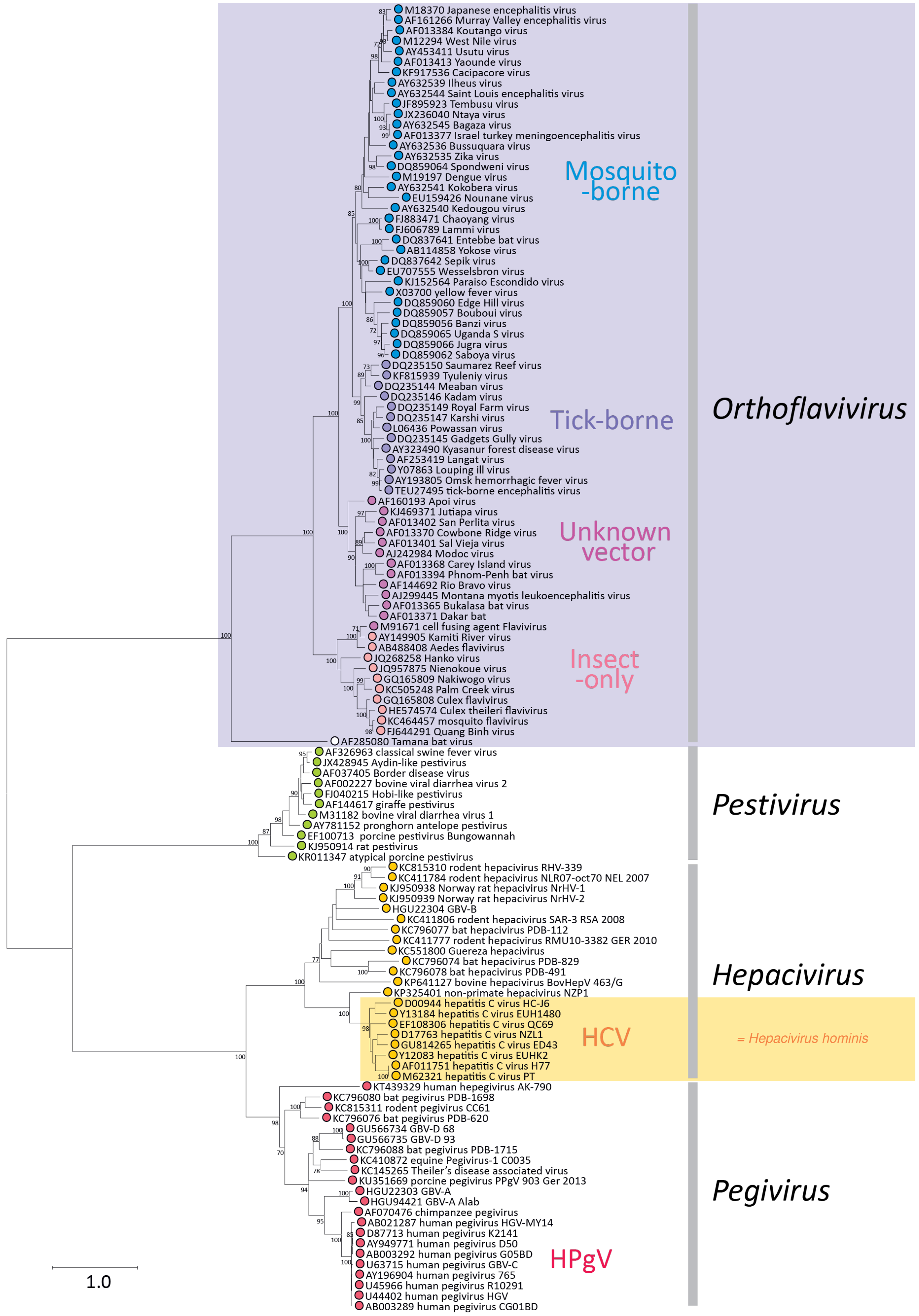


**Figure S4.** Phylogenetic tree of the family *Flaviviridae* (ICTV) including genus *Orthoflavivirus* (violet background) and species *Hepacivirus hominis* (HCV, yellow background). *Flaviviridae* consists of 4 genera including 97 species; *Orthoflavivirus* comprises 53 species; and *Hepacivirus hominis* includes 8 genotypes (only 7 genotypes are displayed in the tree). This tree is based on the partial gene sequences between positions 8,040–8,897 (numbered using positions in the HCV sequence, AF011751).

**Table S4.** Comparing the clustering results of the *Orthoflavivirus* dataset with the ICTV. ✓ – cluster with 1 ICTV representative; ∅ – number of clusters without ICTV representative; Problems – cluster with >= 2 ICTV representatives

Begin of Table

| | ✓ | ∅ | Discrepancies |
|---|---|---|---|
| cd | clstr 29 - Bagaza (AY632545)<br>clstr 30 - Dengue 2 (KF955373)<br>clstr 12 - Dengue 1 (NC_001477)<br>clstr 76 - Dengue 3 (M93130)<br>clstr 86 - Dengue 4 (AF326573)<br>clstr 64 - Yellow fever (X03700)<br>clstr 88 - Ilhéus (AY632539)<br>clstr 33 - Japanese encephalitis (M18370)<br>clstr 58 - Kokobera (AY632541)<br>clstr 85 - Kyasanur Forest disease (NC_039218)<br>clstr 43 - Langat (AF253419)<br>clstr 45 - St. Louis encephalitis (DQ525916)<br>clstr 56 - Louping ill (Y07863)<br>clstr 25 - Murray Valley encephalitis (AF161266)<br>clstr 21 - West Nile (M12294)<br>clstr 44 - Ntaya (JX236040)<br>clstr 19 - Omsk hemorrhagic fever (AY193805)<br>clstr 70 - Powassan (L06436)<br>clstr 24 - Tembusu (JF895923)<br>clstr 15 - Usutu (AY453411)<br>clstr 75 - Wesselsbron (EU707555)<br>clstr 77 - Zika (AY632535)<br>clstr 13 - Zika (KY766069, KJ776791, KX377337) | clstr 5, clstr 11, clstr 14,<br>clstr 16, clstr 17, clstr 20,<br>clstr 22, clstr 23, clstr 26,<br>clstr 27, clstr 28, clstr 31,<br>clstr 32, clstr 34, clstr 35,<br>clstr 37, clstr 40, clstr 46,<br>clstr 47, clstr 51, clstr 52,<br>clstr 54, clstr 55, clstr 57,<br>clstr 61, clstr 62, clstr 66,<br>clstr 67, clstr 68, clstr 69,<br>clstr 72, clstr 73, clstr 82,<br>clstr 87, clstr 95, clstr 99,<br>clstr 102, clstr 104, clstr 106,<br>clstr 107, clstr 112, clstr 113,<br>clstr 121, clstr 128, clstr 145 | clstr 143 - Saboya (NC_033697), Potiskum (NC_029054)<br>18 singletons: Aroa (AY632536)<br>Banzi (NC_043110)<br>Bouboui (NC_033693)<br>Cacipacoré (KF917536)<br>Edge Hill (NC_030289)<br>Gadgets Gully (NC_033723)<br>Jugra (NC_033699)<br>Jutiapa (KJ469371)<br>Kadam (NC_033724)<br>Kédougou (AY632540)<br>Meaban (NC_033721)<br>Modoc (NC_003635)<br>Montana myotis leukoencephalitis (NC_004119)<br>Royal Farm (NC_039219)<br>Saumarez Reef (NC_033726)<br>Tyuleniy (NC_023424)<br>Uganda S (NC_033698)<br>Yokose (AB114858) |
| suma | clstr 20 - Bagaza (AY632545)<br>clstr 0 - Dengue 2 (KF955373)<br>clstr 8 - Dengue 1 (NC_001477)<br>clstr 9 - Dengue 3 (M93130)<br>clstr 1 - Dengue 4 (AF326573)<br>clstr 14 - Yellow fever (X03700)<br>clstr 24 - Ilhéus (AY632539)<br>clstr 31 - Japanese encephalitis (M18370)<br>clstr 54 - Langat (AF253419)<br>clstr 47 - Louping ill (Y07863)<br>clstr 34 - Murray Valley encephalitis (AF161266)<br>clstr 2 - West Nile (M12294)<br>clstr 29 - Ntaya (JX236040)<br>clstr 50 - Omsk hemorrhagic fever (AY193805)<br>clstr 41 - Powassan (L06436)<br>clstr 26 - Tembusu (JF895923)<br>clstr 18 - Usutu (AY453411)<br>clstr 68 - Wesselsbron (EU707555)<br>clstr 84 - Zika (AY632535)<br>clstr 12 - Zika (KY766069, KJ776791, KX377337) | clstr 3, clstr 4, clstr 5,<br>clstr 6, clstr 7, clstr 10,<br>clstr 11, clstr 13, clstr 15,<br>clstr 16, clstr 17, clstr 21,<br>clstr 23, clstr 27, clstr 32,<br>clstr 33, clstr 35, clstr 39,<br>clstr 42, clstr 44, clstr 48,<br>clstr 49, clstr 51, clstr 52,<br>clstr 55, clstr 56, clstr 57,<br>clstr 58, clstr 59, clstr 63,<br>clstr 70, clstr 71, clstr 76,<br>clstr 80, clstr 81, clstr 83,<br>clstr 85, clstr 86, clstr 88,<br>clstr 89, clstr 92, clstr 93,<br>clstr 95, clstr 96, clstr 100,<br>clstr 118, clstr 120, clstr 128,<br>clstr 139, clstr 142 | clstr 114 - Saboya (NC_033697), Potiskum (NC_029054)<br>20 singletons: Aroa (AY632536)<br>Banzi (NC_043110)<br>Bouboui (NC_033693)<br>Cacipacoré (KF917536)<br>Edge Hill (NC_030289)<br>Gadgets Gully (NC_033723)<br>Jugra (NC_033699)<br>Jutiapa (KJ469371)<br>Kadam (NC_033724)<br>Kédougou (AY632540)<br>Kokobera (AY632541)<br>Kyasanur Forest disease (NC_039218)<br>Meaban (NC_033721)<br>Modoc (NC_003635)<br>Montana myotis leukoencephalitis (NC_004119)<br>Royal Farm (NC_039219)<br>Saumarez Reef (NC_033726)<br>Tyuleniy (NC_023424)<br>Uganda S (NC_033698)<br>Yokose (AB114858)<br>sequence of St. Louis encephalitis (DQ525916) removed by algorithm |
| vse | clstr 29 - Bagaza (AY632545)<br>clstr 31 - Dengue 2 (KF955373) | clstr 5, clstr 11, clstr 14,<br>clstr 16, clstr 17, clstr 20, | clstr 137 - Saboya (NC_033697), Potiskum (NC_029054)<br>18 singletons: Aroa (AY632536) |

Continuation of Tab. S4

| | | | |
|---|---|---|---|
| | clstr 12 - Dengue 1 (NC_001477) | clstr 22, clstr 23, clstr 26, | Banzi (NC_043110) |
| | clstr 76 - Dengue 3 (M93130) | clstr 27, clstr 28, clstr 32, | Bouboui (NC_033693) |
| | clstr 111 - Dengue 4 (AF326573) | clstr 34, clstr 35, clstr 37, | Cacipacoré (KF917536) |
| | clstr 64 - Yellow fever (X03700) | clstr 40, clstr 46, clstr 47, | Edge Hill (NC_030289) |
| | clstr 87 - Ilhéus (AY632539) | clstr 51, clstr 52, clstr 54, | Gadgets Gully (NC_033723) |
| | clstr 33 - Japanese encephalitis (M18370) | clstr 55, clstr 57, clstr 61, | Jugra (NC_033699) |
| | clstr 58 - Kokobera (AY632541) | clstr 62, clstr 66, clstr 67, | Jutiapa (KJ469371) |
| | clstr 84 - Kyasanur Forest disease (NC_039218) | clstr 68, clstr 69, clstr 72, | Kadam (NC_033724) |
| | clstr 43 - Langat (AF253419) | clstr 73, clstr 80, clstr 82, | Kédougou (AY632540) |
| | clstr 45 - St. Louis encephalitis (DQ525916) | clstr 85, clstr 86, clstr 97, | Meaban (NC_033721) |
| | clstr 56 - Louping ill (Y07863) | clstr 99, clstr 103, clstr 104, | Modoc (NC_003635) |
| | clstr 25 - Murray Valley encephalitis (AF161266) | clstr 105, clstr 110, clstr 113, | Montana myotis leukoencephalitis (NC_004119) |
| | clstr 21 - West Nile (M12294) | clstr 117, clstr 121, clstr 125, | Royal Farm (NC_039219) |
| | clstr 44 - Ntaya (JX236040) | clstr 141 | Saumarez Reef (NC_033726) |
| | clstr 19 - Omsk hemorrhagic fever (AY193805) | | Tyuleniy (NC_023424) |
| | clstr 70 - Powassan (L06436) | | Uganda S (NC_033698) |
| | clstr 24 - Tembusu (JF895923) | | Yokose (AB114858) |
| | clstr 15 - Usutu (AY453411) | | |
| | clstr 75 - Wesselsbron (EU707555) | | |
| | clstr 78 - Zika (AY632535) | | |
| | clstr 13 - Zika (KY766069, KJ776791, KX377337) | | |
| mmseq | clstr 11 - Bagaza (AY632545) | clstr 1, clstr 4, clstr 5, | clstr 40 - Saboya (NC_033697), Potiskum (NC_029054) |
| | clstr 121 - Dengue 2 (KF955373) | clstr 16, clstr 22, clstr 30, | 18 singletons: Aroa (AY632536) |
| | clstr 51 - Dengue 1 (NC_001477) | clstr 34, clstr 37, clstr 41, | Banzi (NC_043110) |
| | clstr 101 - Dengue 3 (M93130) | clstr 42, clstr 43, clstr 44, | Bouboui (NC_033693) |
| | clstr 12 - Dengue 4 (AF326573) | clstr 45, clstr 46, clstr 49, | Cacipacoré (KF917536) |
| | clstr 118 - Yellow fever (X03700) | clstr 52, clstr 53, clstr 54, | Edge Hill (NC_030289) |
| | clstr 110 - Ilhéus (AY632539) | clstr 58, clstr 62, clstr 63, | Gadgets Gully (NC_033723) |
| | clstr 108 - Japanese encephalitis (M18370) | clstr 71, clstr 84, clstr 86, | Jugra (NC_033699) |
| | clstr 83 - Kokobera (AY632541) | clstr 88, clstr 92, clstr 95, | Jutiapa (KJ469371) |
| | clstr 50 - Kyasanur Forest disease (NC_039218) | clstr 98, clstr 103, clstr 104, | Kadam (NC_033724) |
| | clstr 120 - Langat (AF253419) | clstr 107, clstr 109, clstr 112, | Kédougou (AY632540) |
| | clstr 128 - St. Louis encephalitis (DQ525916) | clstr 124, clstr 126, clstr 127, | Meaban (NC_033721) |
| | clstr 119 - Louping ill (Y07863) | clstr 132, clstr 133, clstr 135 | Modoc (NC_003635) |
| | clstr 137 - Murray Valley encephalitis (AF161266) | | Montana myotis leukoencephalitis (NC_004119) |
| | clstr 13 - West Nile (M12294) | | Royal Farm (NC_039219) |
| | clstr 91 - Ntaya (JX236040) | | Saumarez Reef (NC_033726) |
| | clstr 10 - Omsk hemorrhagic fever (AY193805) | | Tyuleniy (NC_023424) |
| | clstr 116 - Powassan (L06436) | | Uganda S (NC_033698) |
| | clstr 73 - Tembusu (JF895923) | | Yokose (AB114858) |
| | clstr 31 - Usutu (AY453411) | | |
| | clstr 117 - Wesselsbron (EU707555) | | |
| | clstr 139 - Zika (AY632535) | | |
| | clstr 20 - Zika (KY766069, KJ776791, KX377337) | | |
| hdb | clstr 77 - Bagaza (AY632545) | clstr 0, clstr 1, clstr 3, clstr 4, | clstr 59 - Modoc (NC_003635), Saboya (NC_033697), |
| | clstr 71 - Dengue 2 (KF955373) | clstr 5, clstr 6, clstr 7, clstr 10, | Uganda S (NC_033698), Wesselsbron (EU707555) |
| | clstr 53 - Dengue 1 (NC_001477) | clstr 11, clstr 12, clstr 13, clstr 14, | 23 singletons: Aroa (AY632536) |
| | clstr 23 - Dengue 3 (M93130) | clstr 15, clstr 16, clstr 17, clstr 18, | Banzi (NC_043110) |
| | clstr 20 - Dengue 4 (AF326573) | clstr 19, clstr 21, clstr 22, clstr 24, | Bouboui (NC_033693) |

Continuation of Tab. S4

| | | |
|---|---|---|
| clstr 45 - Yellow fever (X03700) | clstr 25, clstr 26, clstr 27, clstr 28, | Cacipacoré (KF917536) |
| clstr 83 - Kyasanur Forest disease (NC_039218) | clstr 30, clstr 31, clstr 32, clstr 33, | Edge Hill (NC_030289) |
| clstr 78 - St. Louis encephalitis (DQ525916) | clstr 34, clstr 35, clstr 36, clstr 37, | Gadgets Gully (NC_033723) |
| clstr 61 - Murray Valley encephalitis (AF161266) | clstr 38, clstr 39, clstr 40, clstr 41, | Ilhéus (AY632539) |
| clstr 43 - West Nile (M12294) | clstr 42, clstr 44, clstr 46, clstr 47, | Japanese encephalitis (M18370) |
| clstr 85 - Omsk hemorrhagic fever (AY193805) | clstr 48, clstr 49, clstr 50, clstr 51, | Jugra (NC_033699) |
| clstr 29 - Powassan (L06436) | clstr 52, clstr 54, clstr 55, clstr 56, | Jutiapa (KJ469371) |
| clstr 76 - Tembusu (JF895923) | clstr 57, clstr 58, clstr 60, clstr 62, | Kadam(NC_033724) |
| clstr 9 - Usutu (AY453411) | clstr 63, clstr 64, clstr 65, clstr 66, | Kédougou (AY632540) |
| clstr 8 - Zika (AY632535) | clstr 67, clstr 68, clstr 69, clstr 70, | Kokobera (AY632541) |
| clstr 2 - Zika (KY766069, KJ776791, KX377337) | clstr 72, clstr 73, clstr 74, clstr 75, | Langat (AF253419) |
| | clstr 79, clstr 80, clstr 81, clstr 82, | Louping ill (Y07863) |
| | clstr 84, clstr 86, clstr 87, clstr 88, | Meaban (NC_033721) |
| | clstr 89, clstr 90, clstr 91, clstr 92, | Montana myotis leukoencephalitis (NC_004119) |
| | clstr 93, clstr 94, clstr 95, clstr 96, | Ntaya (JX236040) |
| | clstr 97, clstr 98, clstr 99, clstr 100 | Royal Farm (NC_039219) |
| | Potiskum (NC_029054) | |
| | Saumarez Reef (NC_033726) | |
| | Tyuleniy (NC_023424) | |
| | Yokose (AB114858) | |

End of Table

**Table S5.** *Orthoflavivirus* species, their ICTV representatives & NCBI RefSeqs and the cluster number in which they were clustered. GenBank – GenBank accession ID; NCBI RefSeq – NCBI RefSeq accession ID; ICTV – category in the ICTV (E – exemplar genome, A – additional genome); Nucl Comp – Nucleotide completeness (C – Complete, CC – Coding-complete P – Partial); dataset – presence of the genome in our *Orthoflavivirus* dataset (G – GenBank in our dataset, N – NCBI RefSeq in our dataset); cd – CD-HIT-EST; sum – SUMACLUST; vse – VSEARCH; mmseq – MMSeqs2; hdb – HDBSCAN;

Begin of Table

| Virus Name | Abbr | Accession ID | | ICTV | Nucl Comp | dataset | Cluster Nr. | | | | |
|---|---|---|---|---|---|---|---|---|---|---|---|
| | | GenBank | NCBI RefSeq | | | | cd | suma | vse | mmseq | hdb |
| Apoi virus | APOIV | AF160193 | NC_003676 | E | C | | | | | | |
| Aroa virus | AROAV | AY632536 | NC_009026 | E | C | G | 74 | 69 | 74 | 61 | -1 |
| Aroa virus | AROAV | AF013362 | | A | P | | | | | | |
| Bussuquara virus | BSQV | AF013366 | | A | P | | | | | | |
| Iguape virus | IGUV | AF013375 | | A | P | | | | | | |
| Naranjal virus | NJLV | AF013390 | | A | P | | | | | | |
| Bagaza virus | BAGV | AY632545 | NC_012534 | E | C | G | 29 | 20 | 29 | 11 | 77 |
| Banzi virus | BANV | DQ859056 | NC_043110 | E | C | N | 140 | 113 | 136 | 74 | -1 |
| Bouboui virus | BOUV | DQ859057 | NC_033693 | E | C | N | 142 | 112 | 138 | 7 | -1 |
| Rio Bravo virus | RBV | AF144692 | NC_003675 | E | C | | | | | | |
| Bukalasa bat virus | BBV | AF013365 | NC_043111 | E | P | | | | | | |
| Cacipacoré virus | CPCV | KF917536 | NC_026623 | E | C | G | 131 | 117 | 126 | 76 | -1 |
| Carey Island virus | CIV | AF013368 | NC_043112 | E | P | | | | | | |
| Cowbone Ridge virus | CRV | AF013370 | NC_043113 | E | P | | | | | | |
| Dakar bat virus | DBV | AF013371 | NC_043114 | E | P | | | | | | |
| Dengue virus type 2 | DENV-2 | U87411 (KF955373) | NC_001474 | E | C | * | 30 | 0 | 31 | 121 | 71 |
| Dengue virus type 1 | DENV-1 | U88536 | NC_001477 | A | C | N | 12 | 8 | 12 | 51 | 53 |
| Dengue virus type 3 | DENV-3 | M93130 | | A | C | G | 76 | 9 | 76 | 101 | 23 |
| Dengue virus type 4 | DENV-4 | AF326573 | | A | C | G | 86 | 1 | 111 | 12 | 20 |
| Edge Hill virus | EHV | DQ859060 | NC_030289 | E | C | N | 139 | 102 | 134 | 111 | -1 |
| Tick-borne encephalitis virus - European | TBEV-Eur | U27495 | NC_001672 | E | C | | | | | | |
| Tick-borne encephalitis virus - Far Eastern | TBEV-FE | X07755 | | A | P | | | | | | |
| Tick-borne encephalitis virus - Siberian | TBEV-Sib | L40361 | | A | C | 2024 | | | | | |

Continuation of Tab. S5

| | | | | | | | | | | | |
|---|---|---|---|---|---|---|---|---|---|---|---|
| Entebbe bat virus | ENTV | DQ837641 | NC_008718 | E | C | | | | | | |
| Sokuluk virus | SOKV | AF013405 | | A | P | | | | | | |
| Yellow fever virus | YFV | X03700 | NC_002031 | E | C | G | 64 | 14 | 64 | 118 | 45 |
| Gadgets Gully virus | GGYV | DQ235145 | NC_033723 | E | C | N | 135 | 99 | 130 | 21 | -1 |
| Ilhéus virus | ILHV | AY632539 | NC_009028 | E | C | G | 88 | 24 | 87 | 110 | -1 |
| Rocio virus | ROCV | AF013397 | | A | P | | | | | | |
| Israel turkey meningoencephalomyelitis virus | ITV | AF013377 | NC_043115 | E | P | | | | | | |
| Japanese encephalitis virus | JEV | M18370 | NC_001437 | E | C | G | 33 | 31 | 33 | 108 | -1 |
| Jugra virus | JUGV | DQ859066 | NC_033699 | E | C | N | 144 | 109 | 139 | 125 | -1 |
| Jutiapa virus | JUTV | KJ469371 | NC_026620 | E | C | G | 146 | 105 | 142 | 105 | -1 |
| Kadam virus | KADV | DQ235146 | NC_033724 | E | C | N | 138 | 101 | 133 | 60 | -1 |
| Kédougou virus | KEDV | AY632540 | NC_012533 | E | C | G | 101 | 22 | 101 | 35 | -1 |
| Kokobera virus | KOKV | AY632541 | NC_009029 | E | C | G | 58 | 37 | 58 | 83 | -1 |
| Stratford virus | STRV | AF013407 | | A | P | | | | | | |
| Koutango virus | KOUV | AF013384 | NC_043116 | E | P | | | | | | |
| Kyasanur Forest disease virus | KFDV | AY323490 | NC_039218 | E | C | N | 85 | 106 | 84 | 50 | 83 |
| Alkhumra hemorrhagic fever virus | AHFV | AF331718 | | A | C | | | | | | |
| Langat virus | LGTV | AF253419 | NC_003690 | E | C | G | 43 | 54 | 43 | 120 | -1 |
| St. Louis encephalitis virus | SLEV | DQ525916 | NC_007580 | E | C | G | 45 | | 45 | 128 | 78 |
| Louping ill virus | LIV | Y07863 | NC_001809 | E | C | G | 56 | 47 | 56 | 119 | -1 |
| Louping ill virus - British | GGEV | D12937 | | A | P | | | | | | |
| Louping ill virus - Irish | LIV-Brit | X86784 | | A | P | | | | | | |
| Louping ill virus - Spanish | LIV-Ir | DQ235152 | | A | CC | | | | | | |
| Greek goat encephalitits virus | LIV-Spain | DQ235153 | | A | CC | | | | | | |
| Turkish sheep encephalitis virus | TSEV | DQ235151 | | A | CC | | | | | | |
| Meaban virus | MEAV | DQ235144 | NC_033721 | E | C | N | 133 | 103 | 128 | 6 | -1 |
| Modoc virus | MODV | AJ242984 | NC_003635 | E | C | N | 120 | 73 | 116 | 66 | 59 |
| Montana myotis leukoencephalitis virus | MMLV | AJ299445 | NC_004119 | E | C | N | 108 | 74 | 106 | 32 | -1 |
| Murray Valley encephalitis virus | MVEV | AF161266 | NC_000943 | E | C | G | 25 | 34 | 25 | 137 | 61 |
| Alfuy virus | ALFV | AF013360 | | A | P | | | | | | |
| West Nile virus | WNV | M12294 | NC_001563 | E | C | G | 21 | 2 | 21 | 13 | 43 |
| Kunjin virus | KUNV | D00246 | | A | C | | | | | | |
| Ntaya virus | NTAV | JX236040 | NC_018705 | E | C | G | 44 | 29 | 44 | 91 | -1 |
| Omsk hemorrhagic fever virus | OHFV | AY193805 | NC_005062 | E | C | G | 19 | 50 | 19 | 10 | 85 |
| San Perlita virus | SPV | AF013402 | NC_043118 | E | P | | | | | | |
| Phnom Penh bat virus | PPBV | AF013394 | NC_074777 | E | P | | | | | | |
| Batu Cave virus | BCV | AF013369 | | A | P | | | | | | |
| Powassan virus | POWV | L06436 | NC_003687 | E | C | G | 70 | 41 | 70 | 116 | 29 |
| Deer tick virus | DTV | AF311056 | | A | CC | | | | | | |
| Royal Farm virus | RFV | DQ235149 | NC_039219 | E | C | N | 134 | 108 | 129 | 122 | -1 |
| Saboya virus | SABV | DQ859062 | NC_033697 | E | C | N | 143 | 114 | 137 | 40 | 59 |
| Potiskum virus | POTV | DQ859067 | NC_029054 | A | CC | N | 143 | 114 | 137 | 40 | -1 |
| Saumarez Reef virus | SREV | DQ235150 | NC_033726 | E | C | N | 132 | 104 | 127 | 15 | -1 |
| Sepik virus | SEPV | DQ837642 | NC_008719 | E | C | | | | | | |
| Tembusu virus | TMUV | JF895923 | NC_015843 | E | C | G | 24 | 26 | 24 | 73 | 76 |
| Tyuleniy virus | TYUV | KF815939 | NC_023424 | E | C | N | 111 | 140 | 109 | 130 | -1 |
| Uganda S virus | UGSV | DQ859065 | NC_033698 | E | C | N | 141 | 111 | 135 | 64 | 59 |
| Usutu virus | USUV | AY453411 | NC_006551 | E | C | G | 15 | 18 | 15 | 31 | 9 |
| Sal Vieja virus | SVV | AF013401 | NC_043117 | E | P | | | | | | |
| Wesselsbron virus | WESSV | EU707555 | NC_012735 | E | C | G | 75 | 68 | 75 | 117 | 59 |
| Yaoundé virus | YAOV | AF013413 | NC_074778 | E | P | | | | | | |
| Yokose virus | YOKV | AB114858 | NC_005039 | E | C | G | 65 | 61 | 65 | 113 | -1 |
| Zika virus | ZIKV | AY632535 | NC_012532 | E | C | G | 77 | 84 | 78 | 139 | 8 |
| Zika virus | ZIKV | KY766069 | | A | C | G | 13 | 12 | 13 | 20 | 2 |
| Zika virus | ZIKV | KJ776791 | | A | C | G | 13 | 12 | 13 | 20 | 2 |
| Zika virus | ZIKV | KX377337 | | A | C | G | 13 | 12 | 13 | 20 | 2 |

End of Table

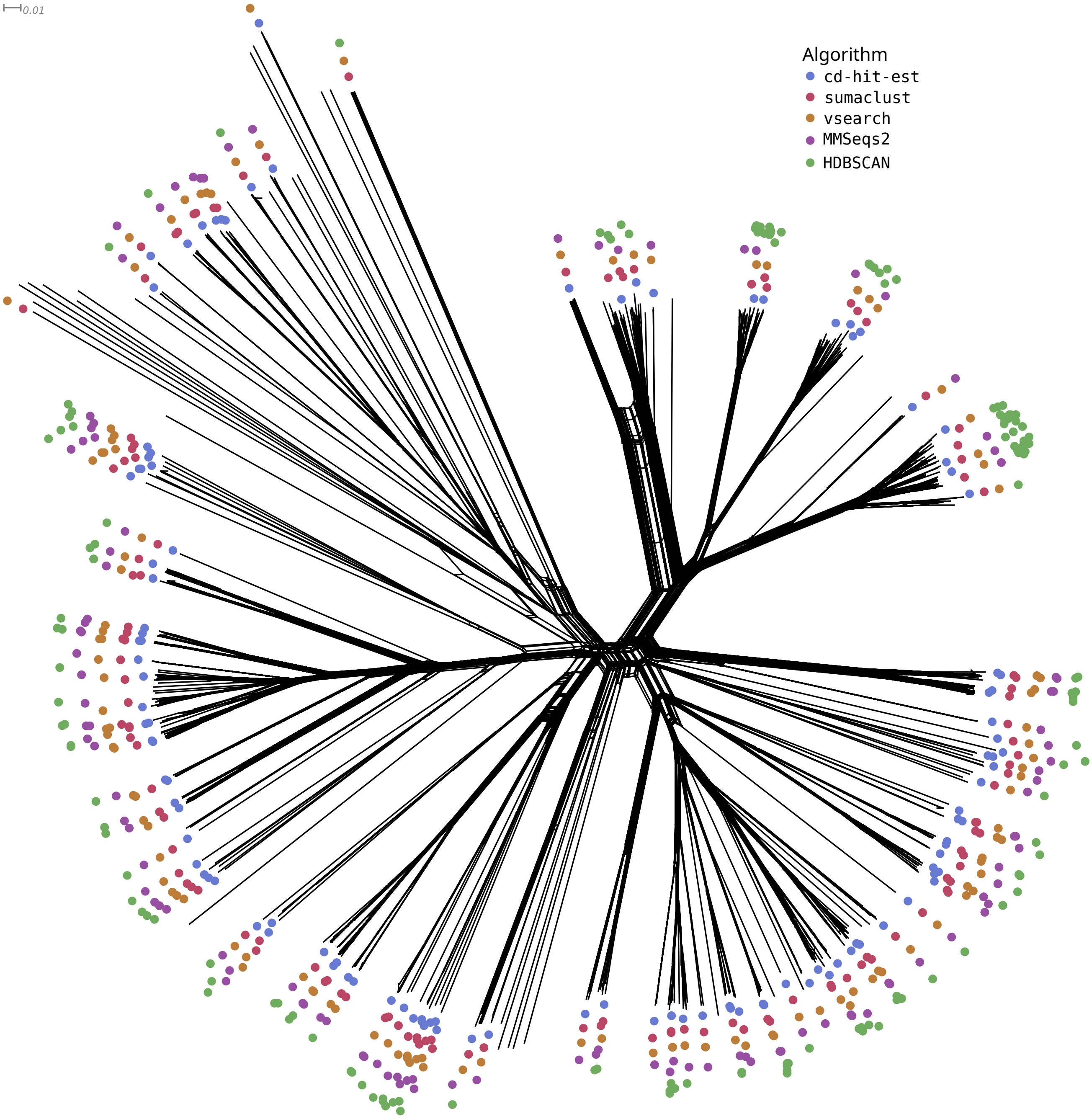


**Figure S5.** Split graph of *Orthoflavivirus* with highlighted cluster representatives of CD-HIT-EST, SUMACLUST, VSEARCH, MMSeqs2, and HDBSCAN. The split graph is reconstructed using SplitsTree based on the MAFFT alignment of the complete genome sequences (7,681 genomes) resulting from the first filtering steps of ViralClust.

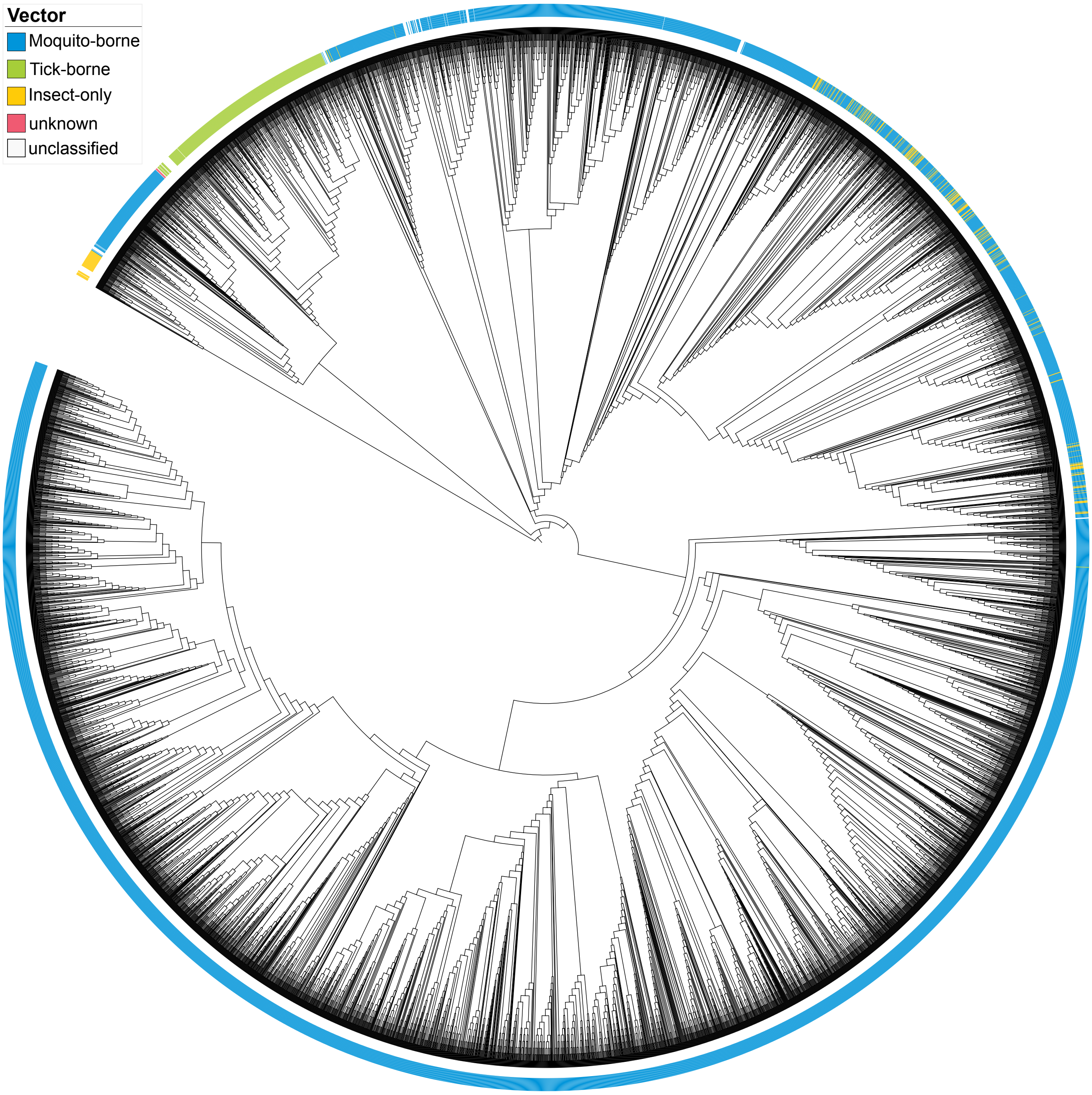


**Figure S6.** Phylogenetic representation of *Orthoflavivirus* showing vectors (Mosquito-borne, Tick-borne, Insect-only, and unknown). The tree is reconstructed using `FastTree` based on the `MAFFT` alignment of the complete genome sequences (7,681 genomes) resulting from the first filtering steps of `ViralClust`. More details in Fig. 5.

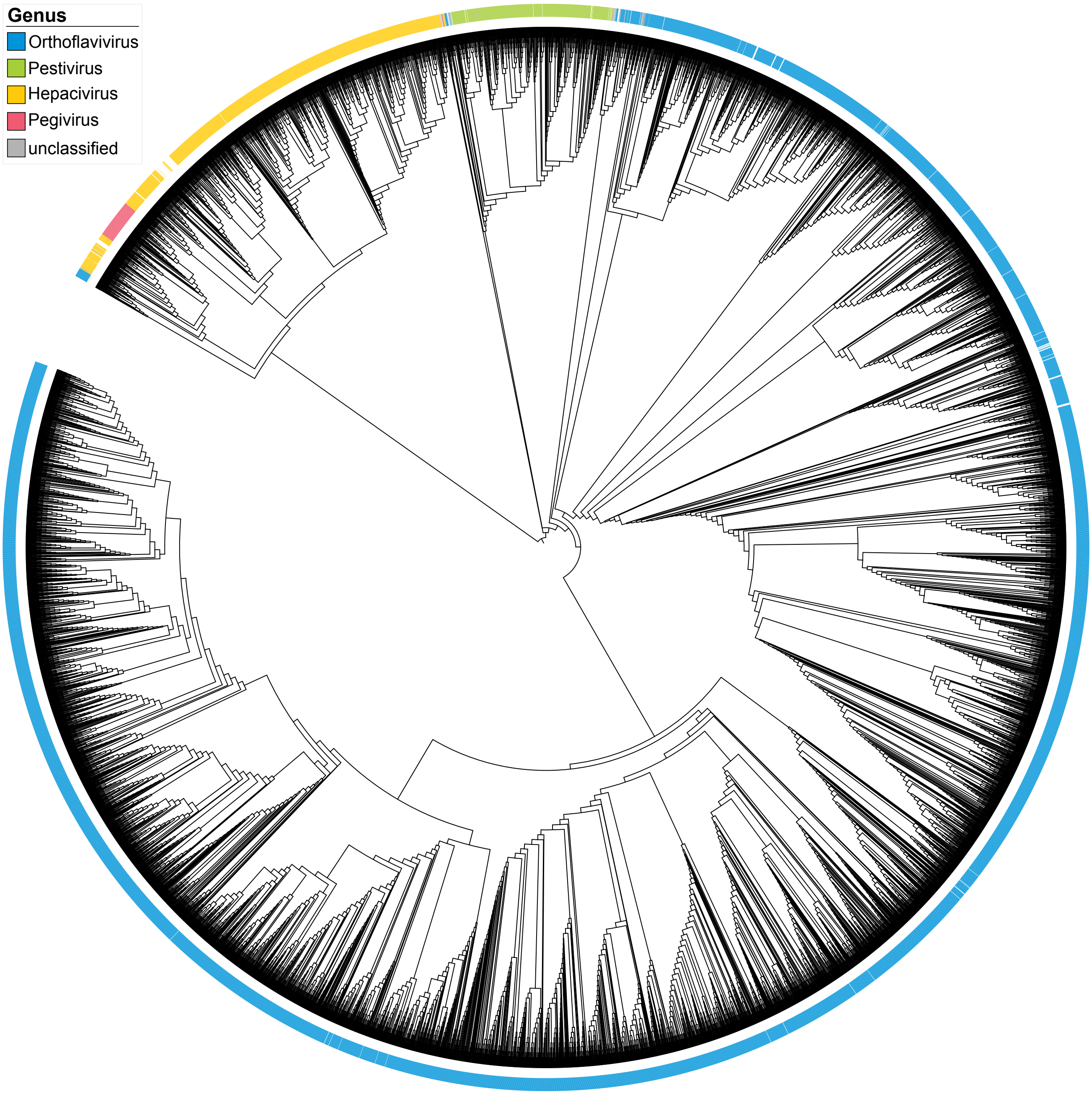


**Figure S7.** Phylogenetic representation of *Flaviviridae* with labeled genera: *Orthoflavivirus*, *Pestivirus*, *Hepacivirus*, and *Pegivirus*. The tree is reconstructed using `FastTree` based on the `MAFFT` alignment of the complete genome sequences (9,478 genomes) resulting from the first filtering steps of `ViralClust`. More details in Fig. 6.

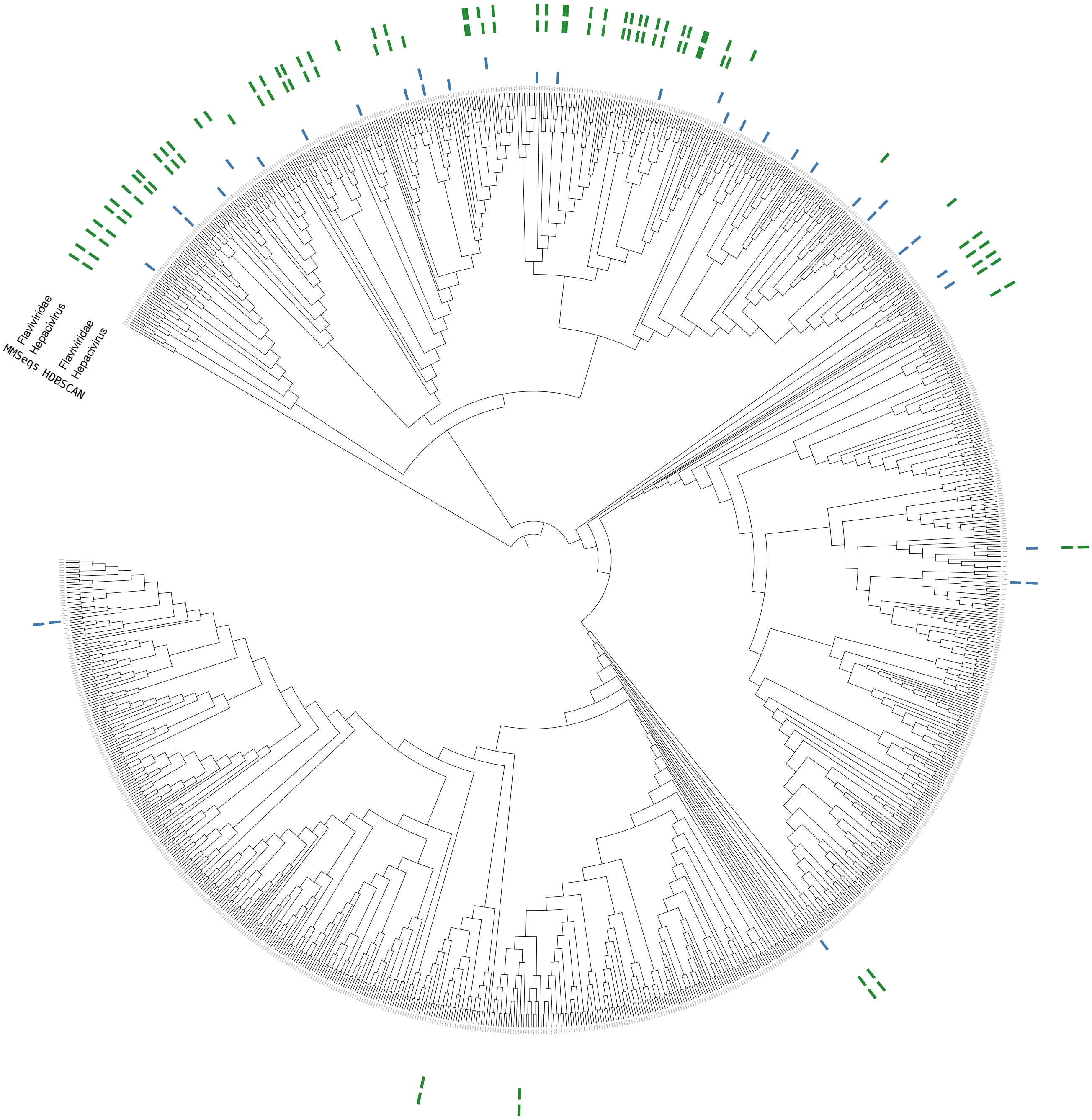


**Figure S8.** Phylogenetic representation of HCV showing the cluster representatives of `MMSeqs2` and `HDBSCAN` since the results show differences in the clustering of HCV only and HCV clustered together with all other *Flaviviridae*. The tree is reconstructed using `FastTree` based on the `MAFFT` alignment of the complete genome sequences (1,071 genomes) resulting from the first filtering steps of `ViralClust`.

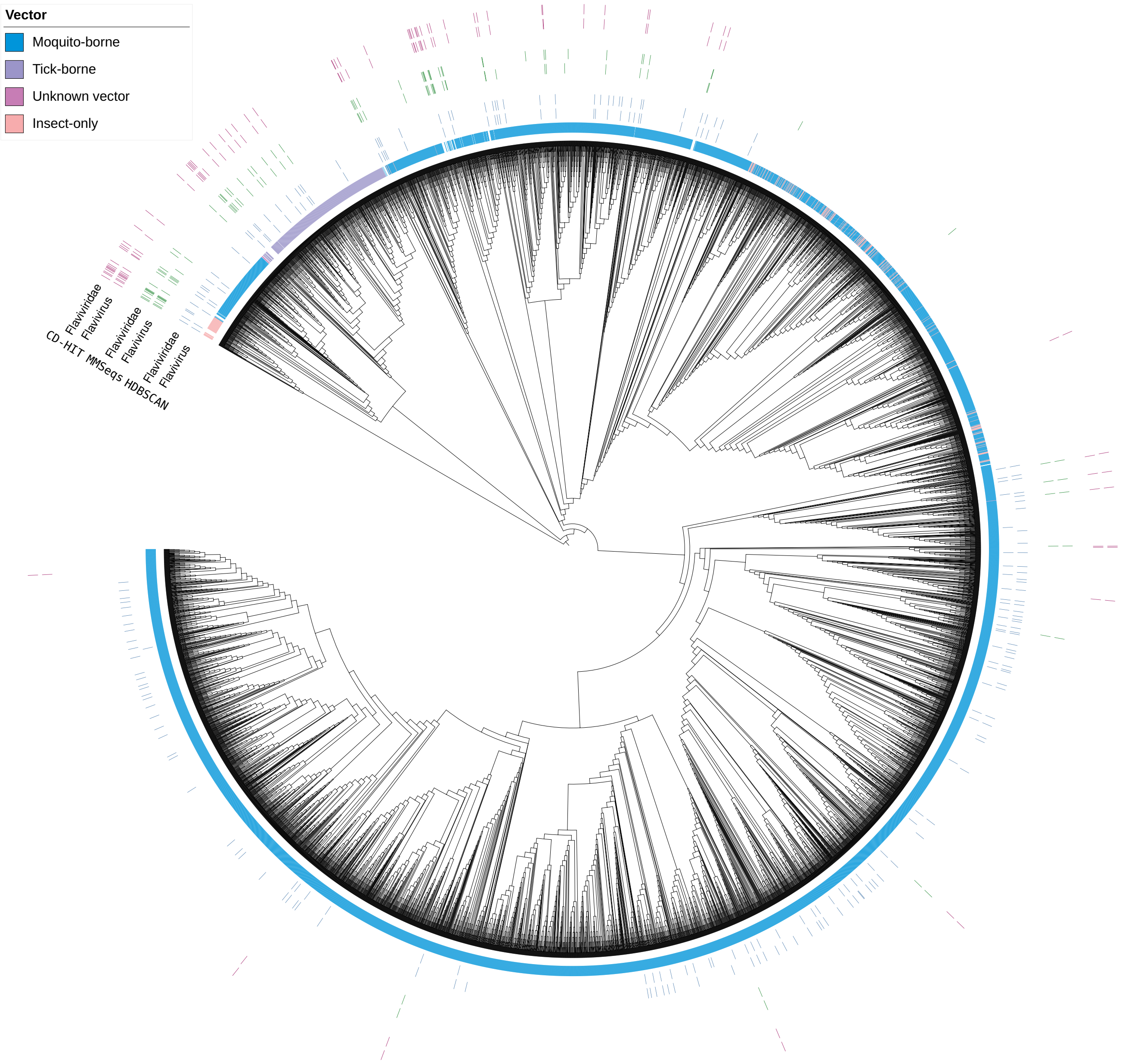


**Figure S9.** Phylogenetic representation of *Orthoflavivirus* showing the cluster representatives and labeled vectors (Mosquito-borne, Tick-borne, Unknown vector, and Insect-only). The clustering of the *Orthoflavivirus* only and the *Orthoflavivirus* among the entire *Flaviviridae* dataset showed differences in the results of `CD-HIT-EST`, `MMSeqs2`, and `HDBSCAN`. The tree is reconstructed using `FastTree` based on the `MAFFT` alignment of the complete genome sequences (7,681 genomes) resulting from the first filtering steps of `ViralClust`.

### *Alphainfluenzavirus influenzae*

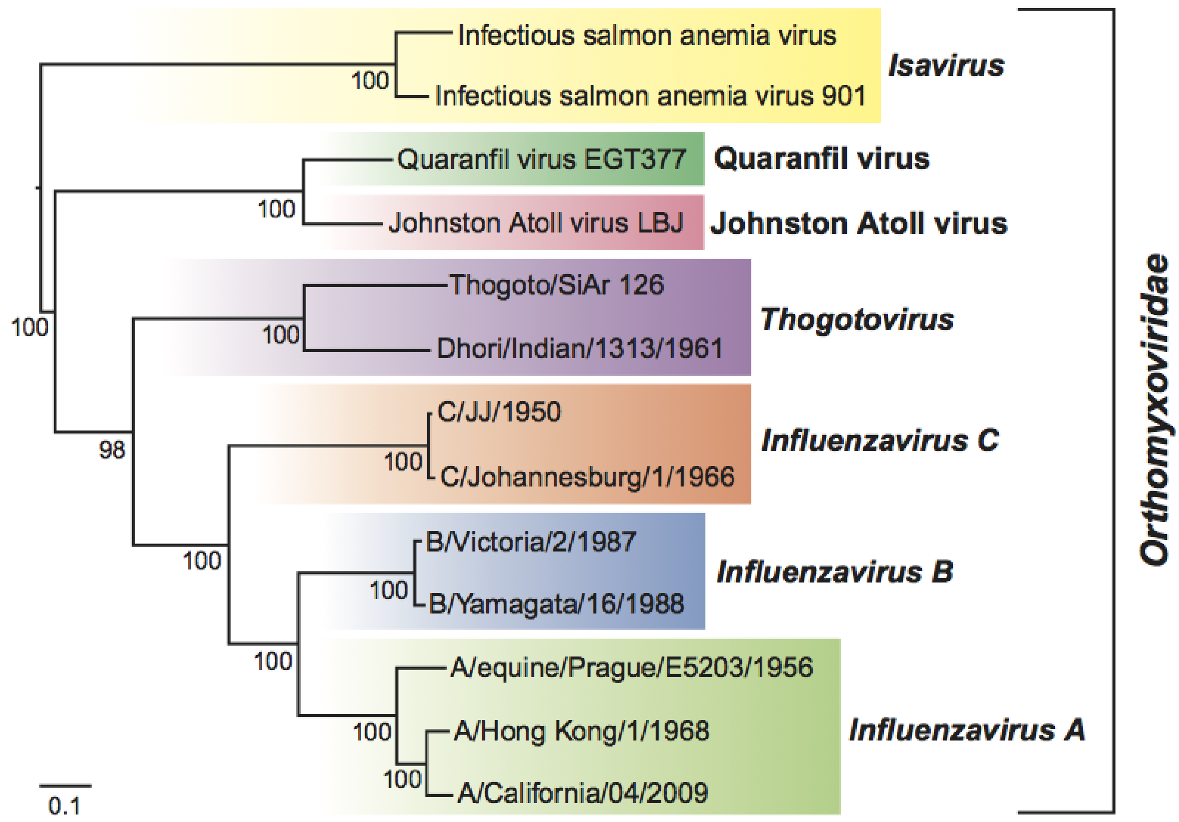


**Figure S10.** Phylogenetic tree of *Orthomyxoviridae* (ICTV) including IAV.

**Table S6.** Clustering IAV with and without (truncated) reverse-complement sequences. Dataset - dataset used as input; rc rep rem – reverse-complement representatives were removed from the representative dataset and used as new results; rc rem – reverse-complement sequences were removed from the non-redundant dataset and used as input for a new clustering run; #Seq – number of input genomes for clustering with `ViralClust`; Number of genomes after clustering – only considering clusters with a size greater than one; cd – `CD-HIT-EST`; sum – `SUMACLUST`; vse – `VSEARCH`; mmseq – `MMSeqs2`; hdb – `HDBSCAN`.

| Segment | Dataset (#Seq) | Number of genomes after clustering | | | | |
|---|---|---|---|---|---|---|
| | | cd | suma | vse | mmseq | hdb |
| 4 (H) | nr (108,514) | 300 | 1,237 | 464 | 1,574 | 2,307 |
| | rc rep rem | 282 | 672 | 289 | 757 | 1,806 |
| | rc rem (87,653) | 933 | 677 | 300 | 728 | 1,767 |
| 6 (N) | nr (73,786) | 228 | 773 | 329 | 640 | 1,622 |
| | rc rep rem | 217 | 430 | 231 | 385 | 1,386 |
| | rc rem (65,393) | 786 | 444 | 233 | 453 | 1,357 |

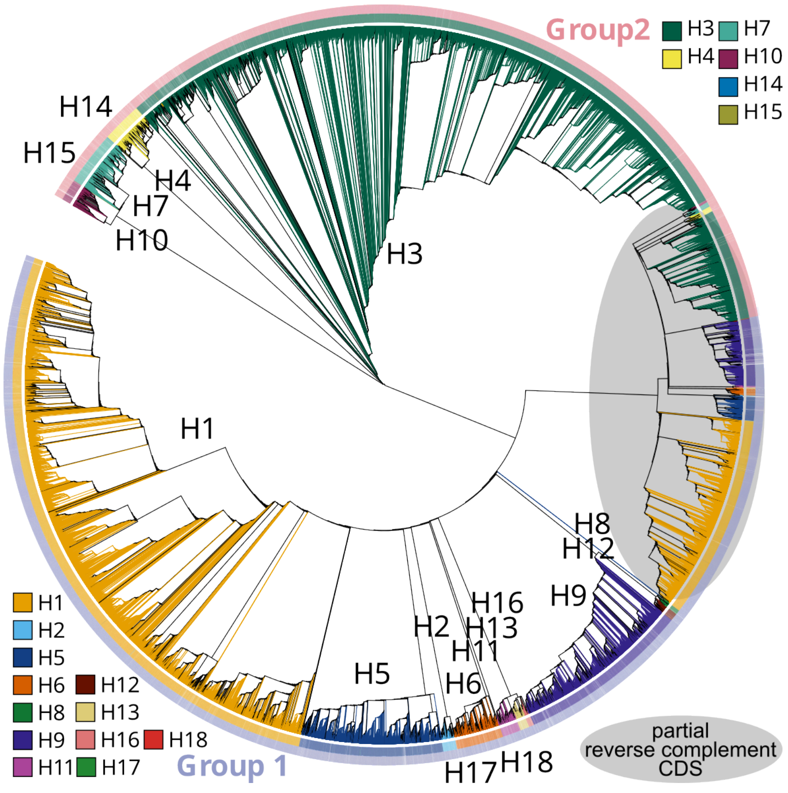


**Figure S11.** Phylogenetic tree of IAV segment 4 (H) with highlighted subtypes H1–H18. Gray area marks genomes with partial reverse complement CDS. That subtree 'mirrors' the rest of the tree. The tree is reconstructed using `FastTree` based on the `MAFFT` alignment of the complete genome sequences (108,514 genomes) resulting from the first filtering steps of `ViralClust`.

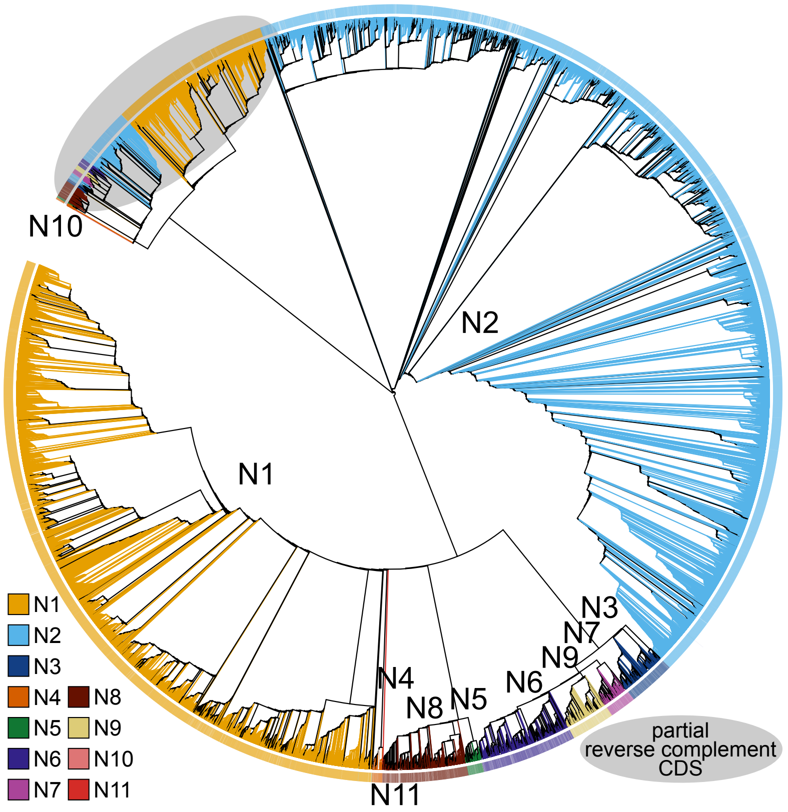


**Figure S12.** Phylogenetic tree of IAV segment 6 (N) with highlighted subtypes N1–N11. Gray area marks genomes with partial reverse complement CDS. That subtree 'mirrors' the rest of the tree. The tree is reconstructed using `FastTree` based on the `MAFFT` alignment of the complete genome sequences (73,786 genomes) resulting from the first filtering steps of `ViralClust`.

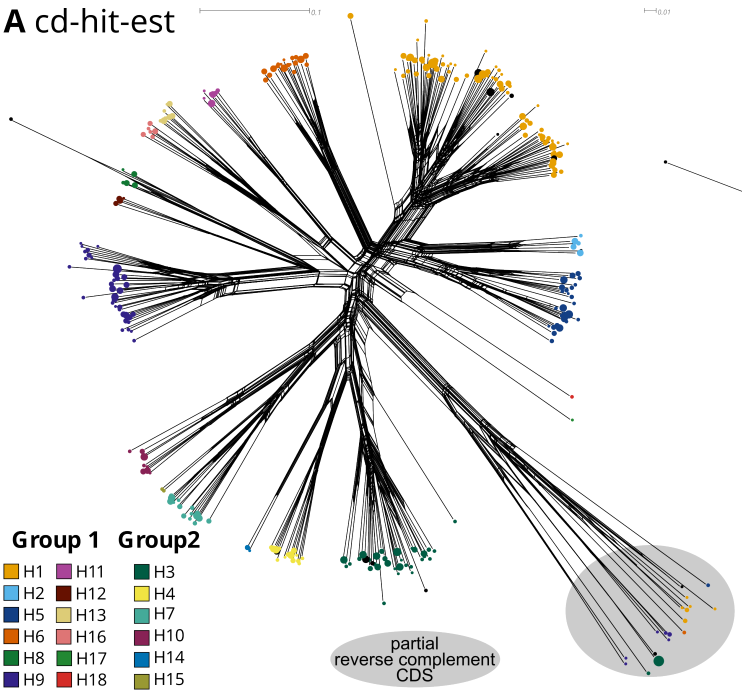


**Figure S13.** Split graphs of IAV segment 4 (H) representatives of (A) CD-HIT-EST. Left – all representatives (including partial CDS); Right – only complete genomes. The node size indicates the cluster size. Subtypes (H1–H18) are color-coded. The split graph is reconstructed using SplitsTree based on the MAFFT alignment of the complete genome sequences (108,514 genomes) resulting from the first filtering steps of ViralClust.

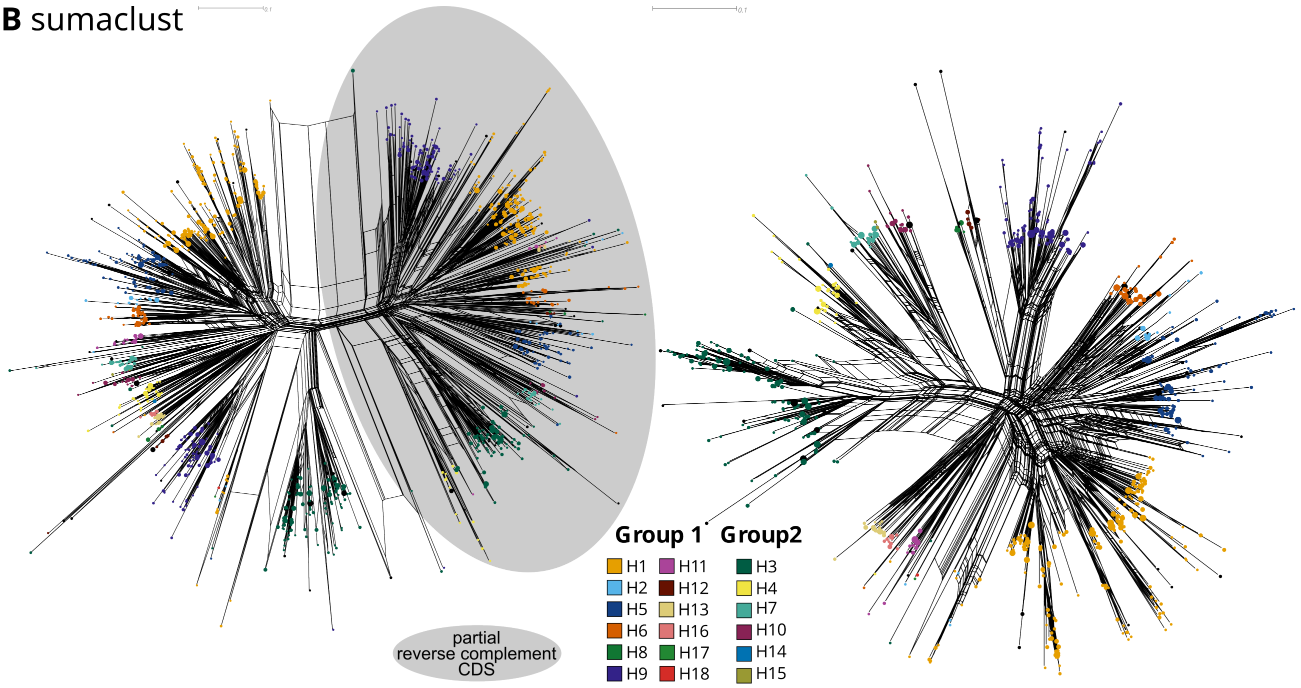


**Figure S14.** Split graphs of IAV segment 4 (H) representatives of (B) SUMACLUST. Left – all representatives (including partial CDS); Right – only complete genomes. The node size indicates the cluster size. Subtypes (H1–H18) are color-coded. The split graph is reconstructed using SplitsTree based on the MAFFT alignment of the complete genome sequences (108,514 genomes) resulting from the first filtering steps of ViralClust.

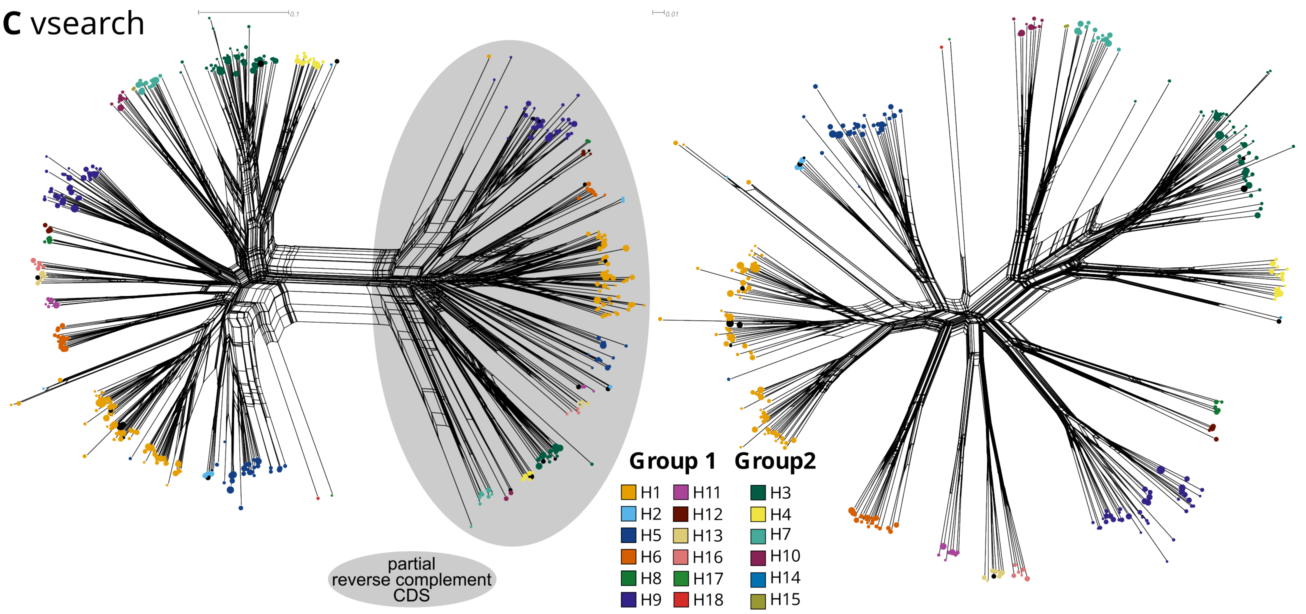


**Figure S15.** Split graphs of IAV segment 4 (H) representatives of (C) `VSEARCH`. Left – all representatives (including partial CDS); Right – only complete genomes. The node size indicates the cluster size. Subtypes (H1–H18) are color-coded. The split graph is reconstructed using `SplitsTree` based on the `MAFFT` alignment of the complete genome sequences (108,514 genomes) resulting from the first filtering steps of `ViralClust`.

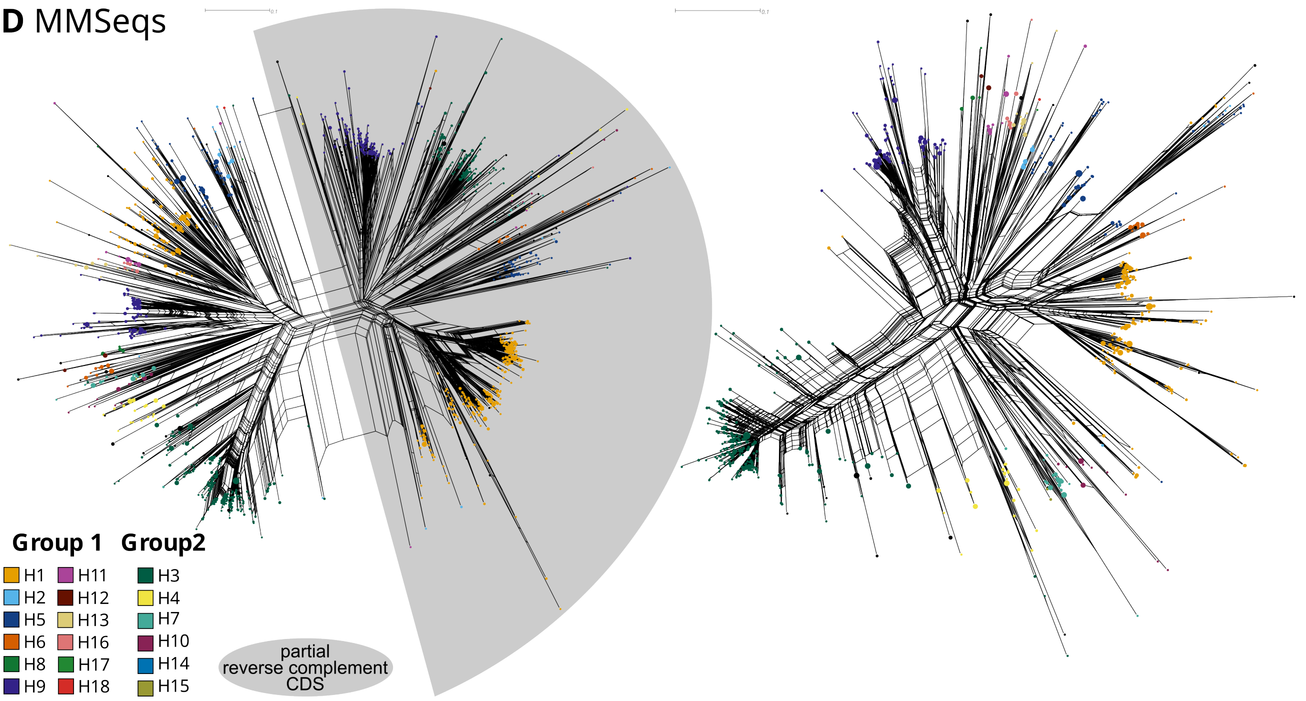


**Figure S16.** Split graphs of IAV segment 4 (H) representatives of (D) `MMSeqs2`. Left – all representatives (including partial CDS); Right – only complete genomes. The node size indicates the cluster size. Subtypes (H1–H18) are color-coded. The split graph is reconstructed using `SplitsTree` based on the `MAFFT` alignment of the complete genome sequences (108,514 genomes) resulting from the first filtering steps of `ViralClust`. `HDBSCAN` did not select any representatives for subtypes H17 and H18. We added these manually to the alignment for split graph reconstruction and marked them as framed circles.

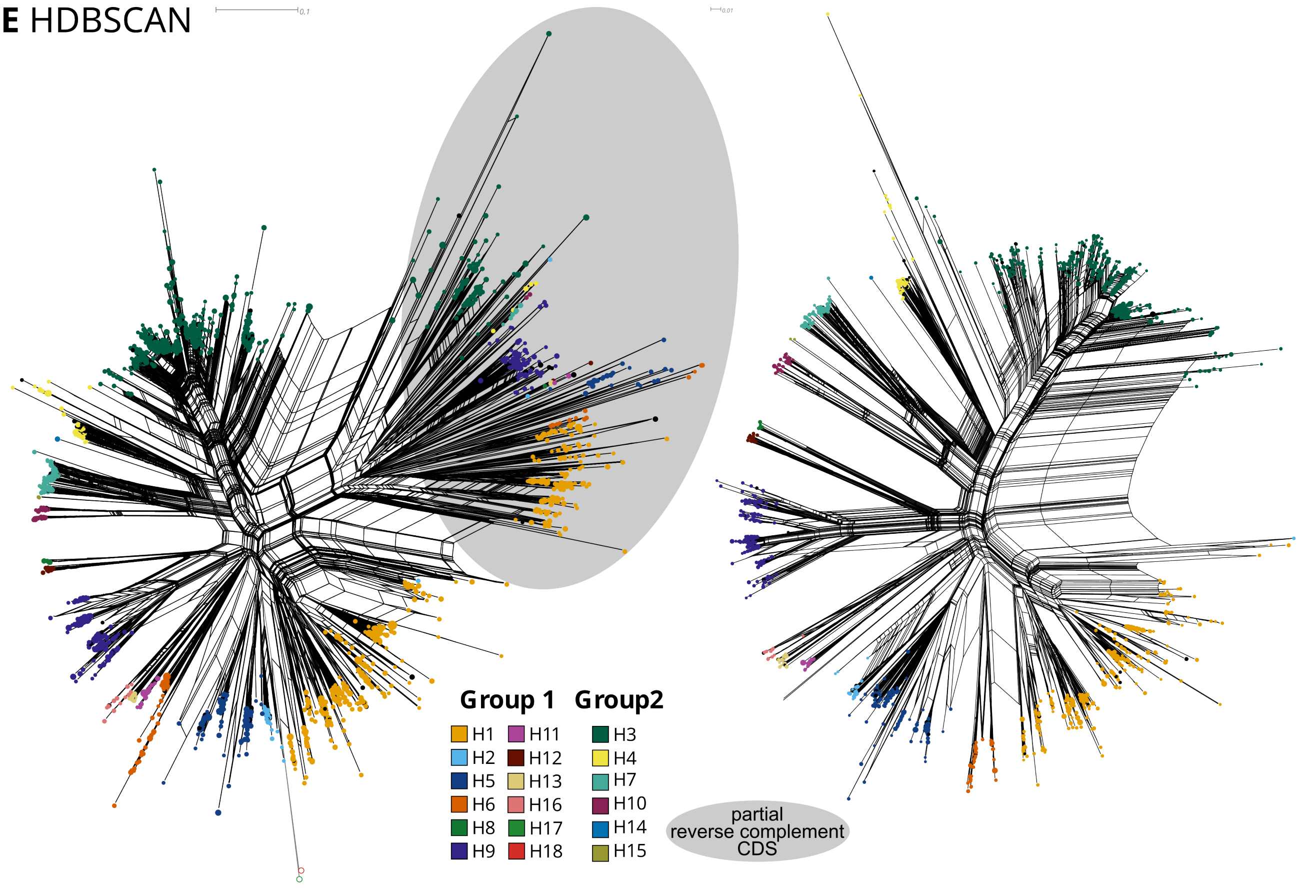


**Figure S17.** Split graphs of IAV segment 4 (H) representatives of (E) HDBSCAN. Left – all representatives (including partial CDS); Right – only complete genomes. The node size indicates the cluster size. Subtypes (H1–H18) are color-coded. The split graph is reconstructed using `SplitsTree` based on the `MAFFT` alignment of the complete genome sequences (108,514 genomes) resulting from the first filtering steps of `ViralClust`.

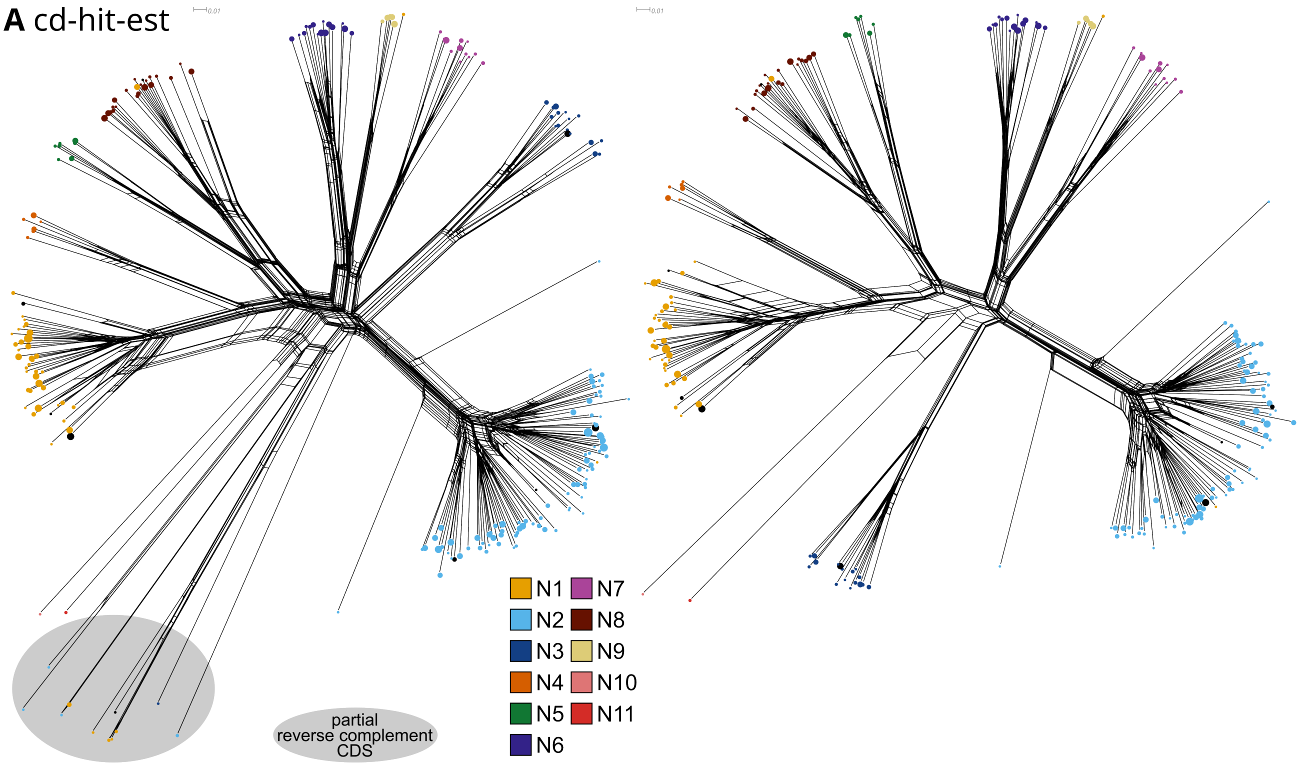


**Figure S18.** Split graphs of IAV segment 6 (N) representatives of (A) CD-HIT-EST. Left – all representatives (including partial CDS); Right – only complete genomes. The node size indicates the cluster size. Subtypes (N1–N11) are color-coded. The split graph is reconstructed using `SplitsTree` based on the `MAFFT` alignment of the complete genome sequences (108,514 genomes) resulting from the first filtering steps of `ViralClust`.

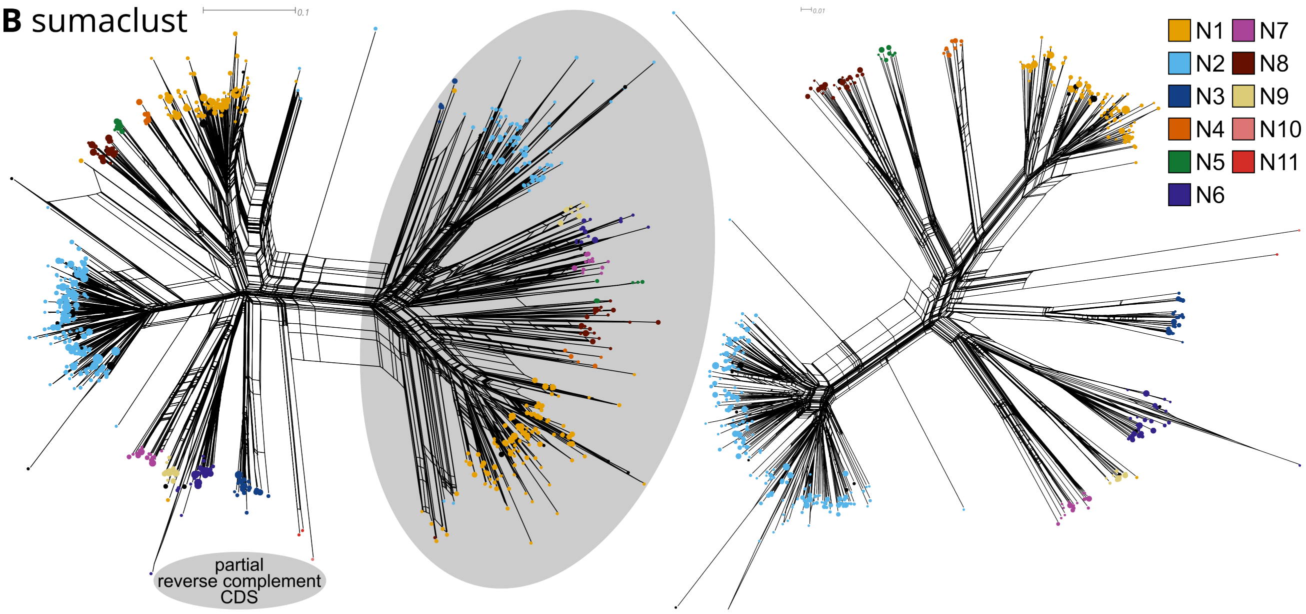


**Figure S19.** Split graphs of IAV segment 6 (N) representatives of (B) SUMACLUST. Left – all representatives (including partial CDS); Right – only complete genomes. The node size indicates the cluster size. Subtypes (N1–N11) are color-coded. The split graph is reconstructed using `SplitsTree` based on the `MAFFT` alignment of the complete genome sequences (108,514 genomes) resulting from the first filtering steps of `ViralClust`.

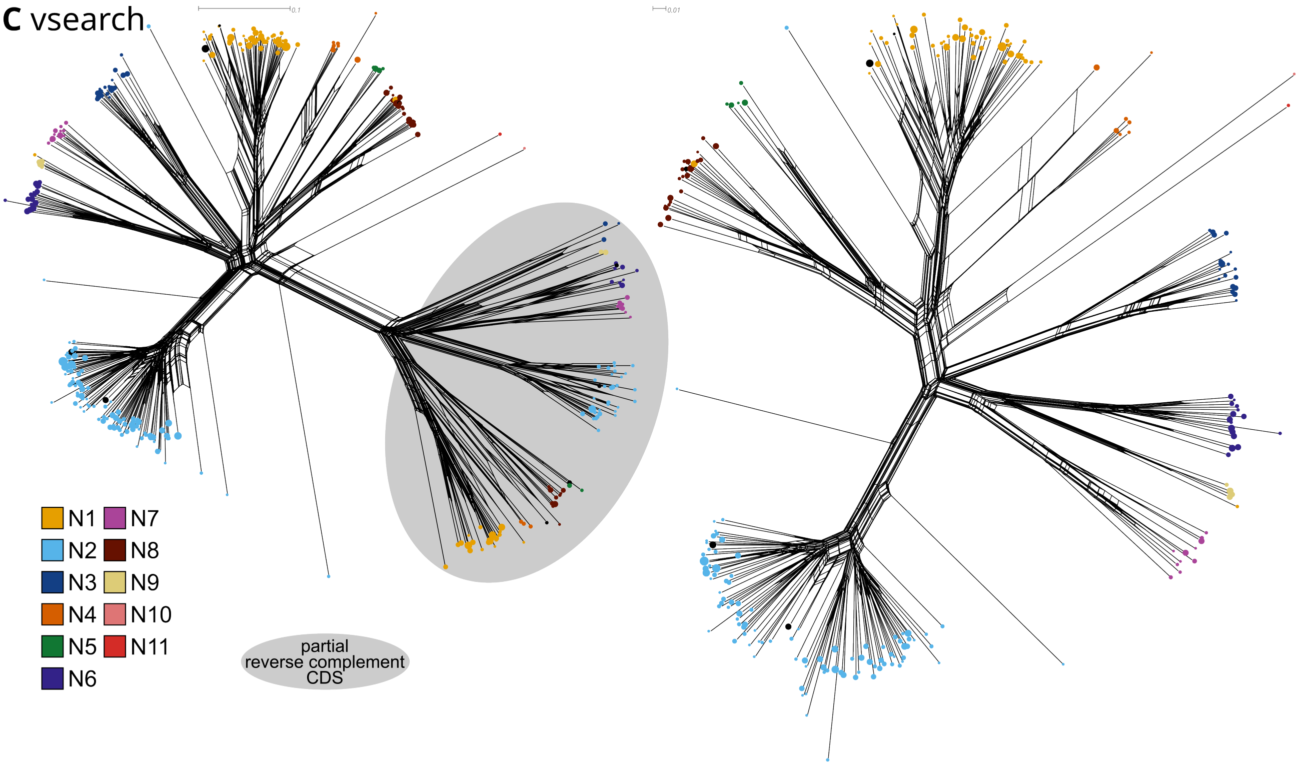


**Figure S20.** Split graphs of IAV segment 6 (N) representatives of (C) VSEARCH. Left – all representatives (including partial CDS); Right – only complete genomes. The node size indicates the cluster size. Subtypes (N1–N11) are color-coded. The split graph is reconstructed using `SplitsTree` based on the `MAFFT` alignment of the complete genome sequences (108,514 genomes) resulting from the first filtering steps of `ViralClust`.

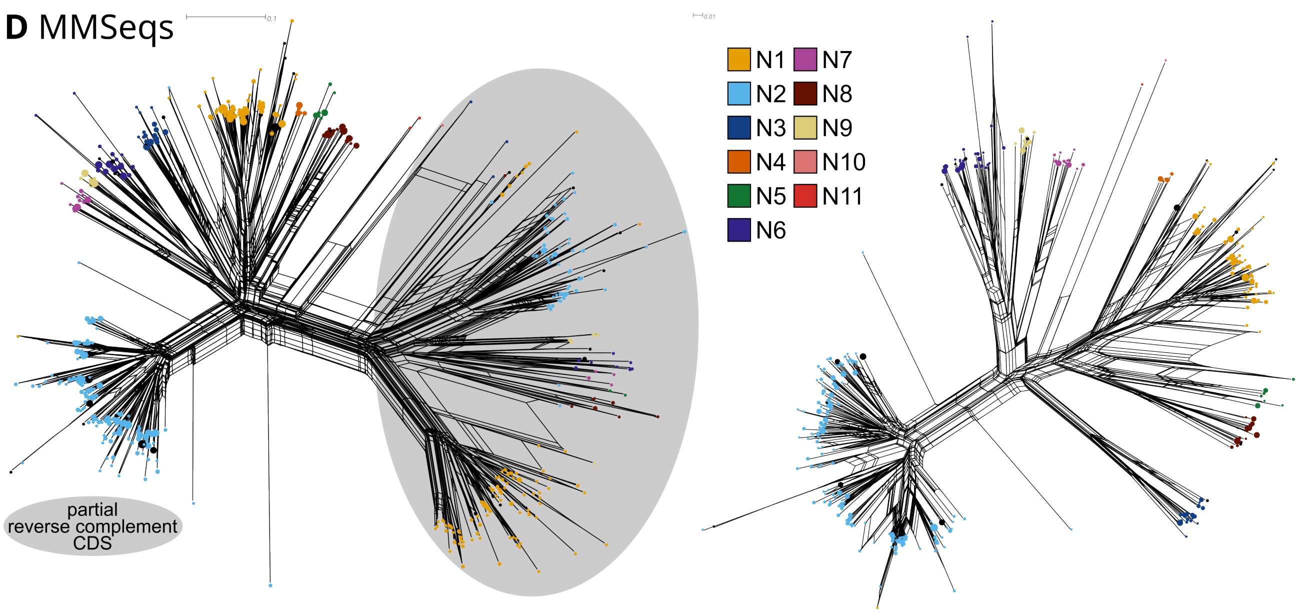


**Figure S21.** Split graphs of IAV segment 6 (N) representatives of (D) `MMSeqs2`. Left – all representatives (including partial CDS); Right – only complete genomes. The node size indicates the cluster size. Subtypes (N1–N11) are color-coded. The split graph is reconstructed using `SplitsTree` based on the `MAFFT` alignment of the complete genome sequences (108,514 genomes) resulting from the first filtering steps of `ViralClust`.

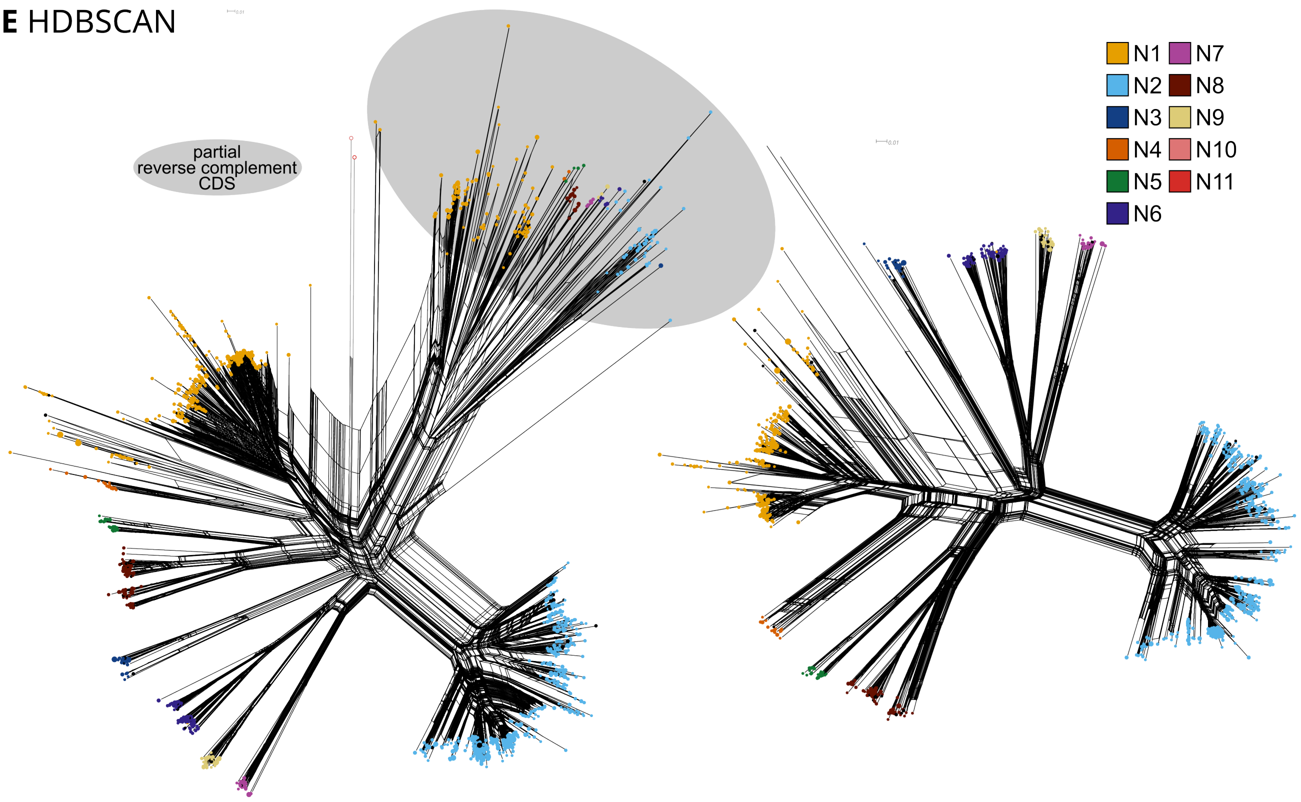


**Figure S22.** Split graphs of IAV segment 6 (N) representatives of (E) `HDBSCAN`. Left – all representatives (including partial CDS); Right – only complete genomes. The node size indicates the cluster size. Subtypes (N1–N11) are color-coded. The split graph is reconstructed using `SplitsTree` based on the `MAFFT` alignment of the complete genome sequences (108,514 genomes) resulting from the first filtering steps of `ViralClust`. `HDBSCAN` did not select any representatives for subtypes N10 and N11. We added these manually to the alignment for split graph reconstruction and marked them as framed circles.

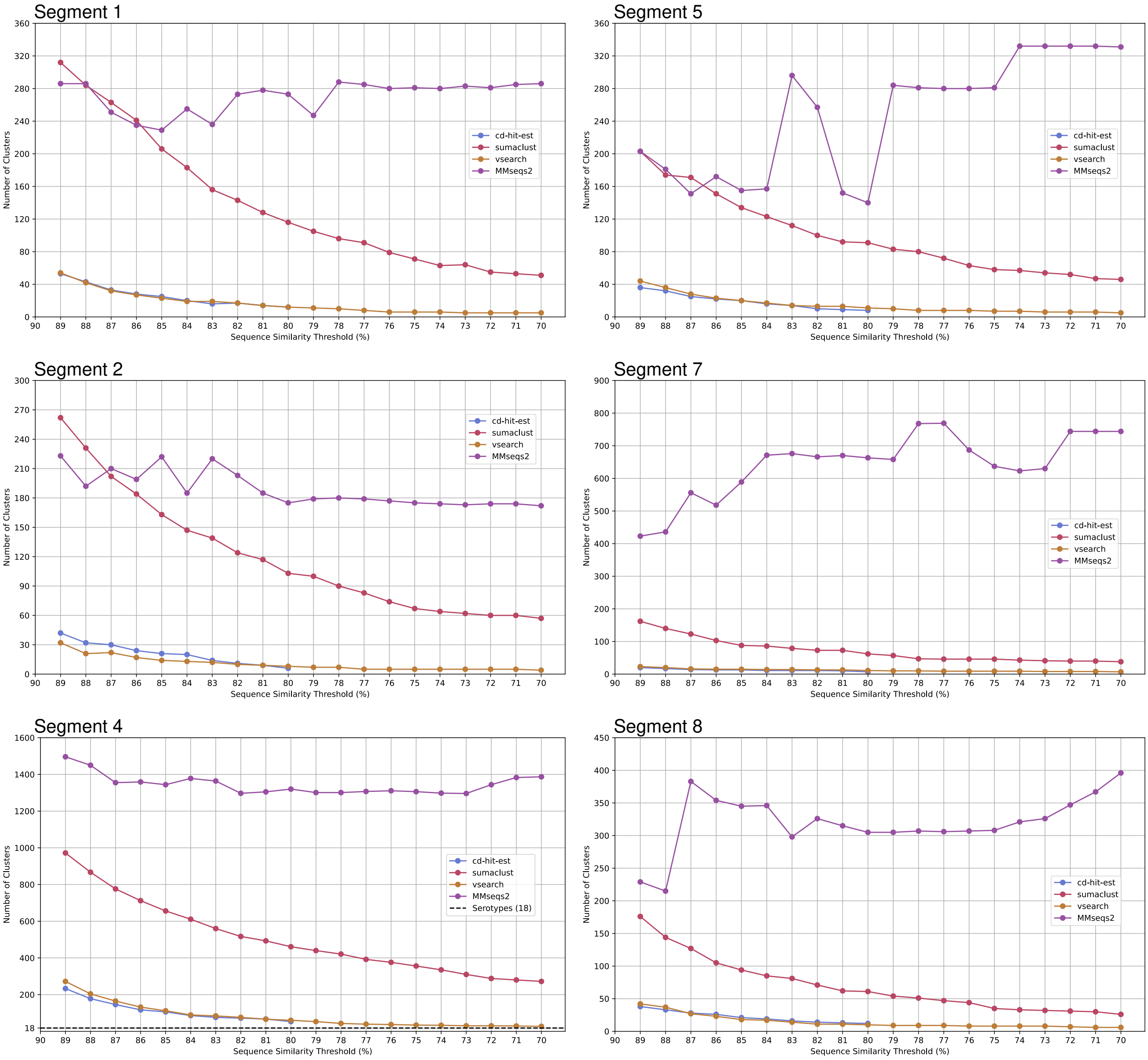


**Figure S23.** Number of clusters of IAV segments 1, 2, 4, 5, 7, and 8 with sequence similarity thresholds from 70 - 90 %. CD-HIT-EST only allows a minimum of 80 %. Only clusters with with a size greater than one are considered.

**Table S7.** Overview of IAV clustering with 70–90 % sequence similarity. Seg – segment; Algo – clustering algorithm; cd – CD-HIT-EST; sum – SUMACLUST; vse – VSEARCH; mmseq – MMSeqs2; Number of genomes after clustering – only considering clusters with a size greater than one.

| Seg | Algo | Sequence similarity threshold (%) | | | | | | | | | | | | | | | | | | | | |
|---|---|---|---|---|---|---|---|---|---|---|---|---|---|---|---|---|---|---|---|---|---|---|
| | | 90 | 89 | 88 | 87 | 86 | 85 | 84 | 83 | 82 | 81 | 80 | 79 | 78 | 77 | 76 | 75 | 74 | 73 | 72 | 71 | 70 |
| 1 | cd | 66 | 53 | 43 | 33 | 28 | 25 | 20 | 16 | 17 | 14 | 12 | | | | | | | | | | |
| | suma | 390 | 312 | 284 | 263 | 241 | 206 | 183 | 156 | 143 | 128 | 116 | 105 | 96 | 91 | 79 | 71 | 63 | 64 | 55 | 53 | 51 |
| | vse | 90 | 54 | 42 | 32 | 27 | 23 | 19 | 19 | 17 | 14 | 12 | 11 | 10 | 8 | 6 | 6 | 6 | 5 | 5 | 5 | 5 |
| | mmseq | 280 | 286 | 286 | 251 | 235 | 229 | 255 | 236 | 273 | 278 | 273 | 247 | 288 | 285 | 280 | 281 | 280 | 283 | 281 | 285 | 286 |
| 2 | cd | 56 | 42 | 32 | 30 | 24 | 21 | 20 | 14 | 11 | 9 | 6 | | | | | | | | | | |
| | suma | 338 | 262 | 231 | 202 | 184 | 163 | 147 | 139 | 124 | 117 | 103 | 100 | 90 | 83 | 74 | 67 | 64 | 62 | 60 | 60 | 57 |
| | vse | 73 | 32 | 21 | 22 | 17 | 14 | 13 | 12 | 10 | 9 | 8 | 7 | 7 | 5 | 5 | 5 | 5 | 5 | 5 | 5 | 4 |
| | mmseq | 216 | 223 | 192 | 210 | 199 | 222 | 185 | 220 | 203 | 185 | 175 | 179 | 180 | 179 | 177 | 175 | 174 | 173 | 174 | 174 | 172 |
| 3 | cd | 63 | 46 | 35 | 31 | 24 | 20 | 18 | 17 | 19 | 15 | 13 | | | | | | | | | | |
| | suma | 305 | 263 | 233 | 200 | 184 | 166 | 153 | 145 | 138 | 119 | 111 | 102 | 94 | 83 | 78 | 72 | 69 | 66 | 60 | 57 | 51 |
| | vse | 75 | 38 | 32 | 25 | 21 | 19 | 17 | 16 | 13 | 13 | 11 | 10 | 8 | 7 | 7 | 7 | 7 | 7 | 7 | 7 | 7 |
| | mmseq | 265 | 193 | 188 | 213 | 208 | 195 | 189 | 551 | 274 | 263 | 298 | 294 | 290 | 308 | 314 | 299 | 325 | 382 | 384 | 384 | 383 |
| 4 (H) | cd | 300 | 233 | 178 | 146 | 117 | 106 | 86 | 76 | 72 | 67 | 53 | | | | | | | | | | |
| | suma | 1,237 | 972 | 867 | 776 | 712 | 656 | 611 | 560 | 517 | 493 | 461 | 440 | 421 | 392 | 376 | 356 | 335 | 310 | 288 | 280 | 272 |
| | vse | 464 | 272 | 204 | 165 | 132 | 112 | 89 | 84 | 76 | 66 | 60 | 53 | 43 | 39 | 37 | 34 | 33 | 30 | 30 | 29 | 27 |
| | mmseq | 1,574 | 1,496 | 1,450 | 1,355 | 1,359 | 1,344 | 1,378 | 1,364 | 1,297 | 1,305 | 1,320 | 1,301 | 1,301 | 1,307 | 1,311 | 1,306 | 1,298 | 1,296 | 1,344 | 1,383 | 1,387 |
| 5 | cd | 46 | 36 | 32 | 25 | 22 | 20 | 16 | 14 | 10 | 9 | 8 | | | | | | | | | | |
| | suma | 262 | 203 | 174 | 171 | 151 | 134 | 123 | 112 | 100 | 92 | 91 | 83 | 80 | 72 | 63 | 58 | 57 | 54 | 52 | 47 | 46 |
| | vse | 76 | 44 | 36 | 28 | 23 | 20 | 17 | 14 | 13 | 13 | 11 | 10 | 8 | 8 | 8 | 7 | 7 | 6 | 6 | 6 | 5 |
| | mmseq | 179 | 203 | 181 | 151 | 172 | 155 | 157 | 296 | 257 | 152 | 140 | 284 | 281 | 280 | 280 | 281 | 332 | 332 | 332 | 332 | 331 |
| 6 (N) | cd | 228 | 190 | 143 | 121 | 94 | 78 | 63 | 63 | 50 | 47 | 43 | | | | | | | | | | |
| | suma | 773 | 618 | 557 | 507 | 457 | 404 | 378 | 357 | 324 | 307 | 298 | 278 | 268 | 244 | 237 | 223 | 206 | 191 | 179 | 173 | 168 |
| | vse | 329 | 215 | 163 | 131 | 104 | 91 | 78 | 69 | 64 | 56 | 48 | 45 | 40 | 40 | 39 | 40 | 32 | 30 | 28 | 24 | 23 |
| | mmseq | 640 | 562 | 541 | 518 | 509 | 484 | 467 | 462 | 530 | 471 | 484 | 554 | 547 | 525 | 533 | 530 | 627 | 636 | 684 | 659 | 704 |
| 7 | cd | 25 | 20 | 17 | 14 | 13 | 13 | 11 | 11 | 11 | 10 | 7 | | | | | | | | | | |
| | suma | 185 | 162 | 140 | 123 | 103 | 88 | 86 | 79 | 73 | 73 | 62 | 57 | 47 | 46 | 46 | 46 | 43 | 41 | 40 | 40 | 38 |
| | vse | 38 | 23 | 20 | 16 | 15 | 15 | 14 | 14 | 13 | 13 | 11 | 10 | 10 | 9 | 9 | 9 | 9 | 8 | 8 | 8 | 7 |
| | mmseq | 981 | 423 | 436 | 556 | 518 | 589 | 671 | 676 | 666 | 670 | 663 | 658 | 768 | 769 | 687 | 637 | 623 | 630 | 744 | 744 | 744 |
| 8 | cd | 51 | 38 | 33 | 28 | 26 | 21 | 19 | 16 | 14 | 13 | 12 | | | | | | | | | | |
| | suma | 185 | 176 | 144 | 127 | 105 | 94 | 85 | 81 | 71 | 62 | 61 | 54 | 51 | 47 | 44 | 35 | 33 | 32 | 31 | 30 | 26 |
| | vse | 38 | 42 | 37 | 27 | 23 | 18 | 17 | 14 | 11 | 11 | 10 | 9 | 9 | 9 | 8 | 8 | 8 | 8 | 7 | 6 | 6 |
| | mmseq | 802 | 229 | 215 | 383 | 354 | 345 | 346 | 298 | 326 | 315 | 305 | 305 | 307 | 306 | 307 | 308 | 321 | 326 | 347 | 367 | 396 |

**Table S8.** Overview of IAV clustering with `HDBSCAN` and k-mer length 1–7. Seg – segment; Number of genomes after clustering – only considering clusters with a size greater than one.

| Seg | k-mer | | | | | | |
|---|---|---|---|---|---|---|---|
| | 7 | 6 | 5 | 4 | 3 | 2 | 1 |
| 1 | 1,412 | 1,426 | 1,130 | 1,158 | 687 | 1,267 | 3,883 |
| 2 | 1,311 | 1,373 | 1,286 | 1,034 | 720 | 1,346 | 3,789 |
| 3 | 1,371 | 1,298 | 1,263 | 1,148 | 798 | 1,168 | 3,968 |
| 4 (H) | 2,307 | 2,333 | 2,298 | 1,854 | 1,560 | 1,466 | 7,103 |
| 5 | 1,218 | 1,082 | 1,006 | 739 | 563 | 3,157 | 2 |
| 6 (N) | 1,622 | 1,351 | 1,369 | 1,167 | 929 | 1,528 | 4,880 |
| 7 | 981 | 948 | 854 | 744 | 354 | 588 | 3,084 |
| 8 | 802 | 590 | 548 | 483 | 295 | 923 | 2 |

#### *Monkeypox virus*

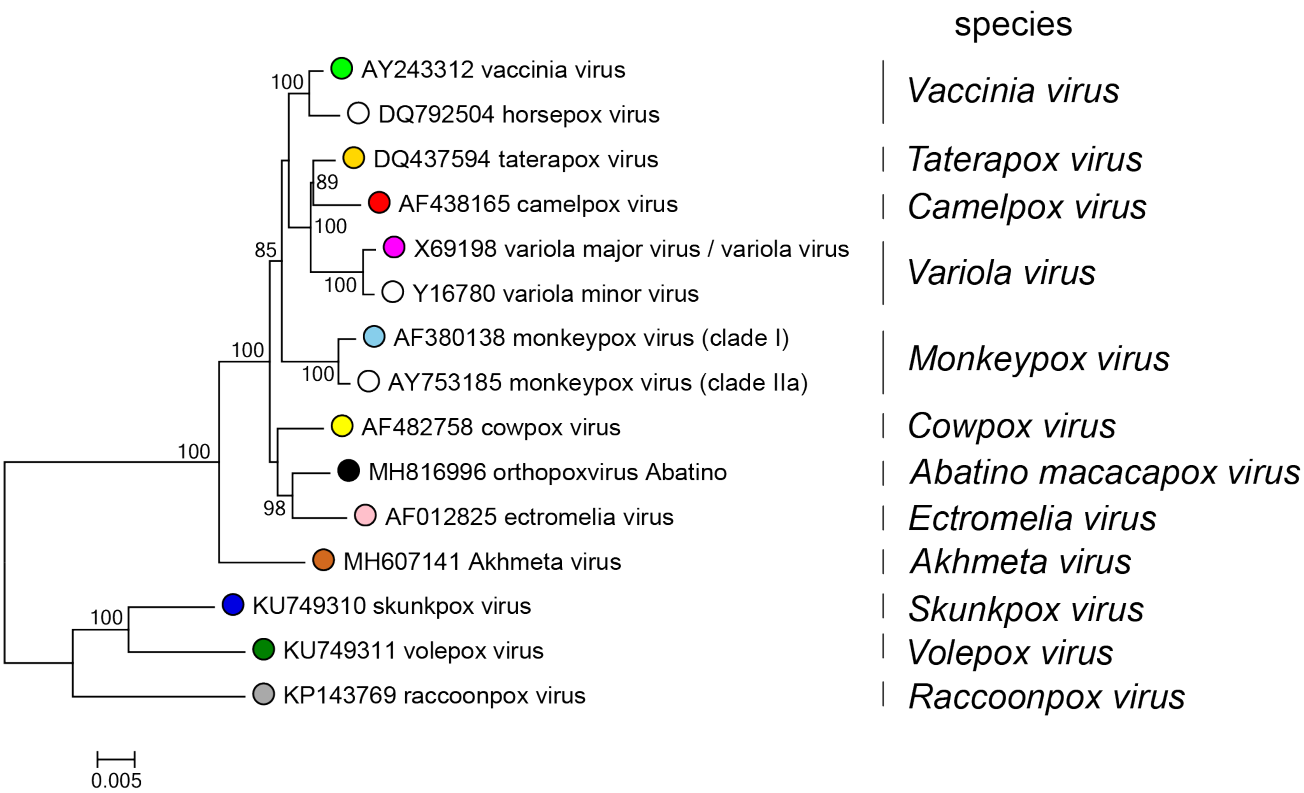


**Figure S24.** Phylogenetic tree of *Orthopoxvirus* (ICTV) including *Monkeypox virus*.
